# A convergent proline biosynthetic step couples cell wall integrity to virulence in Cryptococcus neoformans

**DOI:** 10.64898/2026.10.23.761337

**Authors:** Yujin Lee, Minjae Lee, Seun Kang, Hyunjin Cha, Doyeon Won, Kyung-Ah Lee, Won-Jae Lee, Kyung-Tae Lee, Yong-Sun Bahn

## Abstract

Metabolic adaptation is essential for the pathogenicity of the opportunistic fungus *Cryptococcus neoformans*. Although amino acid biosynthesis pathways are critical fitness determinants in fungal pathogens, the physiological contributions of proline biosynthesis remain incompletely understood, with most studies focusing on proline catabolism. Here, we characterized the proline biosynthetic enzymes Pro1, Pro2, and Pro3 in *C. neoformans*. Proline is produced by two routes that converge on Δ^1^-pyrroline-5-carboxylate (P5C): a de novo pathway from glutamate via Pro1 and Pro2, and an arginine-derived bypass via Car1 and Car2—both requiring Pro3 for the final reduction of P5C to proline. Although all biosynthetic mutants were strictly auxotrophic for proline, their requirements for virulence differed markedly: in a murine infection model, *pro1*Δ and *pro2*Δ mutants retained full virulence, whereas *pro3*Δ exhibited significantly attenuated virulence accompanied by growth defects and impaired cell wall integrity. Genetic disruption of P5C-generating routes partially restored distinct *pro3*Δ phenotypes, with *PRO2* deletion alleviating delayed growth and combined *PRO2* and *CAR2* deletion partially restoring cell size, while cell wall defects remained unresolved. This dissociation indicates that growth requires proper balance of P5C-generating metabolic flux, whereas cell wall integrity depends on a distinct mechanism that is not resolved by blocking P5C production. Together, these findings identify Pro3 as a metabolic gatekeeper coupling proline biosynthesis to cell wall homeostasis, establishing the biosynthetic branch of proline metabolism as a previously unappreciated determinant of fungal pathobiology.

**IMPORTANCE:** *Cryptococcus neoformans* is a leading cause of fungal meningitis worldwide, causing life-threatening infections primarily in immunocompromised individuals. To survive within the human host, this pathogen must synthesize essential nutrients through de novo biosynthesis when host-derived supplies are limited. Although proline catabolism has been studied extensively in fungal pathogens, the physiological roles of individual proline biosynthetic enzymes in human fungal pathogens remain incompletely understood. Here, we show that the terminal enzyme Pro3, but not the upstream enzymes Pro1 and Pro2, is selectively required for full virulence in a murine model. Pro3 supports pathobiological fitness by promoting growth under host-relevant conditions and maintaining cell wall integrity. These findings highlight the importance of proline biosynthesis beyond nutrient production and link this pathway to growth, cell wall integrity, and virulence in *C. neoformans*.

## INTRODUCTION

Pathogenic fungi have evolved sophisticated strategies to adapt to hostile host environments during infection. These include mechanisms that confer resistance to oxidative and osmotic stress, enable survival under nutrient limitation, and allow tolerance of host body temperature (1, 2). Among the metabolic networks supporting these adaptations, amino acid biosynthesis pathways have emerged as critical regulators of fungal physiology and virulence (3, 4). Beyond their canonical role in protein synthesis, these pathways govern metabolic flexibility and stress resistance—traits indispensable for fungal persistence within the host.

Proline, in particular, plays a central role in fungal stress adaptation. In addition to its primary role as a proteinogenic amino acid, proline contributes to stress tolerance through multiple mechanisms: it acts as a compatible solute that protects cells against osmotic imbalance, helps protect cells against oxidative stress, and can serve as an energy source through mitochondrial catabolism (5, 6). Intracellular proline levels are maintained through a dynamic network of biosynthesis, uptake, and catabolism. In the model yeast *Saccharomyces cerevisiae*, proline biosynthesis occurs primarily via the glutamate pathway, a well-characterized three-step enzymatic cascade. First, Pro1, a γ-glutamyl kinase, phosphorylates glutamate to γ-glutamyl phosphate (G5P). G5P is then converted into linear glutamate-γ-semialdehyde (GSA) by Pro2, a γ-glutamyl phosphate reductase. GSA subsequently undergoes spontaneous, non-enzymatic cyclization to form the ring structure Δ^1^-pyrroline-5-carboxylate (P5C). Finally, Pro3, a P5C reductase, catalyzes the reduction of P5C to proline (7). In addition, arginine degradation provides an alternative biosynthetic route to proline. This process involves the hydrolysis of arginine to ornithine by arginase (Car1), followed by conversion of ornithine to GSA by the ornithine aminotransferase (Car2) and subsequent spontaneous cyclization of GSA to P5C. This P5C is then directly reduced to proline by Pro3 (8). Both the canonical proline biosynthesis and the arginine bypass pathways occur within the cytosol (6, 9). Furthermore, proline can be catabolized back to glutamate in the mitochondria via the sequential action of proline oxidase (Put1) and P5C dehydrogenase (Put2). This spatial separation—cytosolic biosynthesis and mitochondrial catabolism—links proline metabolism to intracellular redox balance (NAD(P)^+^/NAD(P)H) and ATP production (6, 9). To maintain homeostasis, *S. cerevisiae* also utilizes the high-affinity amino acid permease Put4 and general amino acid permease Gap1 to take up exogenous proline (6, 9).

While the biochemical framework of proline metabolism is well-established in model yeasts, research in fungal pathogens has predominantly focused on the catabolic branch. In *Candida albicans*, proline catabolism is indispensable not only for morphological switching and survival within host immune cells, but also for maintaining cellular redox homeostasis by preventing excessive ROS accumulation associated with impaired mitochondrial respiration (10). By contrast, de novo proline biosynthesis has received comparatively little attention, likely due to the assumption that pathogens can assimilate abundant proline from host environments. Indeed, physiological studies have shown that the mean concentration of free proline in human plasma is approximately 276 μmol/L (11). However, proline availability is likely to vary substantially among host niches, including extracellular tissues, intracellular phagosomes, and the central nervous system (CNS), raising the possibility that de novo biosynthesis may become important under specific nutrient-restricted conditions. Consequently, despite its fundamental role in cellular physiology, the de novo proline biosynthetic pathway has not been systematically characterized in a human fungal pathogen, and whether it contributes to virulence independently of nutrient provision remains unknown.

*Cryptococcus neoformans* is an opportunistic fungal pathogen that causes life-threatening meningoencephalitis, particularly in immunocompromised individuals (12, 13). During infection, this pathogen can survive within alveolar macrophages, disseminate through the bloodstream, and penetrate the blood-brain barrier (BBB) to establish CNS infection (14). Previous studies on *C. neoformans* have demonstrated that proline catabolism, mediated primarily by the proline oxidase Put5 and the P5C dehydrogenase Put2, is required for proline utilization and the maintenance of mitochondrial ROS homeostasis (15). In particular, disruption of *PUT2* causes excessive mitochondrial ROS accumulation, consistent with a futile P5C-proline cycle, and is accompanied by reduced melanin production and avirulence in a murine model (15).

An earlier insertional mutagenesis screen identified an auxotrophic *C. neoformans* strain carrying a T-DNA insertion in *PRO2*, although genetic linkage between the insertion and the phenotype was not established (16). Large-scale mutagenesis studies have implicated Pro1 as a candidate factor associated with cryptococcal brain infection (17). In a dual signature-tagged mutagenesis (STM) screen that assessed both pulmonary and CNS infectivity, the *pro1*Δ mutant showed reduced recovery from brain tissue, suggesting a potential role for proline metabolism in cryptococcal brain infection. Nevertheless, these large-scale genetic screens did not provide a systematic comparison of the physiological and pathogenic contributions of the individual proline biosynthetic enzymes. A previous direct genetic analysis of proline biosynthetic genes in the plant fungal pathogen *Cryphonectria parasitica* showed that deletion of *PRO1*, *PRO2*, or *PRO3* did not impair virulence, whereas deletion of genes encoding Prodh or P5Cdh resulted in hypovirulence (18). Consequently, whether these findings extend to human fungal pathogens—and whether individual biosynthetic enzymes contribute differentially to virulence—remains an open question.

In this study, we provide the first comprehensive characterization of the proline biosynthetic pathway in a human fungal pathogen. We demonstrate that this pathway is required not only for nutrient-dependent growth but also for tolerance to thermal, oxidative, high-CO_2_, and cell wall stresses. Specifically, we found that *pro3*Δ exhibits impaired cell wall integrity, characterized by significantly reduced levels of key structural components— including chitin, chitosan, and β-glucans—even under nutrient-rich conditions. Furthermore, in a murine infection model, *pro3*Δ displays attenuated virulence, whereas *pro1*Δ and *pro2*Δ exhibit virulence levels comparable to the wild-type (WT) strain. Collectively, these findings reveal that de novo proline biosynthesis contributes to fungal pathobiology beyond simple nutrient provision and provides a foundation for investigating amino acid biosynthetic pathways as determinants of virulence in human fungal pathogens.

## RESULTS

### Identification and evolutionary analysis of the proline biosynthetic pathway in *C. neoformans*

Using the well-characterized *S. cerevisiae* pathway as a reference, we predicted the proline metabolic network in *C. neoformans* by BLAST analysis (**Fig. 1A**) and identified the proline biosynthetic enzymes. We located orthologs of *PRO1* (CNAG_05386), *PRO2* (CNAG_04990), and *PRO3* (CNAG_05899). Phylogenetic analysis revealed that CnPro1 (γ-glutamyl kinase) and CnPro2 (γ-glutamyl phosphate reductase) are evolutionarily distinct from their mammalian counterparts, forming a separate cluster with other fungal and bacterial proteins (**Fig. 1B**). Specifically, whereas plants and mammals encode a bifunctional enzyme, Δ^1^-pyrroline-5-carboxylate synthase (P5CS), in which both the kinase and reductase activities reside in a single polypeptide, *C. neoformans* encodes these activities as two discrete enzymes, similar to *S. cerevisiae*.

**Figure 1.**
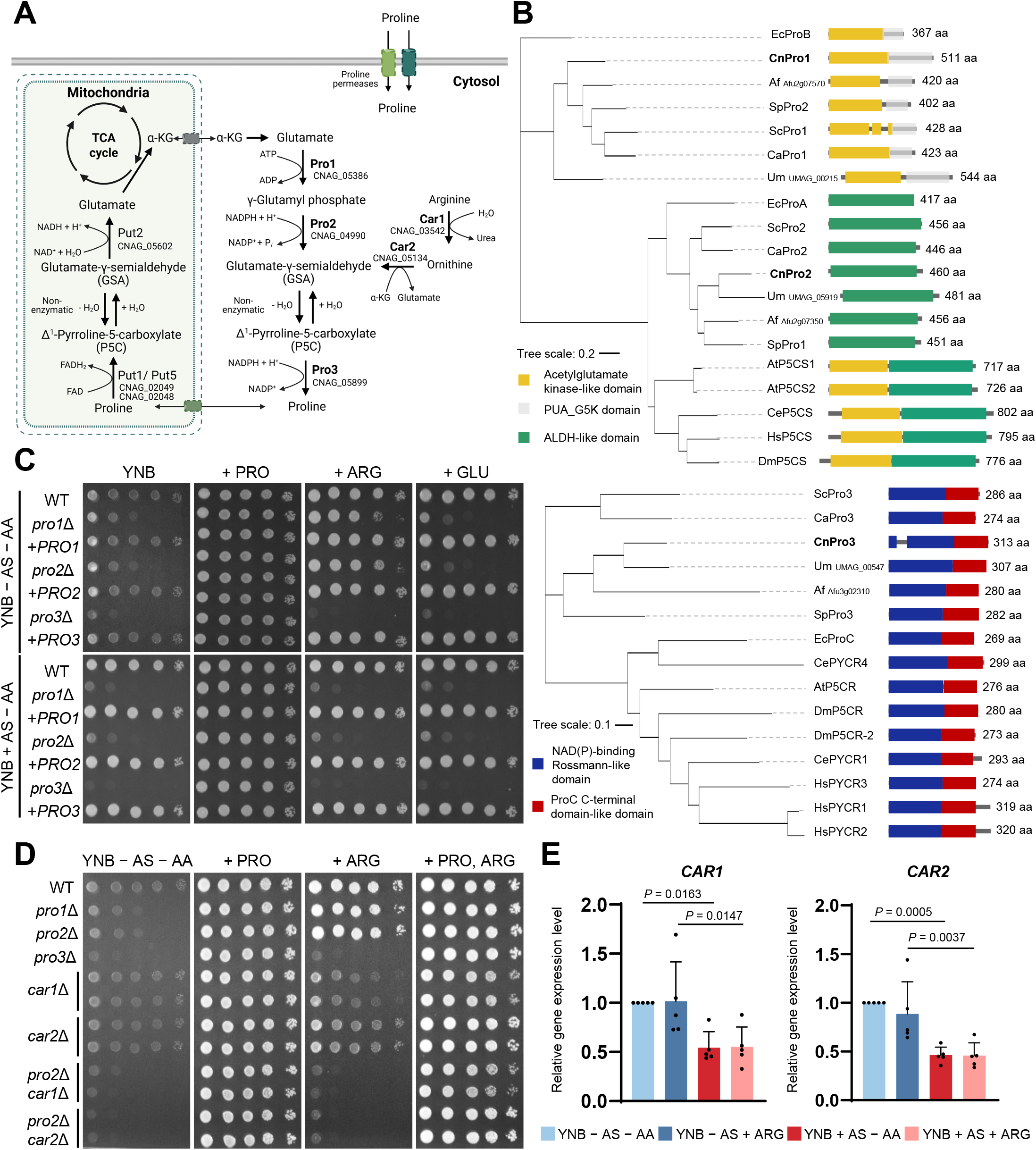
Evolutionary conservation and genetic validation of the proline biosynthetic pathways. (A) Schematic representation of the integrated proline metabolic network in *C. neoformans*. The model illustrates the cytosolic de novo biosynthesis pathway (Pro1-Pro2-Pro3), the alternative arginine-mediated bypass route (Car1-Car2-Pro3), and the mitochondrial catabolic pathway (Put1/Put5 and Put2). The mitochondrial outer and inner membranes are depicted by dashed and solid lines, respectively. The dashed line denotes the comparatively higher permeability of the outer membrane to small solutes. This schematic illustration was created using BioRender (https://BioRender.com/mubyso5). (B) Phylogenetic tree and domain organization of Pro1, Pro2 and Pro3 orthologs. Abbreviations represent the following species: *Cryptococcus neoformans* (Cn), *Ustilago maydis* (Um), *Candida albicans* (Ca), *Saccharomyces cerevisiae* (Sc), *Schizosaccharomyces pombe* (Sp), *Aspergillus fumigatus* (Af), *Escherichia coli* (Ec), *Arabidopsis thaliana* (At), *Caenorhabditis elegans* (Ce), *Drosophila melanogaster* (Dm), and *Homo sapiens* (Hs). Domain information was retrieved from InterPro and UniProt databases. The identified functional domains and their corresponding database identifiers are as follows: the Acetylglutamate kinase-like domain (IPR036393; AGK-like), the PUA_G5K domain (cd21157), the ALDH-like domain (IPR016161), the NAD(P)-binding Rossmann-like domain (G3DSA:3.40.50.720), and the ProC C-terminal domain-like domain (G3DSA:1.10.3730.10). (C) Growth phenotypes of wild type (WT, H99), *pro1*Δ, *pro1*Δ::*PRO1*, *pro2*Δ, *pro2*Δ::*PRO2*, *pro3*Δ, and *pro3*Δ::*PRO3* strains. Strains were grown overnight in YPD at 30°C and then spotted in 10-fold serial dilutions on Yeast Nitrogen Base (YNB) minimal media with (+) or without (-) ammonium sulfate (AS). For nutrient rescue assays, media were individually supplemented with proline (+PRO, 0.2 mg/mL), arginine (+ARG, 0.5 mg/mL), or glutamate (+GLU, 0.5 mg/mL) as indicated. Plates were incubated at 30°C and imaged after 2 days. (D) Growth phenotypes of WT, *pro1*Δ, *pro2*Δ, *pro3*Δ, *car1*Δ, *car2*Δ, *pro2*Δ *car1*Δ, and *pro2*Δ *car2*Δ strains under the indicated nutrient conditions. Growth assays were performed as described in (C), with proline and arginine supplied individually or together as indicated. (E) Transcriptional regulation of the arginine bypass genes, *CAR1* and *CAR2*, by nitrogen availability. WT (H99) cells were subcultured in YPD starting from OD_600_ of 0.2 and harvested upon reaching an OD_600_ of 0.8. Following harvest, the cells were shifted to Yeast Nitrogen Base (YNB) media with (+) or without (-) ammonium sulfate (AS) and/or arginine (ARG, 0.5 mg/mL) for 30 min. Total RNA was extracted, and transcript levels were quantified by qRT-PCR and normalized to *ACT1* expression. Relative expression was calculated using the 2^□ΔΔC^ method, with the YNB condition without AS and amino acid used as the calibrator and assigned a value of 1. Data are presented as mean ± SEM from five independent biological replicates, with individual data points shown. Statistical significance was determined by one-way ANOVA followed by Tukey’s multiple-comparison test separately for each gene.

AlphaFold3-based structural modeling further supported the conservation of these enzymes (**Fig. S1A**). CnPro1 and CnPro2 showed strong spatial similarity to ScPro1 and ScPro2, with sequence similarities of 52.8% and 61.6%, respectively (**Fig. S1B and C**). Despite the structural divergence from mammalian P5CS, individual domains of the cryptococcal enzymes align closely with their human counterparts. The AGK-like domain of CnPro1 and the ALDH-like domain of CnPro2 share 41.4% and 58.0% sequence similarity, respectively, with the corresponding domains in HsP5CS (**Fig. S1D**). These results indicate that the fundamental structural modules required for proline biosynthesis remain highly conserved across eukaryotic species.

CnPro3 also showed conserved folding with ScPro3 (52.8% similarity) and human PYCR3 (44.3% similarity), including the conserved NAD(P)-binding Rossmann-like and ProC C-terminal domains (**Fig. 1B, Fig. S1E to G**). These findings suggest that *C. neoformans* contains a conserved de novo proline biosynthetic pathway.

### The proline biosynthetic pathway is essential for growth under nitrogen-limited conditions

To genetically validate the predicted proline biosynthetic pathway, we used the previously reported *pro1*Δ mutant (19) and constructed *pro2*Δ and *pro3*Δ mutants. Furthermore, we constructed complemented strains for each mutant (*pro1*Δ::*PRO1*, *pro2*Δ::*PRO2*, and *pro3*Δ::*PRO3*) by re-integrating each WT allele into its native locus. Diagnostic PCR and Southern blot analyses confirmed gene disruption in each mutant (**Fig. S2**).

We assessed the physiological requirement of each biosynthetic enzyme through growth assays on nitrogen-limited minimal medium (YNB). All three mutants (*pro1*Δ, *pro2*Δ, and *pro3*Δ) were unable to grow on minimal medium lacking amino acids (AAs), with or without ammonium sulfate (AS) as the nitrogen source, demonstrating complete proline auxotrophy (**Fig. 1C**). Growth was fully restored in all mutants upon the addition of exogenous proline or by gene complementation, confirming that de novo biosynthesis is the primary route for proline prototrophy in *C. neoformans*.

Notably, we observed distinct nutritional requirements for growth recovery when mutants were supplemented with arginine. Arginine partially alleviated the growth defects of *pro1*Δ and *pro2*Δ only under complete nitrogen-limited conditions (YNB-AS−AA), whereas no recovery was observed in the presence of ammonium sulfate (YNB+AS−AA), a preferred nitrogen source (**Fig. 1C**). In contrast, the *pro3*Δ mutant failed to grow on arginine-supplemented medium even in the absence of ammonium sulfate (**Fig. 1A and C**). This result indicates that Pro3 is the obligatory convergence point for both glutamate-derived de novo synthesis and the arginine-mediated bypass route. To confirm that the arginine-mediated recovery of *pro2*Δ is dependent on the predicted bypass enzymes, we further disrupted *CAR1* (CNAG_03542) or *CAR2* (CNAG_05134) in the *pro2*Δ background (*pro2*Δ *car1*Δ and *pro2*Δ *car2*Δ). In growth assays, whereas exogenous arginine partially rescued growth of *pro2*Δ under nitrogen-limited conditions, both *pro2*Δ *car1*Δ and *pro2*Δ *car2*Δ double mutants failed to grow on arginine-supplemented medium, even in the absence of ammonium sulfate (**Fig. 1D**). These results demonstrate that the conversion of arginine to proline requires the sequential actions of Car1 and Car2.

To examine the nitrogen-dependent regulation of this bypass, we analyzed *CAR1* and *CAR2* transcript levels 30 min after shifting mid-log phase WT cells from YPD to YNB medium with or without ammonium sulfate and arginine (**Fig. 1E**). Quantitative reverse transcription-PCR (qRT-PCR) revealed that expression of both *CAR1* and *CAR2* was significantly downregulated in the presence of ammonium sulfate, regardless of arginine supplementation. These results demonstrate that the arginine bypass is regulated by nitrogen source, explaining why exogenous arginine can only compensate for the loss of Pro1 or Pro2 when preferred nitrogen sources are depleted.

### Proline auxotrophy impairs the production of major virulence factors and sexual development

We next addressed whether the proline auxotrophy affects the formation of the antiphagocytic polysaccharide capsule across various inducing conditions (20–22). In DMEM and 10% FBS medium, which are relatively nutrient-rich compared to minimal medium, only the *pro3*Δ mutant exhibited a significant reduction in capsule thickness, whereas *pro1*Δ and *pro2*Δ maintained capsule levels comparable to WT. However, under the more nutrient-restricted conditions of Littman medium, all three biosynthetic mutants showed markedly defective capsule formation (**Fig. 2A**). These results indicate that Pro3 is required for normal capsule formation across the tested media, whereas the requirement for Pro1 and Pro2 is more pronounced under the nutrient-restricted conditions of Littman medium. Exogenous proline fully rescued capsule thickness in all mutants under every condition, confirming these defects reflect intracellular proline availability.

**Figure 2.**
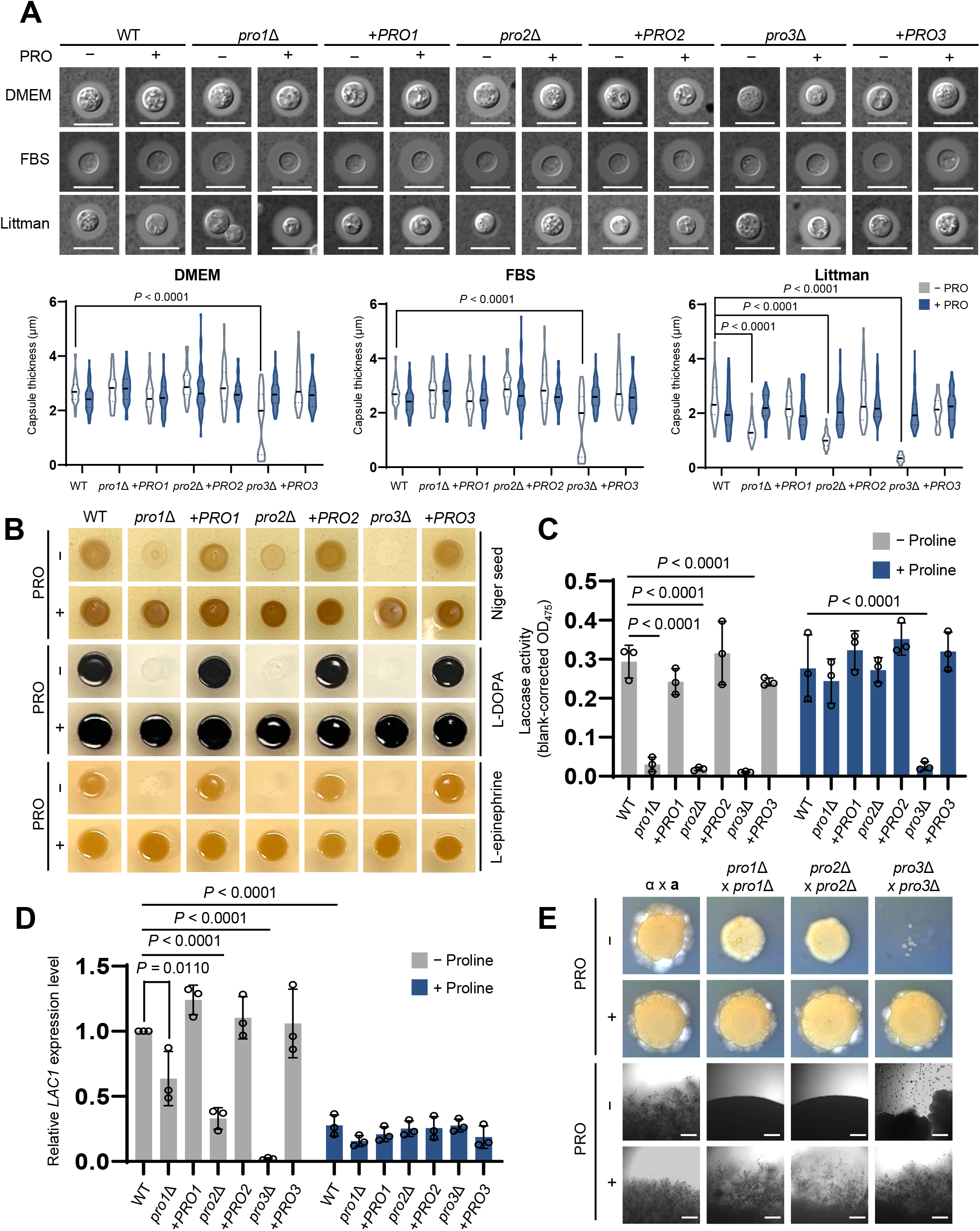
Impact of proline auxotrophy on virulence factor production and sexual development and its rescue by exogenous proline. (A) Capsule thickness of WT and proline biosynthesis mutants. Strains were grown overnight in YPD at 30 °C, washed twice with PBS and spotted on Dulbecco’s modified Eagle medium, 10% FBS medium, or Littman medium with or without proline supplementation (0.1 mg/mL). After 48 h incubation at 37 °C, capsule thickness was measured by India ink staining. Capsule size was quantified from a total of 50 cells per strain. Statistical significance was determined by two-way ANOVA followed by Tukey’s and Šídák’s multiple-comparison tests. Representative images are shown above the quantification. Scale bar, 10 µm. (B) Melanin induction assay of WT and proline biosynthesis mutants. Strains were grown overnight in YPD at 30 °C, washed twice with PBS and spotted on Niger seed, L-DOPA, or L-epinephrine agar containing 0.1% glucose with or without proline supplementation (0.1 mg/mL). Plates were incubated at 37°C for 48 h before imaging. (C) Liquid laccase activity assay. WT (H99), *pro1*Δ, *pro2*Δ, *pro3*Δ, and their complemented strains were incubated in YNB containing ammonium sulfate and 0.1% glucose, with or without L-proline (0.2 mg/mL) for 12 h at 30°C. Following induction, 2 × 10^7^ cells from each culture were collected, washed, and incubated in YNB containing ammonium sulfate, 0.1% glucose, and 10 mM L-DOPA for 10 min at 30°C. Cells were removed by centrifugation, and laccase activity was determined by measuring the absorbance of the supernatant at 475 nm. Values were corrected using cell-free L-DOPA medium incubated under the same conditions as a blank. Data are presented as mean ± SEM from three independent biological replicates, with individual data points shown. Statistical significance was determined by two-way ANOVA followed by Tukey’s multiple-comparison test. (D) *LAC1* expression analysis. WT and mutant strains were grown in YPD to mid-log phase (OD_600_ = 0.6–0.8), washed three times with PBS, and transferred to YNB containing ammonium sulfate without glucose, with or without L-proline (0.2 mg/mL). After incubation for 2 h at 30°C, cells were harvested for RNA extraction and qRT-PCR. *LAC1* transcript levels were normalized to *ACT1* expression, and relative expression was calculated using the 2□^ΔΔC^_t_ method, with the WT sample incubated without proline used as the calibrator and assigned a value of 1. Data are presented as mean ± SEM from three independent biological replicates, with individual data points shown. Statistical significance was determined by two-way ANOVA followed by Tukey’s and Šídák’s multiple-comparison tests. (E) Mating assays were performed by crossing the *MAT*α reference strain H99 with the congenic *MAT***a** strain YL99 (α × **a**), or *MAT*α *pro1*Δ, *pro2*Δ, or *pro3*Δ mutants with the corresponding *MAT***a** deletion mutants. Cells were spotted onto V8 agar medium with or without proline supplementation (0.1 mg/mL) and incubated at 25°C for 14 days before imaging. Representative colony morphologies (top two rows) and filament formation at the colony edges (bottom two rows) are shown. Scale bar, 50 µm.

Next, we investigated the impact of proline auxotrophy on the production of melanin, another critical virulence factor that protects *C. neoformans* from oxidative stress and host immune responses (23, 24). To assess melanization under different conditions, melanin production was examined on three different media: Niger seed, L-DOPA, and L-epinephrine. Under all tested conditions, all three proline biosynthetic mutants displayed markedly reduced melanization compared to the WT (**Fig. 2B**). This defect was fully restored by exogenous proline on solid media. To determine whether the reduced melanization could simply result from differences in cell density, we further assessed laccase activity after normalizing cell numbers following induction. Even after normalization to equal cell numbers, all three biosynthetic mutants showed markedly reduced laccase activity in the absence of proline, whereas their complemented strains exhibited WT-like activity (**Fig. 2C**). Proline supplementation restored laccase activity in the *pro1*Δ and *pro2*Δ mutants, whereas the *pro3*Δ mutant remained defective under the liquid assay conditions. We further examined whether the laccase activity defect was accompanied by altered *LAC1* expression. Under glucose-free induction conditions, *LAC1* transcript levels were markedly reduced in *pro3*Δ without proline but increased following proline supplementation (**Fig. 2D**), despite the persistently low laccase activity observed in the separate activity assay. Because the assays used different induction conditions, these findings do not establish whether the activity defect arises at the transcriptional or post-transcriptional level.

We also evaluated sexual reproduction, a developmental process important for genetic diversity and environmental adaptation (25, 26). Bilateral mating assays were performed on V8 medium by crossing *MAT*α and *MAT***a** strains of each mutant. In contrast to the WT cross, all mutant combinations exhibited severely impaired growth, defective mating, and absence of filament production (**Fig. 2E; Fig. S3**). Notably, the *pro3*Δ mutants exhibited substantially more severe growth inhibition on V8 medium—a relatively nutrient-poor medium— compared to *pro1*Δ and *pro2*Δ mutants, further highlighting the indispensable role of Pro3 as the terminal step of the proline biosynthetic pathway. These developmental defects were fully restored by proline supplementation, with recovery of normal mating and filamentation phenotypes. Collectively, these findings establish that proline biosynthesis is essential for both virulence-factor production and sexual development, with Pro3 serving as the indispensable terminal step, whereas Pro1 and Pro2 are required chiefly under nutrient limitation.

### The proline biosynthetic pathway supports stress adaptation beyond its role in proline provision

To assess stress sensitivity beyond the effects of proline auxotrophy, we first assessed stress sensitivity on YNB minimal medium supplemented with exogenous proline. Notably, all three proline biosynthetic mutants remained sensitive to the oxidative stressor *tert*-butyl hydroperoxide (tBOOH), the cell wall stressor calcofluor white (CFW), and the membrane stressor sodium dodecyl sulfate (SDS). In contrast, sensitivity to hydrogen peroxide (H_2_O_2_), the DNA-damaging agent hydroxyurea (HU), and the antifungal drug amphotericin B (AMB) persisted in the *pro1*Δ and *pro2*Δ mutants but was largely alleviated in the *pro3*Δ mutant (**Fig. 3A**). Sensitivity to tBOOH, the ER stressor dithiothreitol (DTT), CFW, and SDS was also observed in nutrient-rich YPD medium (**Fig. S4B**). Notably, the *pro3*Δ::*PRO3* complemented strain consistently exhibited slightly greater SDS sensitivity than the *pro3*Δ mutant in repeated assays, although the basis for this unexpected phenotype remains unclear. By contrast, sensitivity to most other stressors tested—including the genotoxic agent methyl methanesulfonate (MMS) and antifungal drugs such as fluconazole (FLC) and flucytosine (5-FC)—was fully rescued by proline supplementation, indicating that tolerance to these stressors depends primarily on proline availability (**Fig. S4A and C**).

**Figure 3.**
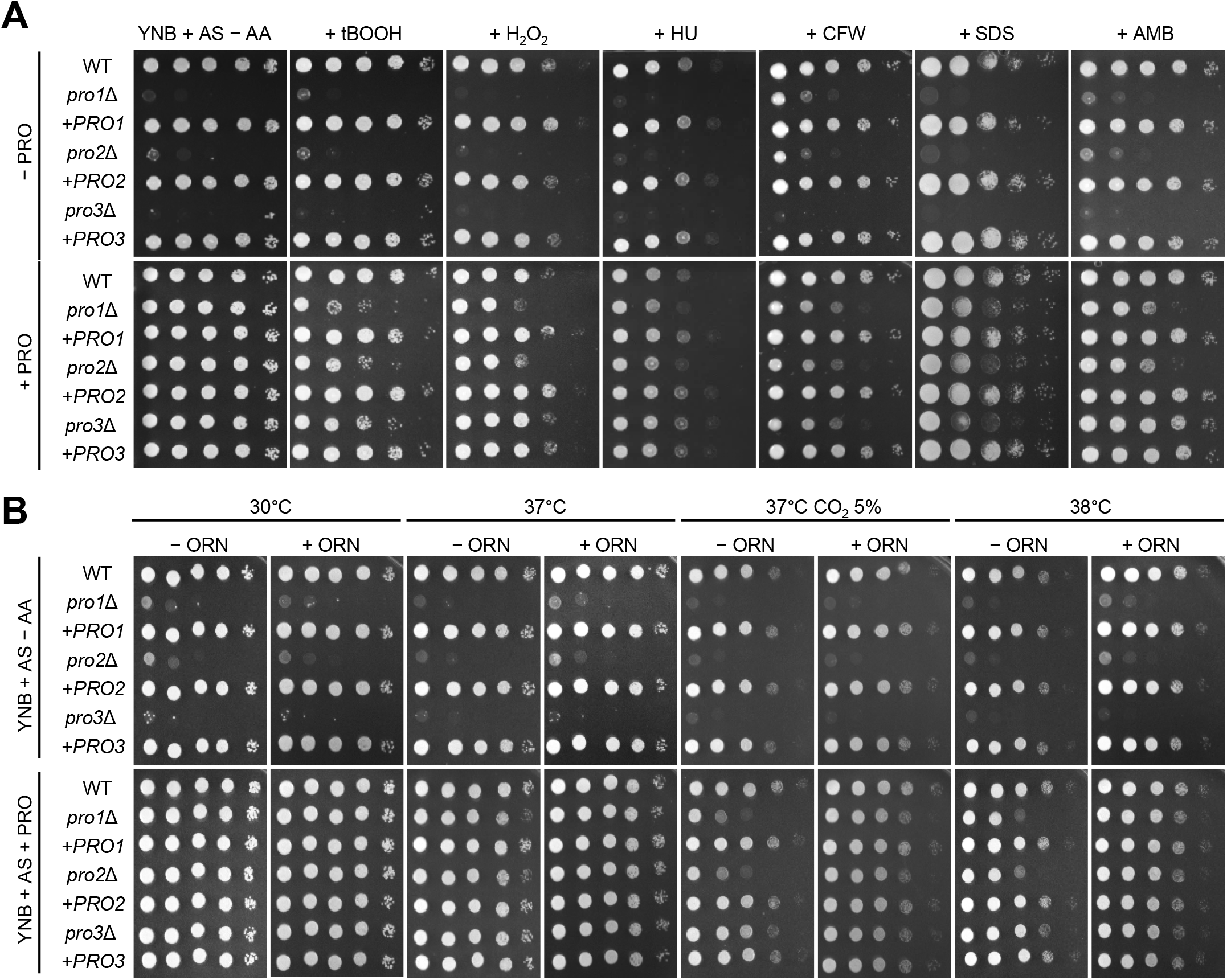
The proline biosynthetic pathway is required for environmental stress response and adaptation independently of proline availability. (A) Stress resistance assay in YNB with ammonium sulfate (AS) media. WT (H99), *pro1*Δ, *pro2*Δ, *pro3*Δ, and their complemented strains were grown overnight in YPD, serially diluted (10-fold), and spotted onto YNB medium containing ammonium sulfate with or without proline (PRO; 0.2 mg/mL). Media contained 0.3 mM *tert*-butyl hydroperoxide (tBOOH), 1.5 mM hydrogen peroxide (H_2_O_2_), 110 mM hydroxyurea (HU), 5 mg/mL CFW, 0.01 % sodium dodecyl sulfate (SDS), or 0.6 μg/mL amphotericin B (AMB). Plates were incubated at 30°C for 2 days and imaged. (B) Thermotolerance and CO_2_ tolerance assay. The same set of strains was grown in YPD, washed, 10-fold serially diluted, and spotted onto YNB + AS medium with or without proline (0.2 mg/mL) or ornithine (ORN; 2 mg/mL). Plates were incubated at 30°C, 37°C, 38°C or 37°C under 5% CO_2_, and imaged after 2 days.

To determine how the proline biosynthetic pathway contributes to host-related stress adaptation, we examined the growth of proline biosynthetic mutants under elevated temperature (37-38°C) and host-mimicking CO_2_ (5% CO_2_) conditions on YNB medium. On YNB medium lacking amino acids, *pro1*Δ and *pro2*Δ mutants showed growth defects under both high-temperature and high-CO_2_ conditions, and these defects persisted when the medium was supplemented with exogenous proline (0.2 mg/mL). In contrast, proline fully restored the growth of the *pro3*Δ mutant (**Fig. 3B**). The high-temperature and high-CO_2_ sensitivities of *pro1*Δ and *pro2*Δ were also observed in YPD medium (**Fig. S4D**). Although proline alone failed to restore the growth of *pro1*Δ and *pro2*Δ under the high-temperature and high-CO_2_ conditions, supplementation with both proline and ornithine partially restored growth (**Fig. 3B**). Ornithine alone did not support growth, consistent with the strict proline auxotrophy of these mutants. Taken together, these results suggest that Pro1 and Pro2 contribute to growth at elevated temperature and high CO_2_ beyond their role in supplying proline.

### Pro3 is required for the full virulence of *C. neoformans*

To assess the contribution of proline biosynthesis to virulence in a mammalian host, we used a murine intranasal infection model. The *pro1*Δ and *pro2*Δ mutants exhibited WT-level virulence, whereas *pro3*Δ mutant resulted in significantly prolonged host survival (*P* = 0.0037) (**Fig. 4A**). This virulence defect was restored in the *pro3*Δ::*PRO3* complemented strain (*P* = 0.0052 vs. *pro3*Δ; *P* = 0.3450 vs. WT), indicating that Pro3 is specifically required for full pathogenicity in this model.

**Figure 4.**
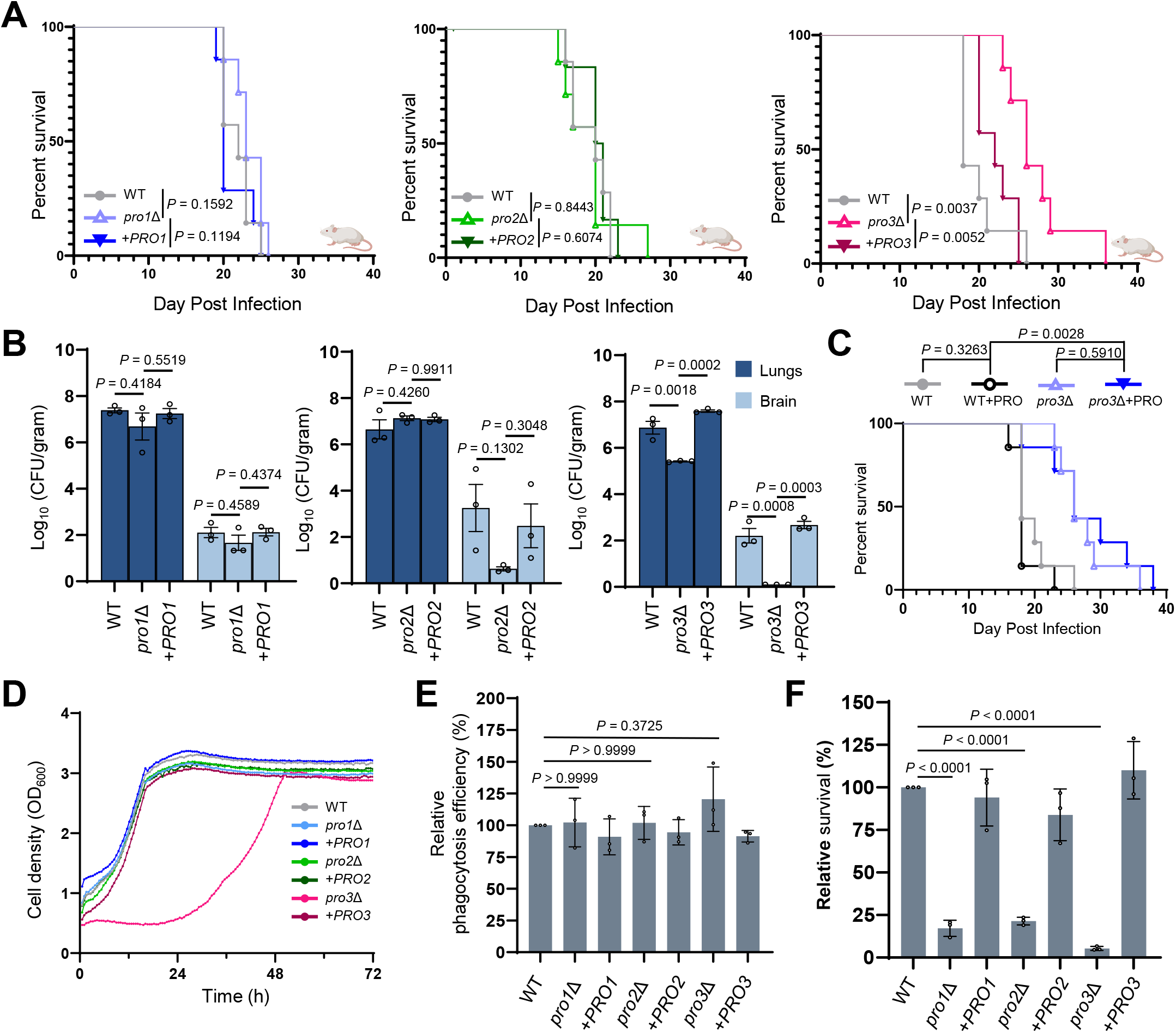
Loss of *PRO3* attenuates virulence and reduces fungal burden in a murine model. (A) Murine survival assay. Mice were intranasally infected with 5 × 10^5^ cells of WT (H99), *pro1*Δ, *pro2*Δ, *pro3*Δ, and their complemented strains (n = 7). Survival was monitored daily. Kaplan–Meier curves were compared using the log-rank (Mantel–Cox) test. Significant differences were observed between WT and *pro3*Δ (*P* = 0.0037). (B) Fungal burden analysis. Lungs and brain fungal loads were quantified at 14 days post-infection and expressed as log10 CFU per gram of tissue (n = 3). Groups were analyzed separately for each organ using one-way ANOVA followed by Tukey’s multiple comparison test. (C) Proline supplementation assay. Mice were infected with WT or *pro3*Δ and maintained with or without continuous proline supplementation in drinking water (0.5% [w/v]; n = 7 per group). Survival was monitored daily and analyzed by the log-rank (Mantel–Cox) test. (D) In vitro growth assay under host-mimicking condition. The growth of *C. neoformans* wild-type (H99), proline auxotrophic mutants (*pro1*Δ, *pro2*Δ, and *pro3*Δ) and their respective complemented strains was monitored in RPMI 1640 medium supplemented with 10% FBS at 37 °C using the RTS-8 bioreactor system. Cell density (OD_600_) was measured every 30 min for 72 hours. The *pro3*Δ mutant exhibited a distinct growth defect compared to the WT. (E-F) Macrophage assays. J774A.1 cells (10^5^ cells/well) were seeded in a 96-well plate and cultured for 16-18 hours and activated with phorbol myristate acetate. *C. neoformans* cells were incubated with 18B7 mAb for 1 hour at 37°C. *C. neoformans* and J774A.1 cells were co-cultured in equal proportion at 37°C with 5% CO_2_ in DMEM media containing 10% FBS. (E) Phagocytosis efficiency. The internalization rate was determined by measuring CFU at 1 h post-infection. After co-incubation, extracellular fungal cells were removed by washing with PBS, and macrophages were lysed to harvest internalized fungal cells. (F) Intracellular survival. The relative survival (%) was determined by measuring CFU at 24 h post-infection. After co-incubation, macrophages were lysed to harvest fungal cells. For panels E and F, data are presented as mean ± SEM from three independent biological replicates (n = 3). Statistical significance was determined separately for each panel by one-way ANOVA followed by Tukey’s multiple-comparison test.

Consistent with the survival data, fungal burden analysis at 14 days post-infection revealed that the *pro3*Δ mutant showed significantly reduced fungal loads in both the lungs and brain compared with the WT strain (**Fig. 4B**). In contrast, the *pro1*Δ and *pro2*Δ mutants exhibited fungal burdens comparable to those of WT-infected mice. To determine whether proline supplementation in drinking water could alleviate the attenuated virulence of *pro3*Δ, we performed infection assays with continuous proline supplementation. However, this treatment did not restore *pro3*Δ virulence (**Fig. 4C**), indicating that proline supplementation under the conditions tested was insufficient to rescue the virulence defect.

We also assessed virulence in *Drosophila melanogaster* as a non-mammalian infection model (27). Unlike in mice, *Drosophila* survival assays did not show a significant reduction in *pro3*Δ virulence (**Fig. S5**). This difference may partly reflect the distinct host physiology and thermal environment of the two infection models. Flies were routinely reared at 25°C, but following infection they were maintained at 30°C to support robust fungal proliferation during the assay period; this temperature, while elevated relative to standard rearing conditions, remains substantially lower than the ∼37°C mammalian host environment encountered by *C. neoformans* during murine infection. Thus, the pathogenic contribution of Pro3 appears to be more evident under mammalian host conditions.

To further define the cellular basis of this attenuation, we examined the growth of the mutants under mammalian tissue culture conditions (RPMI 1640 medium with 10% FBS at 37°C). The *pro1*Δ and *pro2*Δ mutants grew comparably to the WT, whereas the *pro3*Δ mutant exhibited a markedly prolonged lag phase and delayed entry into stationary phase, although it ultimately reached a comparable final cell density (**Fig. 4D**). The growth profile parallels the in vivo attenuation of *pro3*Δ and suggests that Pro3 is uniquely required for efficient proliferation in host-like conditions.

Finally, using J774A.1 macrophages, we found that phagocytosis efficiency was comparable across all strains (**Fig. 4E**), whereas intracellular survival was significantly impaired in all three mutants relative to the WT (**Fig. 4F**). The macrophage survival defect of *pro3*Δ paralleled its in vivo attenuation, whereas the defects of *pro1*Δ and *pro2*Δ did not translate into reduced murine virulence.

### Pro3 is required for maintaining normal cell morphology and cell wall integrity

Given the hypersensitivity of proline biosynthetic mutants to CFW, we examined cell morphology and wall architecture in proline-replete YPD medium to exclude defects arising from proline auxotrophy. We first monitored cell size distribution over 72 h after subculture to mid-exponential phase (OD_600_ = 0.8), defined as 0 h. At 0 h, the cell diameters of all proline biosynthetic mutants were comparable to those of the WT. After 24 h, the WT, *pro1*Δ, and *pro2*Δ strains exhibited an increase in cell size, which was sustained at 48 and 72 h (**Fig. 5A; Fig. S6**). In contrast, from 24 h onward, the *pro3*Δ mutant failed to reach the cell size observed in WT cells and remained significantly smaller throughout the 72-h incubation period (**Fig. 5A**).

**Figure 5.**
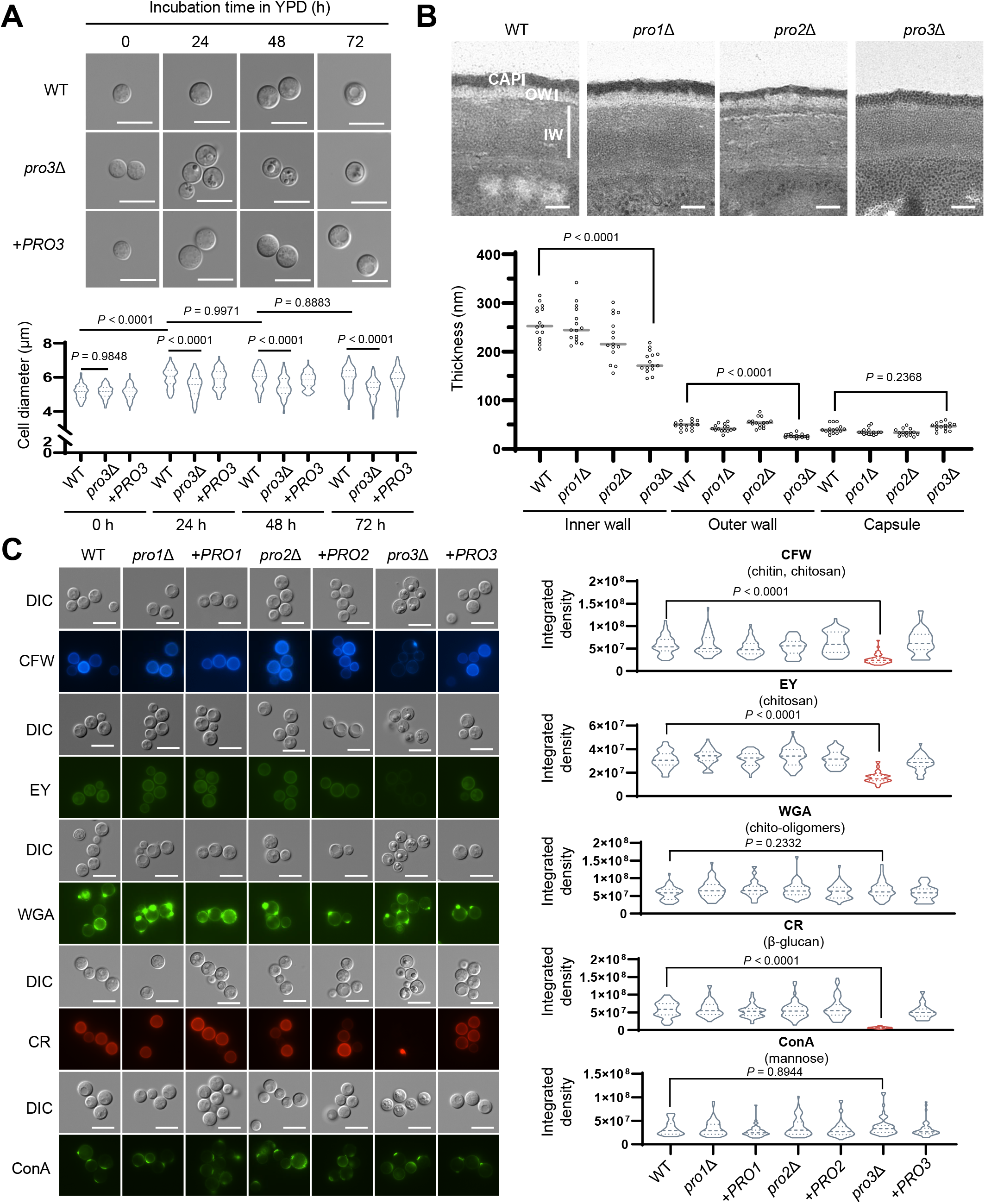
Pro3 is essential for maintaining normal cell morphology and cell wall integrity. **(A)** Cells from an overnight culture were sub-cultured into fresh YPD medium at an initial OD_600_ of 0.2. Once the culture reached the mid-exponential phase (OD_600_ = 0.8), this point was defined as 0 h and the cells were further incubated for 72 h with imaging performed at 24 h intervals. Violin plots show the distribution of cell diameters (n = 100). Scale bar represents 10 µm. Statistical significance was determined by two-way ANOVA followed by Tukey’s multiple-comparison test. (B) Ultrastructural analysis. Representative TEM images of WT (H99), *pro1*Δ, *pro2*Δ, *pro3*Δ cells cultured in YPD for 48 h are shown, along with quantification of the inner wall (IW), outer wall (OW), and capsule (CAP) thickness measured using ImageJ (n = 15 cells per strain). Scale bar represents 100 nm. Statistical significance was determined by one-way ANOVA followed by Tukey’s multiple-comparison test. (C) Cell wall staining. Representative DIC and fluorescence images of cells stained with Calcofluor White (CFW), Eosin Y (EY), Wheat Germ Agglutinin (WGA), Congo Red (CR), and Concanavalin A (ConA) after 48 h of culture in YPD. Fluorescence was quantified as integrated density within individual cell outlines using ImageJ for 50 randomly selected cells per strain. Scale bar represents 10 µm. Statistical significance was determined by one-way ANOVA followed by Tukey’s multiple-comparison test.

To further characterize these morphological abnormalities at higher resolution, we examined cell wall structure by transmission electron microscopy (TEM). The cryptococcal cell envelope consists of a cell wall surrounded by a polysaccharide capsule, and TEM analysis allowed us to distinguish ultrastructural inner and outer wall regions for thickness measurements. In the *pro3*Δ mutant, both the inner and outer wall layers were significantly thinner than in the WT, whereas capsule thickness was comparable across all strains (**Fig. 5B**). In contrast, the *pro1*Δ and *pro2*Δ mutants displayed cell wall thickness comparable to that of the WT strain.

To identify the wall components affected, we stained cells grown in YPD for 48 h with fluorescent probes specific for individual cell wall polymers. The *pro1*Δ and *pro2*Δ mutants exhibited cell wall staining patterns indistinguishable from the WT, whereas the *pro3*Δ mutant displayed a significant reduction in the levels of total chitin and chitosan (visualized by CFW), chitosan (eosin Y, EY), and β-glucan (Congo red, CR) compared with the WT (**Fig. 5C**). However, chitooligomer (wheat germ agglutinin, WGA) and mannoprotein (concanavalin A, ConA) levels were comparable between the *pro3*Δ mutant and the WT. Notably, although all three proline biosynthetic mutants exhibited sensitivity to CFW and SDS (**Fig. 3A**), reduced staining for chitin, chitosan, and β-glucan was observed only in *pro3*Δ under the conditions examined (**Fig. 5C**). These findings indicate that Pro3, but not Pro1 or Pro2, is specifically required for normal cell wall composition and polymer homeostasis.

### Mitochondrial ROS levels and morphology are not detectably altered by disruption of proline biosynthesis

A previous study reported increased mitochondrial superoxide accumulation in the *C. neoformans put2*Δ mutant following exposure to proline (15). We therefore investigated whether disruption of proline biosynthesis alters mitochondrial ROS levels or mitochondrial morphology and distribution under basal, elevated-temperature, and host-mimicking conditions. MitoSOX Red was used to assess mitochondrial superoxide-associated fluorescence, and no significant differences were detected between the WT and the proline biosynthetic mutants within each culture condition (**Fig. S7A**). MitoTracker Green FM was used to visualize mitochondrial morphology and intracellular distribution. Fluorescence imaging revealed no obvious differences in mitochondrial morphology or distribution among the strains, and MitoTracker fluorescence intensity was also comparable between the WT and the mutants (**Fig. S7B**). In addition, nonyl acridine orange (NAO), which preferentially associates with cardiolipin in the mitochondrial inner membrane, showed comparable staining patterns and fluorescence intensities among the strains (**Fig. S7C**). Together, these results indicate that deletion of the proline biosynthetic genes did not produce detectable alterations in mitochondrial superoxide-associated fluorescence, mitochondrial morphology or distribution, or cardiolipin-associated staining under the conditions examined.

### Blocking P5C-generating routes partially restores the growth and morphological defects of *pro3*Δ but not its cell wall defects

We hypothesized that the unique defects observed in the *pro3*Δ mutant might result from an imbalance in P5C-generating metabolic flux. To test this, we generated *pro3*Δ *pro2*Δ, *pro3*Δ *car2*Δ, and *pro3*Δ *pro2*Δ *car2*Δ mutants to block the two major cytosolic P5C-generating routes mediated by Pro2 and Car2 (**Fig. 6A**). We first examined whether blocking P5C-generating routes could restore the growth defect of *pro3*Δ under mammalian tissue culture conditions (RPMI 1640 medium with 10% FBS). As a result, the *pro3*Δ mutant exhibited a prolonged lag phase compared with the WT strain. Deletion of *PRO2* in the *pro3*Δ background partially alleviated the growth defect, whereas *CAR2* deletion had little apparent effect. Consistently, the *pro3*Δ *pro2*Δ *car2*Δ mutant exhibited a growth profile similar to that of the *pro3*Δ *pro2*Δ mutant (**Fig. 6B**). These findings suggest that the partial growth recovery observed under this condition is driven predominantly by disruption of the Pro2-dependent P5C-generating pathway.

**Figure 6.**
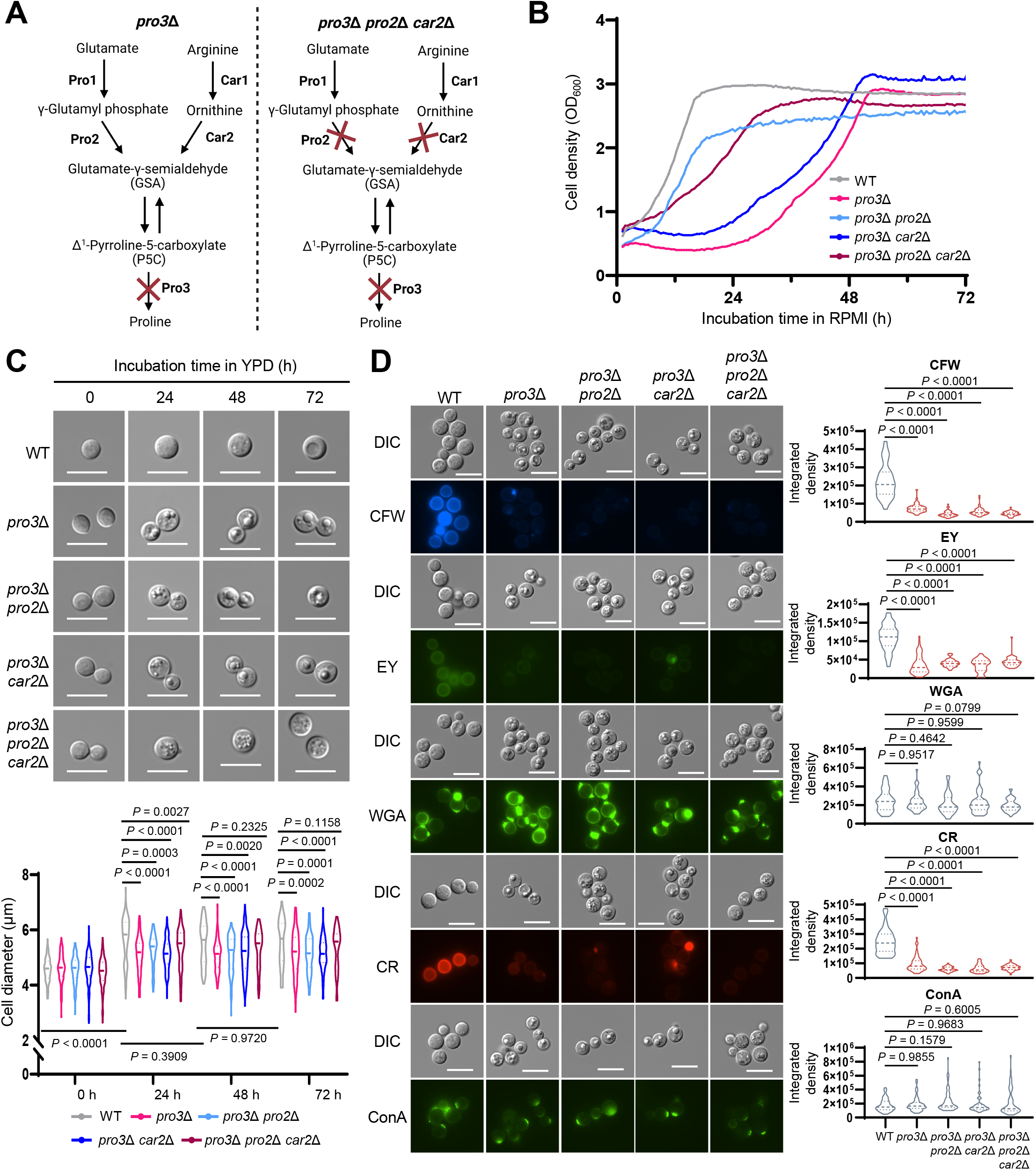
Blocking P5C-generating routes partially restores the growth and morphological defects of the *pro3*Δ mutant. (A) Simplified schematic showing the strategy used to reduce P5C-generating metabolic flux in the *pro3*Δ background. Pro2 mediates P5C production through the glutamate-derived de novo pathway, whereas Car2 mediates P5C production through the arginine/ornithine bypass. Additional deletion of *PRO2* and *CAR2* in the *pro3*Δ background was used to block the major cytosolic P5C-generating routes. This figure was designed using BioRender (https://BioRender.com/mubyso5). (B) In vitro growth assay under host-mimicking condition. The growth of *C. neoformans* wild-type (H99), *pro3*Δ, *pro3*Δ *pro2*Δ, *pro3*Δ *car2*Δ, and *pro3*Δ *pro2*Δ *car2*Δ strains was monitored in RPMI 1640 medium supplemented with 10% FBS at 37 °C using the RTS-8 bioreactor system. Cell density (OD_600_) was measured every 30 min for 72 hours. The *pro3*Δ mutant exhibited a distinct growth defect compared to the WT. (C) Cells from an overnight culture were sub-cultured into fresh YPD media at an initial OD_600_ of 0.2. Once the culture reached the mid-exponential phase (OD_600_ = 0.8), this point was defined as 0 h and the cells were further incubated for 72 h with imaging performed at 24 h intervals. Violin plots show the distribution of cell diameters (n = 100). Scale bar represents 10 µm. Statistical significance was determined by two-way ANOVA followed by Tukey’s multiple-comparison test. (D) Cell wall staining. Representative DIC and fluorescence images of cells stained with Calcofluor White (CFW), Eosin Y (EY), Wheat Germ Agglutinin (WGA), Congo Red (CR), and Concanavalin A (ConA) after 48 h of culture in YPD. Fluorescence was quantified as integrated density within individual cell outlines using ImageJ for 50 randomly selected cells per strain. Scale bar represents 10 µm. Statistical significance was determined by one-way ANOVA followed by Tukey’s multiple-comparison test.

We next assessed whether this partial growth recovery was accompanied by restoration of cell morphology. Representative DIC images showed that the *pro3*Δ mutant maintained a smaller cell morphology after prolonged incubation, whereas the *pro3*Δ *pro2*Δ *car2*Δ mutant showed a partial shift toward WT-like morphology (**Fig. 6C**). Consistent with this observation, cell diameter quantification revealed that the *pro3*Δ *pro2*Δ *car2*Δ mutant was no longer significantly different in cell size from the WT strain at later time points, in contrast to *pro3*Δ and the other mutants, which remained significantly smaller. These results indicate that blocking P5C-generating routes partially restores the morphological defect of *pro3*Δ. In contrast, cell wall staining analysis revealed that the reduced levels of chitin, chitosan, and β-glucan in the *pro3*Δ mutant were not restored in the *pro3*Δ *pro2*Δ *car2*Δ mutant, remaining comparable to those of *pro3*Δ (**Fig. 6D**). To determine whether the growth and morphological defects were associated with detectable P5C accumulation, we quantified intracellular P5C. However, no statistically significant increase in intracellular P5C was detected in *pro3*Δ relative to WT at either 30°C or 37°C (**Fig. S8**).

Taken together, these genetic data suggest that the growth and morphological defects of *pro3*Δ are not caused solely by proline deficiency but are partly associated with P5C-generating metabolic flux. In contrast, the cell wall defects of *pro3*Δ were not rescued by blocking P5C-generating pathways, suggesting that the cell wall phenotype is mechanistically distinct from the growth and morphological defects and may involve P5C homeostasis or an additional, as-yet-uncharacterized function of Pro3.

### Aap6 is a proline-responsive amino acid permease but is dispensable for proline-dependent growth

Because *pro1*Δ and *pro2*Δ retained full virulence and exogenous proline rescued multiple phenotypes of the proline biosynthetic mutants, we asked whether *C. neoformans* uses exogenous proline to bypass biosynthetic deficiencies. Since the specific proline transporters remain uncharacterized, we profiled ten amino acid permease (*AAP*) homologs (*AAP1-8*, *MUP1*, and *MUP3*) by qRT-PCR after transfer from YPD to YNB medium lacking ammonium sulfate, with or without proline (28). The 15-and 30-min time points were selected to capture early transcriptional responses following the nutrient shift. Differences in statistical significance between these time points may reflect transient expression responses or experimental variability. A significant effect at one time point but not the other does not itself establish a difference between time points. Among the genes tested, *AAP6* (CNAG_07449) was strongly induced in response to proline. At 15 min, its transcript was approximately 40-fold higher than that in the YPD control sampled immediately before the nutrient shift (time zero) and more than 6-fold higher than that under nitrogen-starved conditions without proline at the same time point (**Fig. 7A**). To determine whether this induction reflects a general response to amino acid availability, we further analyzed *AAP6* expression in response to arginine supplementation. Arginine supplementation did not induce *AAP6* expression, indicating that its induction is not a universal response to amino acids (**Fig. S9A**). This selective induction identifies Aap6 as a candidate proline-responsive permease gene.

**Figure 7.**
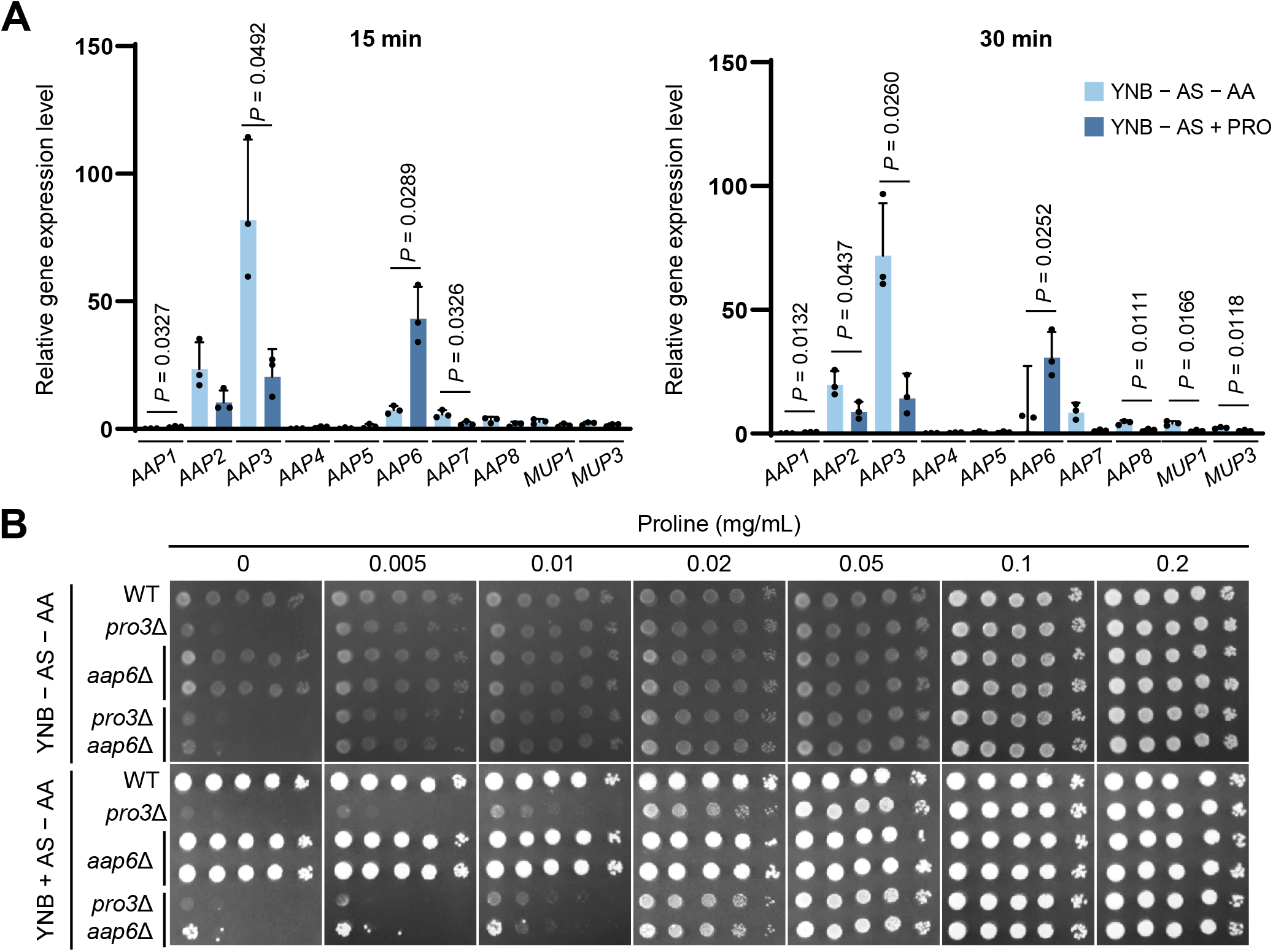
Aap6 is a putative proline-responsive amino acid permease. (A) Expression profiles of putative amino acid permease genes. *Cryptococcus neoformans* H99 cells were grown overnight in YPD at 30°C and then subcultured in YPD from an initial OD_600_ of 0.2 until reaching an OD_600_ of 0.8. Cells were washed three times and transferred to YNB medium without ammonium sulfate (AS) and with or without proline (0.2 mg/mL) for 15 or 30 min. Transcript levels of *AAP1-8, MUP1,* and *MUP3* were quantified by qRT-PCR and normalized to *ACT1* expression. Relative expression was calculated using the 2^□ΔΔC^ method, with the YPD control sampled immediately before the nutrient shift (time zero) used as the calibrator and assigned a value of 1. Data are presented as mean ± SEM from three biological replicates, with individual data points shown. Statistical comparisons were performed using unpaired two-tailed t-tests with Welch’s correction for each gene separately at each time point. (B) Proline supplementation spotting assay. WT, *pro3*Δ, *aap6*Δ, and *pro3*Δ *aap6*Δ strains were grown overnight in YPD at 30°C and then spotted in 10-fold serial dilutions on Yeast Nitrogen Base (YNB) minimal media with (+) or without (-) ammonium sulfate (AS). For nutrient rescue assays, media were individually supplemented with different concentrations of proline.

To investigate the physiological role of Aap6 in proline acquisition, we generated *aap6*Δ and *pro3*Δ *aap6*Δ mutants and assessed growth on YNB minimal medium supplemented with a range of proline concentrations (0–0.2 mg/mL). The *aap6*Δ single mutant grew indistinguishably from the WT across all proline concentrations tested, displaying no detectable auxotrophy. Likewise, the *pro3*Δ *aap6*Δ double mutant exhibited proline-dependent growth recovery indistinguishable from that of the *pro3*Δ single mutant, with no additive defect (**Fig. 7B**). These results indicate that deletion of *AAP6* alone is insufficient to impair proline-dependent growth under the conditions tested.

To assess whether the absence of a growth phenotype in *aap6*Δ resulted from transcriptional compensation by other amino acid permeases, we compared the expression profiles of all ten amino acid permeases (*AAP1–8*, *MUP1*, and *MUP3*) between the WT and *aap6*Δ strains under proline supplementation (**Fig. S9B**). Notably, none of the permease genes exhibited significant upregulation in *aap6*Δ relative to the WT, indicating the absence of compensatory transcriptional induction. These findings suggest that other amino acid permeases, expressed constitutively, can mediate proline uptake and thereby mask the *aap6*Δ phenotype.

## DISCUSSION

The physiological and pathogenic contributions of the de novo proline biosynthetic pathway remain incompletely understood in human fungal pathogens, where research has focused predominantly on proline catabolism. To our knowledge, this study provides the first comprehensive and systematic functional characterization of this pathway in a human fungal pathogen, *C. neoformans* (summarized in **Fig. 8**). Our findings demonstrate that the terminal enzyme Pro3, but not the upstream enzymes Pro1 and Pro2, serves as an indispensable regulator of physiological proline homeostasis, cell wall integrity, and pathobiological fitness in *C. neoformans*.

**Figure 8.**
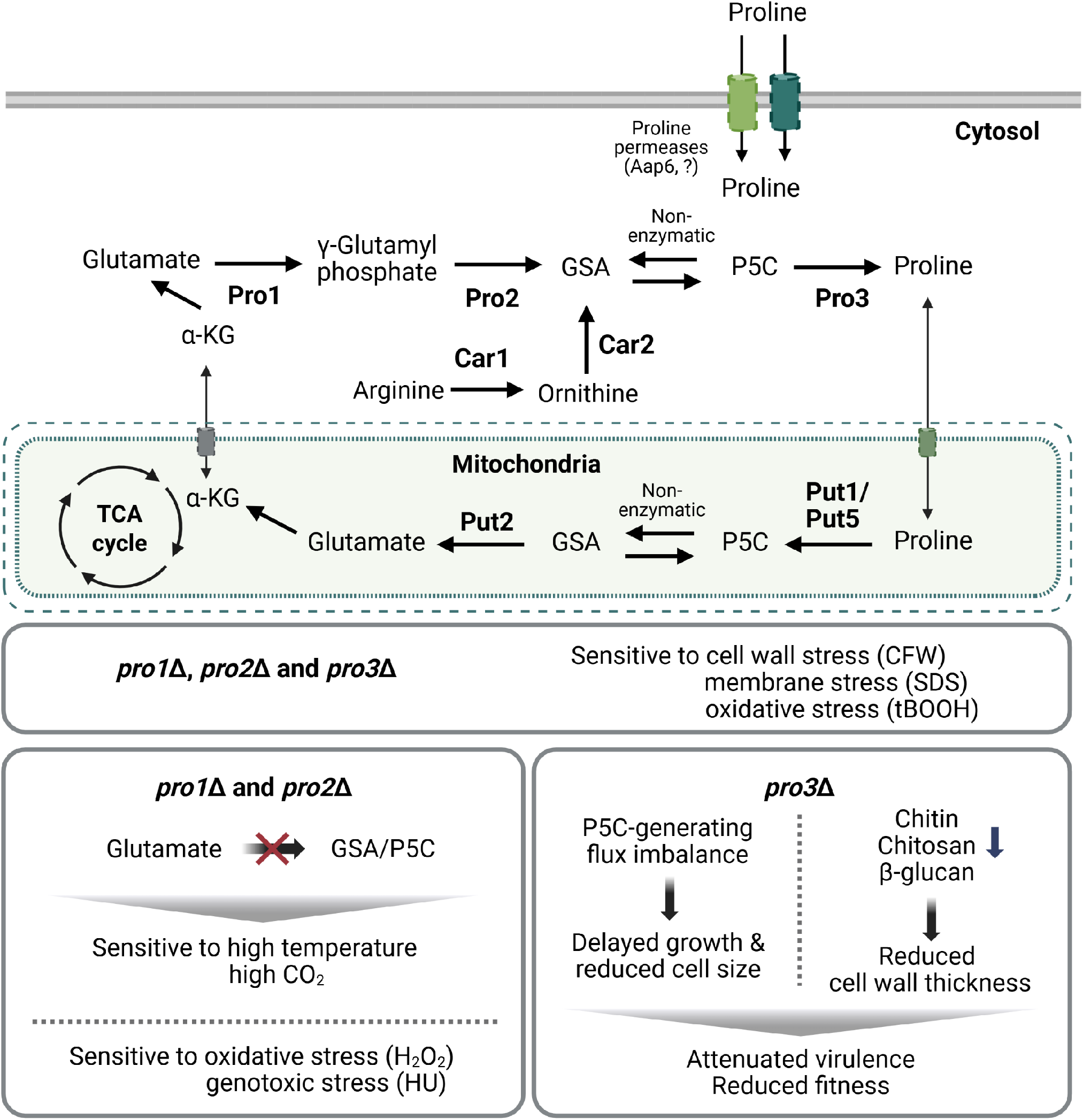
Proposed model for the role of the proline biosynthetic pathway in the physiology and pathobiology of *C. neoformans*. Schematic summary of proline metabolism in *C. neoformans* and the distinct pathobiological consequences of disrupting individual biosynthetic steps. In the cytosol, proline is synthesized de novo from glutamate through the sequential actions of Pro1 (γ-glutamyl kinase), Pro2 (γ-glutamyl phosphate reductase), and Pro3 (P5C reductase), with glutamate-γ-semialdehyde (GSA) and Δ^1^-pyrroline-5-carboxylate (P5C) as intermediates. An alternative arginine-mediated bypass, in which arginine is converted to ornithine by Car1 and subsequently to GSA/P5C by Car2, converges on Pro3 for the terminal reduction of P5C to proline. Exogenous proline is imported across the plasma membrane by proline permeases. In the mitochondria, proline is catabolized back to glutamate via Put1/Put5 and Put2, feeding α-ketoglutarate (α-KG) into the tricarboxylic acid (TCA) cycle. The dashed and solid boundaries represent the mitochondrial outer and inner membranes, respectively, with the dashed boundary indicating the greater permeability of the outer membrane to small solutes. All three proline biosynthetic mutants exhibit sensitivity to tert-butyl hydroperoxide (tBOOH), calcofluor white (CFW), and sodium dodecyl sulfate (SDS). Loss of Pro1 or Pro2 blocks the glutamate-derived route to GSA/P5C and is associated with sensitivity to high temperature, high CO_2_, hydrogen peroxide (H_2_O_2_), and hydroxyurea (HU). Loss of Pro3 is associated with an imbalance in P5C-generating metabolic flux, delayed growth under host-mimicking conditions, reduced cell size, and reduced levels of chitin, chitosan, and β-glucan together with decreased cell wall thickness. Disruption of the major cytosolic P5C-generating routes mediated by Pro2 and Car2 partially alleviates the growth and morphological defects of *pro3*Δ, whereas the cell wall defects persist. The *pro3*Δ mutant also exhibits attenuated virulence in a murine model. This summary figure was designed using BioRender (https://BioRender.com/mubyso5).

A notable distinction in our in vivo data is the divergence in virulence between the upstream biosynthetic mutants (*pro1*Δ and *pro2*Δ) and the terminal mutant (*pro3*Δ). Although all three strains are strict proline auxotrophs, only *pro3*Δ exhibits significant attenuation in the murine infection model, indicating that its virulence defect is unlikely to reflect simple nutrient limitation alone. Because proline supplementation did not restore the virulence of *pro3*Δ in our murine model, we considered whether the supplementation regimen was sufficient to increase host proline availability. In a previous study of pregnant C57BL/6J mice, supplementation of the diet with 0.5% L-proline significantly increased maternal plasma proline concentrations from approximately 143 to 180 μM (29). This supplemented plasma concentration (∼0.021 mg/mL) overlaps with the range associated with improved *pro3*Δ growth in our in vitro assay. However, because plasma and tissue proline concentrations were not directly measured in our supplemented mice, the actual proline availability experienced by fungal cells in vivo remains uncertain. The different supplementation routes and mouse models further limit direct comparison between the two studies.

In contrast to our findings in *C. neoformans*, deletion of *PRO3* in the plant pathogen *C. parasitica* did not significantly affect virulence or asexual sporulation under the conditions examined (18), suggesting that the contribution of Pro3 to fungal pathogenesis may differ among species. In *C. albicans*, the disruption of either the first (Put1) or second (Put2) catabolic step results in attenuated virulence (10). This attenuation is linked to the role of proline degradation in hyphal morphogenesis and P5C-associated toxicity. Unlike *C. albicans*, systemic infection by *C. neoformans* does not depend on hyphal formation. Indeed, in *C. neoformans*, the *put1*Δ *put5*Δ double mutant maintains wild-type levels of virulence, whereas the *put2*Δ mutant is completely avirulent (15). This difference in virulence suggests that the avirulence of *put2*Δ cannot be explained solely by the loss of proline-derived energy production and may instead involve P5C-associated toxicity.

Similar P5C-associated toxicity has been reported in *Arabidopsis thaliana p5cdh* mutants exposed to excess proline, supporting the broader concept that impaired P5C turnover can disrupt cellular physiology (30). Although disruption of either Put2 or Pro3 may alter P5C-associated metabolic flux, the phenotypic outcomes differ substantially between the catabolic *put2*Δ mutant and the biosynthetic *pro3*Δ mutant. One possible explanation for the absence of detectable P5C accumulation in *pro3*Δ is regulation of the glutamate-derived P5C biosynthetic pathway. If feedback inhibition of the initial biosynthetic step, as described in other organisms (31, 32), also operates in *C. neoformans*, P5C production from glutamate could be constrained. Consequently, *pro3*Δ may experience a more limited or transient P5C-associated metabolic imbalance, which could contribute to attenuated virulence rather than absolute avirulence. Such a perturbation may occur without producing a detectable increase in the intracellular P5C pool, although its nature and duration remain unresolved.

The cell wall defects of the *pro3*Δ mutant—the marked reduction in chitin, chitosan, and β-glucan—were not restored in the *pro3*Δ *pro2*Δ *car2*Δ triple mutant, even though the growth and morphological defects were partially rescued. Whereas *pro3*Δ disrupts P5C reduction, the triple mutant additionally disrupts the major cytosolic P5C-generating routes; nevertheless, the cell wall defects persist in both strains. These findings suggest that the cell wall requirement may reflect an additional function of Pro3 beyond its role in P5C reduction. We cannot fully exclude an alternative interpretation in which cell wall integrity depends on balanced P5C homeostasis, such that distinct perturbations of P5C-associated metabolism in *pro3*Δ and the triple mutant impair cell wall integrity through imbalance in either direction. The reduction in staining for several cell wall polymers, together with decreased wall thickness, suggests a broader defect in cell wall production or assembly rather than an isolated alteration in a single polymer. Pro3-dependent metabolism may therefore support the cellular processes required to coordinate cell wall synthesis and maintenance. If Pro3 indeed has a P5C-independent function, one possibility is that Pro3 supplies a localized or temporally critical proline pool required for proper cell wall polymer biosynthesis or assembly; alternatively, it may contribute to cell wall homeostasis through a moonlighting or regulatory function beyond its canonical reductase activity. Distinguishing between these possibilities will require further mechanistic investigation.

Proline supplementation restored pigmentation of *pro3*Δ on solid media but failed to restore cell-associated laccase activity in the liquid assay. Although *LAC1* transcript levels increased following proline supplementation, the expression and activity assays used different induction conditions, limiting direct comparison. Cryptococcal laccase is strongly associated with the cell wall, and proper intracellular trafficking contributes to its cell wall localization (33, 34). These observations raise the possibility that the altered cell wall composition of *pro3*Δ affects laccase association with the cell surface, thereby reducing the activity detected in washed cells. Furthermore, laccase activity has been demonstrated in extracellular vesicles isolated from culture supernatants, indicating that enzymatically active laccase can also occur outside the cell-associated fraction (35). Extracellular activity could contribute to pigment formation on solid media but would not be captured by our liquid assay, in which the culture supernatant was discarded before activity measurement. Thus, reduced retention of active laccase at the cell surface could account for the recovery of pigmentation on solid media despite persistently low activity in the liquid assay following proline supplementation. However, laccase abundance, localization, and extracellular activity were not directly measured in *pro3*Δ; altered laccase distribution therefore remains to be determined.

The ability of all proline biosynthetic mutants to retain at least partial pathogenic potential suggests that *C. neoformans* can access host-derived nutrients in some infection niches. Although our screen identified *AAP6* as a proline-responsive permease gene at the transcriptional level, deletion of *AAP6* caused no detectable defect in proline-dependent growth, even in the *pro3*Δ background. This suggests that other permeases can support sufficient proline uptake in the absence of Aap6, without excluding a contribution from Aap6 itself. The absence of compensatory transcriptional induction among the other permease genes examined is consistent with uptake supported by their basal expression. Such functional redundancy could mask the contribution of Aap6 in uptake assays using the single deletion mutant. Identifying additional candidate proline transporters and combining their deletions with *AAP6* deletion, followed by direct uptake measurements, will therefore help resolve their individual and overlapping contributions. Aap6 thus remains a candidate permease whose role in proline transport requires direct validation.

Our study reveals that Pro3 serves as an obligatory convergence point for two distinct metabolic routes: the glutamate-derived de novo pathway and the nitrogen-regulated arginine bypass mediated by Car1 and Car2. The transcriptional repression of *CAR1* and *CAR2* in the presence of ammonium sulfate indicates that the arginine bypass is activated specifically under preferred-nitrogen-limited conditions. During infection, activation of this bypass may confer an advantage within nutrient-restricted microenvironments. The medium-dependent capsule defects highlight a metabolic requirement dictated by exogenous nitrogen availability. In Littman’s minimal medium, which lacks both proline and arginine, disruption of any biosynthetic step markedly impairs capsule formation, consistent with limited proline availability (36). Standard DMEM (Gibco) contains L-arginine (∼400 μM) but lacks proline. At 10% concentration, commercial FBS contains approximately 1.1 μg/mL proline, 1.5 μg/mL ornithine, and 0.34 μg/mL arginine on average (37, 38). In these environments, *pro1*Δ and *pro2*Δ successfully maintain capsule production because they possess the metabolic flexibility to scavenge exogenous arginine or ornithine and synthesize proline via the functional Car1-Car2-Pro3 bypass. Conversely, *pro3*Δ lacks this terminal convergence enzyme and is therefore strictly reliant on free exogenous proline. Thus, the ability to exploit alternative nitrogen sources through the Pro3-mediated arginine bypass may support *C. neoformans* fitness under varying environmental constraints. The complete restoration of capsule formation by exogenous proline suggests that the capsule defects primarily reflect proline auxotrophy and the associated growth or metabolic limitations, rather than a capsule-specific function of the biosynthetic pathway. Thus, the medium-dependent capsule phenotypes provide an example of how alternative nitrogen availability can differentially expose the requirements for individual steps of proline biosynthesis.

Our study reveals a clear divergence between the initial (Pro1, Pro2) and terminal (Pro3) steps under host-mimicking stresses. Although the *pro3*Δ mutant exhibits the most severe defects in virulence and cell wall integrity, it maintains near-WT levels of thermotolerance and adaptation to 5% CO_2_. In contrast, *pro1*Δ and *pro2*Δ are hypersensitive to high temperature and 5% CO_2_, and these vulnerabilities cannot be rescued by exogenous proline. Strikingly, the addition of ornithine together with proline partially restored growth of *pro1*Δ and *pro2*Δ under these conditions, whereas ornithine alone did not. The partial growth recovery following combined proline and ornithine supplementation suggests that the defects of *pro1*Δ and *pro2*Δ are not attributable solely to proline deficiency. Ornithine is also used in polyamine biosynthesis, being converted to putrescine by ornithine decarboxylase and subsequently to spermidine by spermidine synthase. Because spermidine biosynthesis is functional in *C. neoformans* (39), the additional benefit of ornithine supplementation in the presence of proline may partly involve its utilization for polyamine production. This possibility is supported by studies in *S. cerevisiae* showing that polyamine homeostasis contributes to heat-and oxidative-stress tolerance (40, 41). Whether a similar mechanism operates in *C. neoformans*, and the relative contributions of polyamine metabolism, P5C, and other associated metabolites, remain to be clarified.

The discrepancy between the macrophage assays and lung fungal burden may reflect niche-specific differences in nutrient availability. While *pro1*Δ and *pro2*Δ maintain WT levels of fungal burden in the murine lung, they exhibit significantly impaired survival within J774A.1 macrophages. This suggests that the macrophage phagolysosome may present a more sequestered and nutrient-deprived environment compared with the lung niche. Our capsule assays are consistent with the ability of *pro1*Δ and *pro2*Δ to bypass their biosynthetic defects when exogenous proline or arginine-derived substrates are available. One possible explanation is that the intracellular macrophage environment provides less accessible proline or related nitrogen sources than other lung-associated niches, although this was not directly examined.

Collectively, this study identifies the proline biosynthetic pathway as a critical contributor to *C. neoformans* pathobiology, extending beyond its canonical role in amino acid provision. Our findings suggest that Pro3 acts as a metabolic gatekeeper with dual roles in balancing P5C-generating flux and maintaining cell wall integrity, the latter potentially involving a mechanism distinct from P5C flux control. Future investigations should define the mechanism by which Pro3 supports cell wall homeostasis— a function not restored by disrupting the major cytosolic P5C-generating routes—as well as the intricate crosstalk between proline metabolism and global nutrient-sensing networks, such as nitrogen catabolite repression and central carbon metabolism.

## MATERIALS AND METHODS

### Ethics statement

All animal experiments were conducted in strict accordance with institutional and national guidelines for the care and use of laboratory animals. The study protocol was reviewed and approved by the Institutional Animal Care and Use Committee (IACUC) of Jeonbuk National University (approval number: NON2026-095). All efforts were made to minimize animal suffering and to reduce the number of animals used.

### Strains and culture conditions

The *C. neoformans* strains used in this study are listed in **Table S1**. MATα strains were derived from the WT H99L (serotype A) background (42), whereas MAT**a** strains were derived from YL99 (43). Cryogenic stocks were maintained at -80°C in 20% glycerol, and routine propagation was conducted on YPD agar (1% yeast extract, 2% peptone, 2% glucose) at 30°C for 48–72 h. For liquid cultivation, cells were aerated in YPD broth at 30°C with shaking at 200 rpm.

### Strain construction

Targeted gene disruption was performed using a split-marker/double-joint PCR method (44, 45). In the first round of PCR, the 5′ and 3′ flanking regions (∼800 bp) of the target genes were amplified from *C. neoformans* H99 genomic DNA using the primer pairs L1/L2 and R1/R2, respectively. Simultaneously, the selection markers—nourseothricin acetyltransferase (*NAT*), neomycin phosphotransferase (*NEO^R^*), or hygromycin B phosphotransferase (*HYG^R^*)—were amplified from the plasmids pNAT, pNEO, or pHYG using universal primers (M13FWD and M13REV). In the second round of overlap extension PCR, the 5′ flanking region was fused to a portion of the selection marker, and the 3′ flanking region was fused to the remaining part, generating 5′- and 3′- split gene disruption cassettes. These cassettes were further amplified using internal split primers (L1/SM2 and SM1/R2). Single deletion mutants were constructed using the *NAT* or *NEO* marker, and additional gene deletions in these backgrounds were performed sequentially using a different marker (*NAT*, *NEO*, or *HYG*) to avoid marker overlap. The amplified split-marker cassettes were introduced into *C. neoformans* via biolistic transformation using the PDS-1000/He particle delivery system (Bio-Rad). To prepare recipient cells, the H99 strain or the parental strain was cultured overnight in YPD broth at 30°C, and cells were collected and resuspended in 5 mL YPD. Around 1 mL of this suspension was spread onto YPD plates containing 1 M sorbitol for 3 h. For each transformation, 0.6 μm gold microcarrier beads (Bio-Rad) were coated with the disruption DNA cassettes in the presence of CaCl_2_ and spermidine. These were then biolistically transformed into cells. Post-transformation, cells were incubated for 4 h at 30°C to restore membrane integrity and then transferred to the selection medium containing nourseothricin (100 μg/mL), G418 (45 μg/mL), or hygromycin B (300 μg/mL). Correct integration of the cassettes was verified by diagnostic PCR, and Southern blot analysis.

To generate complemented strains, full-length DNA fragments encompassing the native promoter, open reading frame (ORF), and terminator of *PRO1*, *PRO2*, and *PRO3* were amplified and cloned into a pNEO-based vector using Gibson assembly to generate pNEO_*PRO1*, pNEO_*PRO2*, and pNEO_*PRO3*. The resulting plasmids were linearized by digestion with appropriate restriction enzymes—ClaI for *PRO1*, PpuMI for *PRO2*, and StuI for *PRO3*—to facilitate stable re-integration into the native loci. The linearized constructs were introduced into the corresponding deletion mutants via biolistic transformation. Targeted insertion was verified by diagnostic PCR, and phenotypic restoration was confirmed.

### Identification, phylogenetic analysis, and structural modeling

Orthologs of Pro1, Pro2, and Pro3 in *C. neoformans* were identified via BLASTp analysis against the H99 genome database, using characterized sequences from *Saccharomyces cerevisiae* and *Homo sapiens* as queries. Orthologous protein sequences from diverse taxa were retrieved from the National Center for Biotechnology Information (NCBI; *H. sapiens*), EcoCyc (*Escherichia coli*), FungiDB (*C. neoformans, S. cerevisiae, Schizosaccharomyces pombe, Candida albicans, Aspergillus fumigatus, Ustilago maydis*), The Arabidopsis Information Resource (TAIR; *Arabidopsis thaliana*), Wormbase (*Caenorhabditis elegans*) and FlyBase (*Drosophila melanogaster*). Domain architectures and functional motifs were confirmed using InterPro and UniProt. Phylogenetic trees were constructed using the maximum likelihood method based on the JTT matrix-based model with MEGA 11 software (46, 47). To assess structural conservation, protein tertiary structures were predicted using AlphaFold3 (48). Structural alignments were subsequently conducted with *S. cerevisiae* orthologs and human P5CS or PYCR3 to calculate sequence identities and similarities.

### Growth and stress sensitivity assays

To evaluate nutrient utilization and stress sensitivity, strains were cultured overnight in YPD broth at 30°C, serially diluted 10-fold (ranging from 10^0^ to 10^-4^) or 2-fold (ranging from 2^0^ to 2^-11^) using sterile distilled water. For nutrient utilization assays, cells were washed twice with 1×PBS and spotted onto Yeast Nitrogen Base (YNB) agar plates (containing 2% D-glucose) with or without ammonium sulfate. As indicated, the media were supplemented with specific amino acids, including L-proline, L-arginine, or L-glutamic acid monosodium salt monohydrate. The plates were grown at 30°C and imaged after 2–3 days of incubation. Temperature tolerance was examined by incubating the plates at 30°C, 37°C, and 38°C. CO_2_ tolerance was assessed by incubating plates in a 5% CO_2_ incubator at 37°C. The plates were imaged after 2–3 days of incubation. For stress sensitivity assays, cells were spotted onto YPD or YNB agar plates containing various stressors. These included oxidative stressors [hydrogen peroxide (H_2_O_2_), *tert-*butyl hydroperoxide (tBOOH), menadione (MD) or diamide (DIA)], ER stressor [dithiothreitol (DTT) or tunicamycin (TM)], a membrane stressor (SDS), cell wall stressors [Congo red (CR), caffeine and calcofluor white (CFW)], an osmotic stressor (sorbitol), cation and salt stressors (NaCl or KCl), and antifungal agents [amphotericin B (AMB), fluconazole (FLC), fludioxonil (FDX), flucytosine (5-FC)]. The plates were grown at 30°C and imaged after 2–3 days of incubation under stress conditions.

### Virulence factor analysis

To evaluate melanin production, cells were pre-cultured in YPD liquid medium for 16 h at 30°C, harvested, and washed twice with phosphate-buffered saline (PBS). The resulting cell suspensions were spotted onto Niger seed (35 g niger seed per L), L-DOPA, or epinephrine agar plates (1 g L-asparagine, 3 g KH_2_PO_4_, 250 mg MgSO_4_, 1 mg thiamine, 5 μg biotin, and 100 mg L-DOPA or epinephrine hydrochloride per L). The plates were incubated at 37°C for 2 to 3 days, and imaged. To examine the polysaccharide capsule formation, cells were cultured overnight in YPD, washed twice with PBS, and transferred to capsule induction media, including Dulbecco’s modified Eagle medium, Littman’s (36), and 10% fetal bovine serum (FBS) agar plates. After a 48 h incubation at 37°C, cells were stained with India ink and observed under differential interference contrast (DIC) microscopy (ECLIPSE Ni; Nikon) equipped with a digital camera (DS-Qi2; Nikon). Capsule thickness was quantified by calculating the difference between the total diameter and the cell body diameter (22). Significant differences in capsule production were determined through two-way analysis of variance (ANOVA), followed by Tukey’s or Šídák’s multiple-comparison test.

For the liquid laccase activity assay (49), strains were grown overnight in YPD at 30°C, washed three times with PBS, and inoculated at 1 × 10^7^ cells/mL into YNB containing ammonium sulfate, 0.1% glucose with or without L-proline (0.2 mg/mL). After incubation at 30°C with shaking for 12 h, cell numbers were determined, and 2 × 10 cells from each culture were collected and washed three times with YNB. Cells were then resuspended in 2 mL of YNB containing ammonium sulfate, 0.1% glucose, and 10 mM L-DOPA and incubated at 30°C with shaking for 10 min. Cells were removed by centrifugation, and the absorbance of the supernatant was measured at 475 nm. Cell-free L-DOPA medium incubated under the same conditions was used as a blank, and laccase activity was expressed as the blank-corrected OD_475_ value. Statistical analysis was conducted using GraphPad Prism 11. Significant differences were determined through two-way analysis of variance (ANOVA), followed by Tukey’s multiple-comparison test.

### Sexual development assay

To assess the impact of proline biosynthesis on the sexual differentiation of *C. neoformans*, unilateral and bilateral mating efficiencies were evaluated using opposite mating-type strains. The *MAT*α strains (WT H99, *pro1*Δ, *pro2*Δ, *pro3*Δ, and complemented strains) and the *MAT***a** strains (YL99, *pro1*Δ, *pro2*Δ, *pro3*Δ) were cultured in liquid YPD medium at 30°C for 16 h. Following incubation, cells were harvested and washed three times with PBS. For the mating assays, *MAT*α cells were mixed with the corresponding *MAT***a** cells at a final concentration of 10^7^ cells/mL each. These mixtures were spotted onto V8 agar medium (pH 5.0) with or without L-proline. The plates were then incubated at 25°C in the dark for 7 to 14 days. The progression of filamentous growth was regularly monitored. Mating filaments were observed and imaged using DIC microscopy (ECLIPSE Ni; Nikon) equipped with a digital camera (DS-Qi2; Nikon).

### RNA extraction and quantitative RT-PCR

qRT-PCR was performed to analyze nutrient-responsive gene expression and *LAC1* transcript levels in WT and mutant strains. For nutrient-responsive gene expression analysis, mid-log-phase cells were harvested, washed three times with sterile distilled water, and transferred to YNB containing 2% D-glucose, with or without ammonium sulfate and with or without L-proline or L-arginine, as indicated. Cells were harvested after 15 or 30 min of incubation.

For *LAC1* expression analysis, WT and mutant strains were cultured overnight in 50 mL of YPD at 30°C with shaking and then subcultured into 80 mL of fresh YPD until the OD_600_ reached 0.6–0.8. Cells from a 40-mL aliquot were collected as the YPD basal control. Cells from the remaining 40 mL were harvested by centrifugation, washed three times with PBS, and resuspended in 40 mL of YNB containing ammonium sulfate without glucose, with or without L-proline supplementation (0.2 mg/mL). After incubation at 30°C with shaking for 2 h, cells were harvested.

All harvested samples were rapidly frozen in liquid nitrogen and lyophilized overnight. Total RNA was isolated using easy-BLUE reagent (iNtRON Biotechnology), and cDNA was synthesized from 3 μg of purified total RNA using reverse transcriptase (Thermo Scientific) according to the manufacturer’s instructions. qRT-PCR was performed using a CFX96 Real-Time System (Bio-Rad) with gene-specific primers listed in Table S2. Target transcript levels were normalized to *ACT1* expression, and relative expression was calculated using the 2 ^ΔΔC^_t_ method (50). For nutrient-responsive expression analysis, either the WT sample incubated in YPD or YNB without ammonium sulfate or amino acid supplementation at the corresponding time point was used as the calibrator, as specified in the corresponding figure legends. The calibrator was assigned a value of 1. For *LAC1* expression analysis, the WT sample incubated in glucose-free YNB containing ammonium sulfate without proline was used as the calibrator and assigned a value of 1. Experiments were performed with at least three independent biological replicates, each analyzed in technical triplicate.

### Murine infection

Animal experiments were conducted at the Core Facility Center for Zoonosis Research (Jeonbuk National University, South Korea). SPF/VAF-confirmed inbred 6-week-old female BALB/cAnNCrlOri mice (ORIENT BIO INC.) were acclimatized to the breeding environment for one week before the experiment. For intranasal infection, *C. neoformans* strains were cultured overnight in YPD at 30°C, washed twice with PBS, and resuspended at the desired concentration. Mice were anesthetized with isoflurane, and 5 × 10^5^ yeast cells in 30 μL PBS were administered intranasally. Mice were monitored daily for survival and clinical signs of disease. Humane endpoints were applied according to IACUC guidelines, including 25% weight loss, neurological symptoms, or severe lethargy. Survival data were recorded until predetermined endpoints were reached. To assess whether exogenous proline could restore virulence, mice were provided with continuous dietary proline supplementation beginning one day prior to infection and maintained throughout the experiment. L-proline was dissolved in drinking water at a concentration of 0.5% (w/v), and solutions were replaced every 2–3 days to maintain stability. Infection procedures and survival monitoring were conducted as described above. To determine tissue fungal burdens, mice were euthanized at 14 days post-infection. Lungs and brains were harvested, weighed, and homogenized in PBS using a mechanical tissue homogenizer. Serial 10-fold dilutions of homogenates were plated on YPD agar supplemented with 100 μg/mL chloramphenicol and incubated at 30°C for 48 hours. Colony-forming units (CFUs) were counted and expressed as CFU per gram of tissue.

### Drosophila melanogaster infection

*Drosophila melanogaster* w^1118^ flies were reared on standard cornmeal-agar medium obtained from the Bloomington Stock Center at 25°C under a 12-h light/12-h dark cycle. Cells from *C. neoformans* wild-type and mutant strains were grown overnight in YPD medium at 30°C. Cells were collected, washed three times with PBS, and adjusted to a final concentration of 4 × 10^8^ cells/mL. The suspensions were then centrifuged, and the cell pellets were resuspended in PBS supplemented with 1% (w/v) blue food dye (FD&C Blue #1) to facilitate visualization of inoculation. For the survival assay, 4- to 5-day-old adult female flies were anesthetized with CO_2_ and infected by pricking the dorsal thorax with an HPLC needle (33G, 10 mm) that had been dipped into the fungal suspension. Approximately 60 flies were infected per strain and distributed into six vials, with 10 flies per vial. Each vial contained standard cornmeal-agar medium. Infected flies were maintained at 30°C, and survival was monitored daily until 5 days post-infection (dpi). Survival data were analyzed by the Kaplan–Meier method using GraphPad Prism 11, and statistical significance was determined by the log-rank (Mantel–Cox) test.

### In vitro growth assay under host-mimicking conditions

Growth under host-mimicking conditions was monitored using an automated multichannel bioreactor (RTS-8, Biosan). *C. neoformans* strains were cultured in YPD at 30°C for 48 h, harvested, and washed with PBS. Cells were resuspended in RPMI 1640 medium (pH 7.4) supplemented with 50 mM HEPES, 0.5 g/L NaHCO_3_, and 10% fetal bovine serum (FBS), at an initial OD_600_ of 0.2. The cultures were incubated at 37°C in the RTS-8 system, and cell density (OD_600_) was recorded automatically every 30 min for 72 h.

### Macrophage phagocytosis and intracellular survival assay

The J774A.1 murine macrophage-like cell line was obtained from the Korean Cell Line Bank (KCLB). The cells were maintained in DMEM medium supplemented with 10% heat-inactivated FBS, 1% penicillin-streptomycin, and 1% MEM non-essential amino acid solution at 37°C, 5% CO_2_ incubator. For the assays, 1×10^5^ viable macrophages were seeded into each well of a 96-well tissue culture plate and incubated for 16–18 h to ensure adherence. Prior to infection, the macrophages were activated by replacing the medium with fresh DMEM containing 10 nM phorbol 12-myristate 13-acetate (PMA) for 1 h at 37°C. *C. neoformans* strains were grown overnight in YPD, re-inoculated into fresh YPD at an initial OD600 of 0.3, and cultured for 16–18 h at 30°C. The fungal cells were harvested, washed twice with sterile PBS, and adjusted to a concentration of 1×10^6^ cells/mL. For opsonization, 10^6^ yeast cells were incubated with 2 μg of anti-capsular monoclonal antibody 18B7 in macrophage medium for 1 h at 37°C (51). 1×10^5^ of opsonized fungal cells were introduced to the activated macrophages at a multiplicity of infection (MOI) of 1. After 1 h of co-culture at 37°C, each well was washed three times with PBS to remove non-adherent extracellular yeast.

To determine the initial phagocytic efficiency, the macrophages were lysed by adding distilled water and incubating on ice for at least 30 min. The lysates were collected, serially diluted, and plated onto YPD agar containing chloramphenicol (30 μg/mL). To evaluate intracellular survival and proliferation, the infected macrophages were further incubated in fresh medium for 24 h following the initial 1-h co-culture and subsequent washes. At the specified time point, the macrophages were lysed using the same method. Fungal survival was quantified by counting CFUs after 2–3 days of incubation on YPD plates. To verify the actual number of fungal cells used for the initial infection, the concentration of the input fungal cell suspension was confirmed by serial dilution and quantitative plating on YPD agar. The resulting CFU counts from the macrophage assays were then normalized to this verified input fungal concentration. Statistical significance was determined using one-way ANOVA followed by Tukey’s multiple-comparison test in GraphPad Prism 11.

### Cell size and morphological analysis

To evaluate cell size and morphology, strains were initially cultured overnight in liquid YPD medium at 30°C and subsequently re-inoculated into fresh YPD at a starting OD_600_ of 0.2. The cultures were grown until they reached a mid-log phase (OD_600_ = 0.8), which was defined as 0-h time point. Following this baseline, cellular morphology and size were monitored at specific time intervals: 24 h, 48 h, and 72 h of incubation. The cells were observed using a DIC microscope (ECLIPSE Ni; Nikon) equipped with a digital camera (DS-Qi2; Nikon). For quantitative assessment of cell size, the diameter of the cell body was measured for at least 100 randomly selected cells per strain using ImageJ/Fiji software. Statistical significance was determined using two-way ANOVA, followed by Tukey’s multiple comparison test in GraphPad Prism 11.

### Transmission electron microscopy for ultrastructural analysis

For ultrastructural analysis of the cell wall, cells were grown in liquid YPD at 30 °C for 48 h with shaking. Cells were harvested by centrifugation (3,000 rpm, 1 min) and washed once with 0.1 M sodium phosphate buffer (pH 7.4). The cell pellets were resuspended in fixative containing 2.5% glutaraldehyde and 2% paraformaldehyde in 0.1 M sodium phosphate buffer (pH 7.4) and fixed for 2 h at room temperature in the dark, followed by overnight incubation at 4 °C. After primary fixation, cells were washed with 0.1 M sodium phosphate buffer, post-fixed with 1% osmium tetroxide in 0.1 M phosphate buffer for 2 h, and washed several times with distilled water. Samples were then en bloc stained with 1% aqueous uranyl acetate for 1 h to enhance the contrast between the cell wall layers. Specimens were dehydrated through an ascending ethanol series (50, 60, 70, 80, 90, 95, 100, and 100%; 10 min each), infiltrated with propylene oxide for 10 min, embedded in Poly/Bed 812 epoxy resin (Polysciences), and polymerized in an oven (TD-700; DOSAKA) at 70 °C for 12 h. Semithin sections (200 nm) were cut with a diamond knife (Diatome) on an ultramicrotome (EM UC7; Leica Microsystems), stained with toluidine blue, and examined by light microscopy to select the region of interest. Ultrathin sections (80 nm) were collected on copper grids (Gilder Grids), double-stained with 5% uranyl acetate (EMS) for 20 min and 3% lead citrate (Leica) for 7 min, and imaged with a transmission electron microscope (HT7800; HITACHI) at an accelerating voltage of 90 kV equipped with an RC camera. Sample preparation and imaging were performed at the Yonsei Biomedical Research Institute, Yonsei University College of Medicine. The thicknesses of the inner cell wall, outer cell wall, and capsule were measured from TEM images using ImageJ/Fiji software (n = 15 cells per strain). Statistical significance was determined by one-way ANOVA followed by Tukey’s multiple-comparison test.

### Cell wall staining

To visualize and quantify individual cell wall components, strains were cultured in liquid YPD medium at 30°C for 48 h. One milliliter of the culture was harvested and washed once with an appropriate buffer —1× PBS for CFW, WGA, and Congo Red staining, or McIlvaine’s buffer (pH 6.0; 0.2 M Na_2_HPO_4_ and 0.1 M citric acid) for Eosin Y staining, or Concanavalin A (ConA) binding buffer (10 mM HEPES, 150 mM NaCl, 1 mM MnCl_2_, 1 mM CaCl_2_, pH 7.4) for ConA staining. For the assessment of total chitin/chitosan and exposed chitooligomers, cells were incubated with 25 μg/mL of CFW (Sigma-Aldrich) or 50 μg/mL of WGA (Invitrogen), respectively, for 30 min in the dark. For detecting mannose, cells were incubated with 20 μg/mL of ConA (Invitrogen) for 15 min in the dark. Additionally, β-glucan levels were measured by staining with 50 μg/mL of Congo Red (Sigma-Aldrich) for 30 min. For chitosan-specific staining, live cells were incubated with 1.5 mg/mL of Eosin Y (Sigma-Aldrich) for 5 min in the dark (52). Following incubation, cells were washed three times with their respective buffers and mounted onto glass slides for imaging via fluorescence microscopy (ECLIPSE Ni; Nikon). Fluorescence intensity was quantified for 50 randomly selected cells per strain using ImageJ/Fiji software. Statistical significance was determined by one-way ANOVA followed by Tukey’s multiple-comparison test. Scale bars in all images represent 10 μm.

### Mitochondrial staining

WT (H99), *pro1*Δ, *pro2*Δ, and *pro3*Δ strains were cultured overnight in YPD at 30°C with shaking, inoculated into fresh YPD at an initial OD_600_ of 0.2, and grown to an OD_600_ of 0.8. Cells harvested at this point were used for the basal condition. For exposure to elevated temperature or host-mimicking conditions (HMC), cells were washed three times with PBS and resuspended in either YPD or RPMI 1640 containing sodium bicarbonate and supplemented with 10% fetal bovine serum (FBS). Cultures were incubated with shaking for 3 h at 39°C in YPD or at 37°C under 5% CO_2_ in RPMI 1640, respectively.

Following harvest, cells were stained separately in 500 µL of PBS containing 5 µM MitoSOX Red, 100 nM MitoTracker Green FM, or 1 µM nonyl acridine orange (NAO). Staining was performed at 30°C in the dark for 10 min with MitoSOX Red or 30 min with MitoTracker Green FM and NAO. After staining, cells were washed three times with PBS, resuspended in 30–50 µL of PBS, and mounted on 2% agarose pads for immediate DIC and fluorescence imaging. Mean fluorescence intensity was quantified using ImageJ and expressed in arbitrary units.

### Intracellular P5C quantification

Strains were cultured in YPD at either 30°C or 37°C for 72 h. Cells were harvested, washed once with 1× PBS, and immediately frozen in liquid nitrogen to halt metabolism. Frozen pellets were resuspended in ice-cold trichloroacetic acid (TCA), thawed slowly on ice, and lysed by bead-beating with 0.5 mm zirconia beads (10 cycles, with thorough cooling on ice between cycles). Lysates were incubated on ice for 15 min in the dark to allow complete P5C extraction and clarified by centrifugation (13,000 rpm, 10 min). Supernatants were lyophilized and resuspended in autoclaved distilled water to a final concentration of 70-fold relative to the original volume. The concentrated extract was mixed with 2-aminobenzaldehyde (2-ABZ; 25 mg/mL in 99.5% ethanol) at a 2:1 ratio, incubated in the dark, and clarified by centrifugation. Absorbance was measured at 444 nm using a microplate reader (10, 53). A TCA-buffer-only sample processed in parallel served as the negative control. Absorbance values were corrected by subtracting the blank, and expressed as fold-change relative to the WT strain. Statistical significance was determined by one-way ANOVA followed by Tukey’s multiple-comparison test using GraphPad Prism 11.

### Statistics and reproducibility

All experiments were conducted with at least three independent biological replicates. For qRT-PCR analysis, at least three biological replicates were evaluated, each containing three technical replicates. Technical replicates were averaged and were not treated as independent data points. For single-cell measurements, such as capsule thickness and cell body diameter, the specific number of cells analyzed (n) is indicated in the corresponding figure legends, and the figures present representative images and data. Data are presented as the mean ± standard error of the mean (SEM), with independent data points displayed where applicable. Statistical analyses were performed using GraphPad Prism 11. Comparisons between two groups were performed using an unpaired two-tailed t-test with Welch’s correction. Comparisons among three or more groups involving a single factor were performed using one-way ANOVA followed by Tukey’s multiple-comparison test. Data involving two factors were analyzed using two-way ANOVA followed by Tukey’s or Šídák’s multiple-comparison test, as specified in the corresponding figure legends. Multiple comparisons were performed between strains within each condition and/or between conditions within each strain, as indicated. Survival curves for the *D. melanogaster* and murine infection models were analyzed using the log-rank (Mantel–Cox) test.

## Data availability

All data are available in the manuscript and the Supporting Information files. Other supporting raw data files were uploaded to figshare as follows: https://doi.org/10.6084/m9.figshare.32676075

## ACKNOWLEDGEMENTS

This work was supported by grants from the National Research Foundation of Korea (NRF) funded by the Korea government (MSIT) (RS-2025-18362970, RS-2025-02215093, and RS-2025-00555365 to Y.-S.B.; RS-2022-NR072215 and RS-2021-NF000550 to K.-T.L.); by a grant from the Korea Health Technology R&D Project through the Korea Health Industry Development Institute (KHIDI), funded by the Ministry of Health & Welfare, Republic of Korea (RS-2026-25518347 to Y.-S.B.); and by the Korea Institute for Advancement of Technology (KIAT), funded by the Ministry of Trade, Industry and Energy in 2026 (RS-2024-00418203 to Y.-S.B.).

## AUTHOR CONTRIBUTIONS

Y.L.: Data curation, Conceptualization, Investigation, Methodology, Formal analysis, Validation, Visualization, Writing – original draft, Writing – review & editing. M.L.: Investigation, Methodology. S.K.: Investigation, Methodology, Formal analysis, Validation. H.C.: Investigation, Methodology, Formal analysis, Validation. D.W.: Investigation, Methodology, Formal analysis, Validation. K.-A.L.: Investigation, Methodology, Formal analysis, Validation. W.-J.L.: Resources, Supervision, Writing – original draft, Writing – review & editing. K.-T.L.: Conceptualization, Supervision, Project administration, Resources, Funding acquisition, Writing – original draft, Writing – review & editing. Y.-S.B.: Conceptualization, Supervision, Project administration, Resources, Funding acquisition, Writing – original draft, Writing – review & editing.

## REFERENCES

1. Brown AJP, Cowen LE, di Pietro A, Quinn J. 2017. Stress adaptation. Microbiol Spectr 5. doi:10.1128/microbiolspec.FUNK-0048-2016.

2. Jung KW, Lee SH, Lee KT, Bahn YS. 2024. Sensing and responding to host-derived stress signals: lessons from fungal meningitis pathogen. Curr Opin Microbiol 80:102514. doi:10.1016/j.mib.2024.102514.

3. Garbe E, Vylkova S. 2019. Role of amino acid metabolism in the virulence of human pathogenic fungi. Curr Clin Microbiol Rep 6:108–119. doi:10.1007/s40588-019-00124-5.

4. Alcazar-Fuoli L. 2016. Amino acid biosynthetic pathways as antifungal targets for fungal infections. Virulence 7:376–378. doi:10.1080/21505594.2016.1169360.

5. Chen C, Dickman MB. 2005. Proline suppresses apoptosis in the fungal pathogen *Colletotrichum trifolii*. Proc Natl Acad Sci U S A 102:3459–3464. doi:10.1073/pnas.0407960102.

6. Takagi H. 2008. Proline as a stress protectant in yeast: physiological functions, metabolic regulations, and biotechnological applications. Appl Microbiol Biotechnol 81:211–223. doi:10.1007/s00253-008-1698-5.

7. Tomenchok DM, Brandriss MC. 1987. Gene-enzyme relationships in the proline biosynthetic pathway of *Saccharomyces cerevisiae*. J Bacteriol 169:5364–5372. doi:10.1128/jb.169.12.5364-5372.1987.

8. Davis RH. 1986. Compartmental and regulatory mechanisms in the arginine pathways of *Neurospora crassa* and *Saccharomyces cerevisiae*. Microbiol Rev 50:280–313. doi:10.1128/mr.50.3.280-313.1986.

9. Ljungdahl PO, Daignan-Fornier B. 2012. Regulation of amino acid, nucleotide, and phosphate metabolism in *Saccharomyces cerevisiae*. Genetics 190:885–929. doi:10.1534/genetics.111.133306.

10. Silao FGS, Jiang T, Bereczky-Veress B, Kuhbacher A, Ryman K, Uwamohoro N, Jenull S, Nogueira F, Ward M, Lion T, Urban CF, Rupp S, Kuchler K, Chen C, Peuckert C, Ljungdahl PO. 2023. Proline catabolism is a key factor facilitating *Candida albicans* pathogenicity. PLoS Pathog 19:e1011677. doi:10.1371/journal.ppat.1011677.

11. Corte Z, Venta R. 2010. Biological variation of free plasma amino acids in healthy individuals. Clin Chem Lab Med 48:99–104. doi:10.1515/CCLM.2010.008.

12. May RC, Stone NR, Wiesner DL, Bicanic T, Nielsen K. 2016. *Cryptococcus*: from environmental saprophyte to global pathogen. Nat Rev Microbiol 14:106–117. doi:10.1038/nrmicro.2015.6.

13. Kronstad JW, Attarian R, Cadieux B, Choi J, D’Souza CA, Griffiths EJ, Geddes JM, Hu G, Jung WH, Kretschmer M, Saikia S, Wang J. 2011. Expanding fungal pathogenesis: *Cryptococcus* breaks out of the opportunistic box. Nat Rev Microbiol 9:193–203. doi:10.1038/nrmicro2522.

14. Chen Y, Shi ZW, Strickland AB, Shi M. 2022. *Cryptococcus neoformans* infection in the central nervous system: the battle between host and pathogen. J Fungi (Basel) 8:1069. doi:10.3390/jof8101069.

15. Lee IR, Lui EY, Chow EW, Arras SD, Morrow CA, Fraser JA. 2013. Reactive oxygen species homeostasis and virulence of the fungal pathogen *Cryptococcus neoformans* requires an intact proline catabolism pathway. Genetics 194:421–433. doi:10.1534/genetics.113.150326.

16. Idnurm A, Walton FJ, Floyd A, Reedy JL, Heitman J. 2009. Identification of *ENA1* as a virulence gene of the human pathogenic fungus *Cryptococcus neoformans* through signature-tagged insertional mutagenesis. Eukaryot Cell 8:315–326. doi:10.1128/EC.00375-08.

17. Lee KT, Hong J, Lee DG, Lee M, Cha S, Lim YG, Jung KW, Hwangbo A, Lee Y, Yu SJ, Chen YL, Lee JS, Cheong E, Bahn YS. 2020. Fungal kinases and transcription factors regulating brain infection in *Cryptococcus neoformans*. Nat Commun 11:1521. doi:10.1038/s41467-020-15329-2.

18. Yao Z, Zou C, Zhou H, Wang J, Lu L, Li Y, Chen B. 2013. Δ^1^-pyrroline-5-carboxylate/glutamate biogenesis is required for fungal virulence and sporulation. PLoS One 8:e73483. doi:10.1371/journal.pone.0073483.

19. Lee KT, So YS, Yang DH, Jung KW, Choi J, Lee DG, Kwon H, Jang J, Wang LL, Cha S, Meyers GL, Jeong E, Jin JH, Lee Y, Hong J, Bang S, Ji JH, Park G, Byun HJ, Park SW, Park YM, Adedoyin G, Kim T, Averette AF, Choi JS, Heitman J, Cheong E, Lee YH, Bahn YS. 2016. Systematic functional analysis of kinases in the fungal pathogen *Cryptococcus neoformans*. Nat Commun 7:12766. doi:10.1038/ncomms12766.

20. Kozel TR, Gotschlich EC. 1982. The capsule of *Cryptococcus neoformans* passively inhibits phagocytosis of the yeast by macrophages. J Immunol 129:1675–1680.

21. O’Meara TR, Alspaugh JA. 2012. The *Cryptococcus neoformans* capsule: a sword and a shield. Clin Microbiol Rev 25:387–408. doi:10.1128/CMR.00001-12.

22. Jang EH, Kim JS, Yu SR, Bahn YS. 2022. Unraveling capsule biosynthesis and signaling networks in *Cryptococcus neoformans*. Microbiol Spectr 10:e0286622. doi:10.1128/spectrum.02866-22.

23. Wang Y, Aisen P, Casadevall A. 1995. *Cryptococcus neoformans* melanin and virulence: mechanism of action. Infect Immun 63:3131–3136. doi:10.1128/iai.63.8.3131-3136.1995.

24. Lee D, Jang EH, Lee M, Kim SW, Lee Y, Lee KT, Bahn YS. 2019. Unraveling melanin biosynthesis and signaling networks in *Cryptococcus neoformans*. mBio 10:e02267–19. doi:10.1128/mBio.02267-19.

25. Lin X, Heitman J. 2006. The biology of the *Cryptococcus neoformans* species complex. Annu Rev Microbiol 60:69–105. doi:10.1146/annurev.micro.60.080805.142102.

26. Lin X, Hull CM, Heitman J. 2005. Sexual reproduction between partners of the same mating type in *Cryptococcus neoformans*. Nature 434:1017–1021. doi:10.1038/nature03448.

27. Mpamhanga CD, Kounatidis I. 2024. The utility of *Drosophila melanogaster* as a fungal infection model. Front Immunol 15:1349027. doi:10.3389/fimmu.2024.1349027.

28. Martho KF, de Melo AT, Takahashi JP, Guerra JM, Santos DC, Purisco SU, Melhem MS, Fazioli RD, Phanord C, Sartorelli P, Vallim MA, Pascon RC. 2016. Amino acid permeases and virulence in *Cryptococcus neoformans*. PLoS One 11:e0163919. doi:10.1371/journal.pone.0163919.

29. Liu N, Dai Z, Zhang Y, Chen J, Yang Y, Wu G, Tso P, Wu Z. 2019. Maternal L-proline supplementation enhances fetal survival, placental development, and nutrient transport in mice. Biol Reprod 100:1073–1081. doi:10.1093/biolre/ioy240.

30. Deuschle K, Funck D, Forlani G, Stransky H, Biehl A, Leister D, van der Graaff E, Kunze R, Frommer WB. 2004. The role of Δ^1^-pyrroline-5-carboxylate dehydrogenase in proline degradation. Plant Cell 16:3413–3425. doi:10.1105/tpc.104.023622.

31. Fichman Y, Gerdes SY, Kovacs H, Szabados L, Zilberstein A, Csonka LN. 2015. Evolution of proline biosynthesis: enzymology, bioinformatics, genetics, and transcriptional regulation. Biol Rev Camb Philos Soc 90:1065–1099. doi:10.1111/brv.12146.

32. Sekine T, Kawaguchi A, Hamano Y, Takagi H. 2007. Desensitization of feedback inhibition of the *Saccharomyces cerevisiae* gamma-glutamyl kinase enhances proline accumulation and freezing tolerance. Appl Environ Microbiol 73:4011–4019. doi:10.1128/AEM.00730-07.

33. Zhu X, Gibbons J, Garcia-Rivera J, Casadevall A, Williamson PR. 2001. Laccase of *Cryptococcus neoformans* is a cell wall-associated virulence factor. Infect Immun 69:5589–5596. doi:10.1128/IAI.69.9.5589-5596.2001.

34. Park YD, Chen SH, Camacho E, Casadevall A, Williamson PR. 2020. Role of the ESCRT Pathway in laccase trafficking and virulence of *Cryptococcus neoformans*. Infect Immun 88:e00954–19. doi:10.1128/IAI.00954-19.

35. Rodrigues ML, Nakayasu ES, Oliveira DL, Nimrichter L, Nosanchuk JD, Almeida IC, Casadevall A. 2008. Extracellular vesicles produced by *Cryptococcus neoformans* contain protein components associated with virulence. Eukaryot Cell 7:58–67. doi:10.1128/EC.00370-07.

36. Littman ML. 1958. Capsule synthesis by *Cryptococcus neoformans*. Trans N Y Acad Sci 20:623–648. doi:10.1111/j.2164-0947.1958.tb00625.x.

37. Lee DY, Yun SH, Lee SY, Lee J, Mariano EJ, Joo ST, Choi I, Seok Choi J, Kim GD, Lee J, Choi SH, Jin Hur S. 2024. Corrigendum to “Analysis of commercial fetal bovine serum (FBS) and its substitutes in the development of cultured meat” [Food Res. Int.174P1 (2023) 113617]. Food Res Int 188:114303. doi:10.1016/j.foodres.2024.114303.

38. Lee DY, Yun SH, Lee SY, Lee J, Jr Mariano E, Joo ST, Choi I, Choi JS, Kim GD, Lee J, Choi SH, Hur SJ. 2023. Analysis of commercial fetal bovine serum (FBS) and its substitutes in the development of cultured meat. Food Res Int 174:113617. doi:10.1016/j.foodres.2023.113617.

39. Kingsbury JM, Yang Z, Ganous TM, Cox GM, McCusker JH. 2004. Novel chimeric spermidine synthase-saccharopine dehydrogenase gene (*SPE3*-*LYS9*) in the human pathogen *Cryptococcus neoformans*. Eukaryot Cell 3:752–763. doi:10.1128/EC.3.3.752-763.2004.

40. Kaur J, Goldsmith J, Tankka A, Bustamante Eguiguren S, Gimenez AA, Vick L, Debnath J, Vlahakis A. 2021. Atg32-dependent mitophagy sustains spermidine and nitric oxide required for heat-stress tolerance in *Saccharomyces cerevisiae*. J Cell Sci 134:jcs253781. doi:10.1242/jcs.253781.

41. Kruger A, Vowinckel J, Mulleder M, Grote P, Capuano F, Bluemlein K, Ralser M. 2013. Tpo1-mediated spermine and spermidine export controls cell cycle delay and times antioxidant protein expression during the oxidative stress response. EMBO Rep 14:1113–1119. doi:10.1038/embor.2013.165.

42. Arras SDM, Ormerod KL, Erpf PE, Espinosa MI, Carpenter AC, Blundell RD, Stowasser SR, Schulz BL, Tanurdzic M, Fraser JA. 2017. Convergent microevolution of *Cryptococcus neoformans* hypervirulence in the laboratory and the clinic. Sci Rep 7:17918. doi:10.1038/s41598-017-18106-2.

43. Semighini CP, Averette AF, Perfect JR, Heitman J. 2011. Deletion of *Cryptococcus neoformans* AIF ortholog promotes chromosome aneuploidy and fluconazole-resistance in a metacaspase-independent manner. PLoS Pathog 7:e1002364. doi:10.1371/journal.ppat.1002364.

44. Kim MS, Kim SY, Jung KW, Bahn YS. 2012. Targeted gene disruption in *Cryptococcus neoformans* using double-joint PCR with split dominant selectable markers. Methods Mol Biol 845:67–84. doi:10.1007/978-1-61779-539-8_5.

45. Jung KW, Lee KT, So YS, Bahn YS. 2018. Genetic manipulation of *Cryptococcus neoformans*. Curr Protoc Microbiol 50:e59. doi:10.1002/cpmc.59.

46. Tamura K, Stecher G, Kumar S. 2021. MEGA11: Molecular Evolutionary Genetics Analysis Version 11. Mol Biol Evol 38:3022–3027. doi:10.1093/molbev/msab120.

47. Jones DT, Taylor WR, Thornton JM. 1992. The rapid generation of mutation data matrices from protein sequences. Comput Appl Biosci 8:275–282. doi:10.1093/bioinformatics/8.3.275.

48. Abramson J, Adler J, Dunger J, Evans R, Green T, Pritzel A, Ronneberger O, Willmore L, Ballard AJ, Bambrick J, Bodenstein SW, Evans DA, Hung CC, O’Neill M, Reiman D, Tunyasuvunakool K, Wu Z, Zemgulyte A, Arvaniti E, Beattie C, Bertolli O, Bridgland A, Cherepanov A, Congreve M, Cowen-Rivers AI, Cowie A, Figurnov M, Fuchs FB, Gladman H, Jain R, Khan YA, Low CMR, Perlin K, Potapenko A, Savy P, Singh S, Stecula A, Thillaisundaram A, Tong C, Yakneen S, Zhong ED, Zielinski M, Zidek A, Bapst V, Kohli P, Jaderberg M, Hassabis D, Jumper JM. 2024. Accurate structure prediction of biomolecular interactions with AlphaFold 3. Nature 630:493–500. doi:10.1038/s41586-024-07487-w.

49. Jung WH, Sham A, White R, Kronstad JW. 2006. Iron regulation of the major virulence factors in the AIDS-associated pathogen *Cryptococcus neoformans*. PLoS Biol 4:e410. doi:10.1371/journal.pbio.0040410.

50. Livak KJ, Schmittgen TD. 2001. Analysis of relative gene expression data using real-time quantitative PCR and the 2^-ΔΔCT^ method. Methods 25:402–408. doi:10.1006/meth.2001.1262.

51. Casadevall A, Cleare W, Feldmesser M, Glatman-Freedman A, Goldman DL, Kozel TR, Lendvai N, Mukherjee J, Pirofski LA, Rivera J, Rosas AL, Scharff MD, Valadon P, Westin K, Zhong Z. 1998. Characterization of a murine monoclonal antibody to *Cryptococcus neoformans* polysaccharide that is a candidate for human therapeutic studies. Antimicrob Agents Chemother 42:1437–1446. doi:10.1128/AAC.42.6.1437.

52. Baker LG, Specht CA, Donlin MJ, Lodge JK. 2007. Chitosan, the deacetylated form of chitin, is necessary for cell wall integrity in *Cryptococcus neoformans*. Eukaryot Cell 6:855–867. doi:10.1128/EC.00399-06.

53. Mezl VA, Knox WE. 1976. Properties and analysis of a stable derivative of pyrroline-5-carboxylic acid for use in metabolic studies. Anal Biochem 74:430–440. doi:10.1016/0003-2697(76)90223-2.

