## Supplementary Information for "A convergent proline biosynthetic step couples cell wall integrity to virulence in Cryptococcus neoformans"

**Figure S1 (Lee et al.)**

**A**

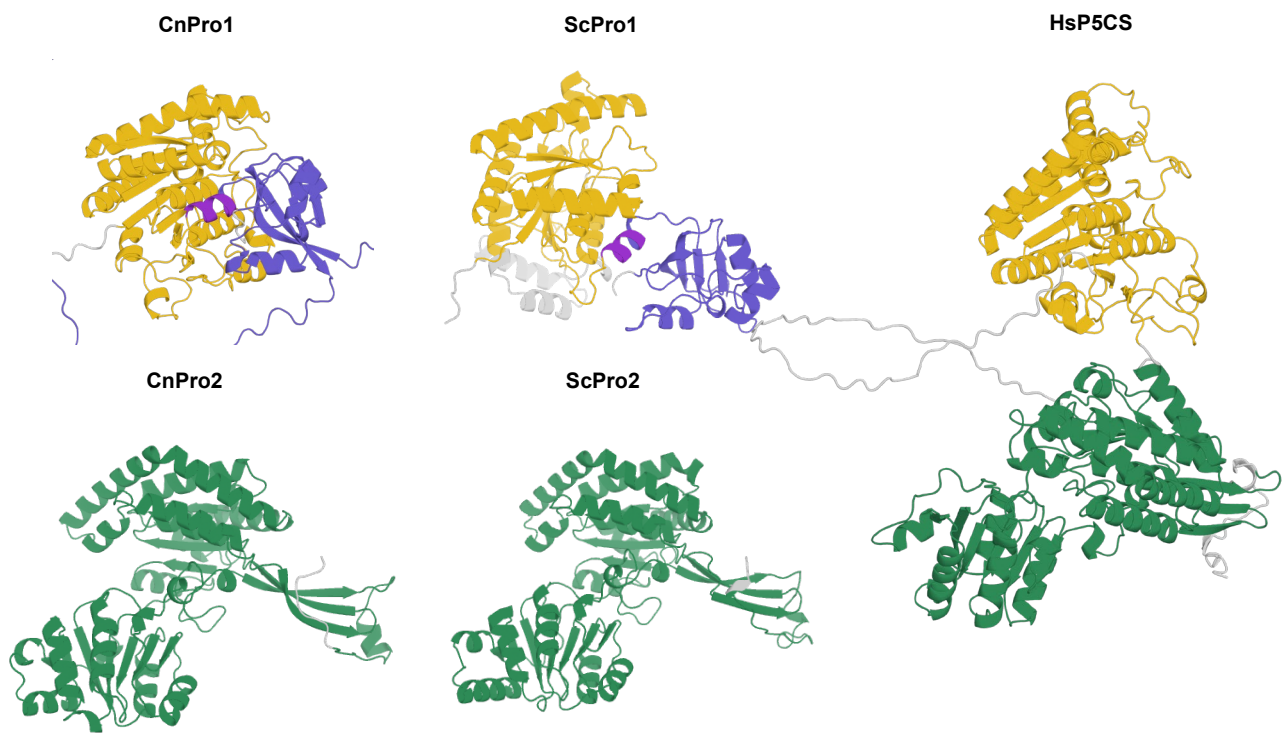

**B**

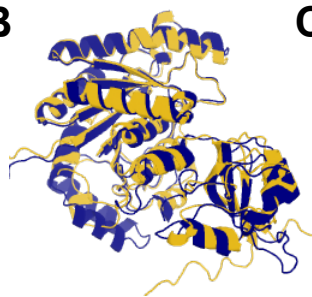

CnPro1: yellow
ScPro1: navy

**C**

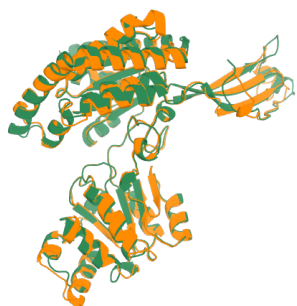

CnPro2: green
ScPro2: orange

**D**

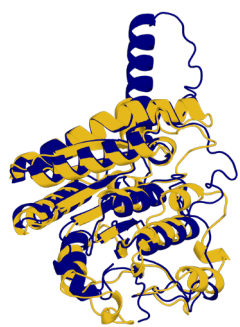

Cn AGK-like domain: yellow
Hs AGK-like domain: navy

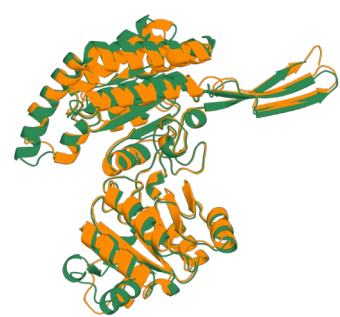

Cn ALDH-like domain: green
Hs ALDH-like domain: orange

(Continued)

**Figure S1 (Lee et al.)**

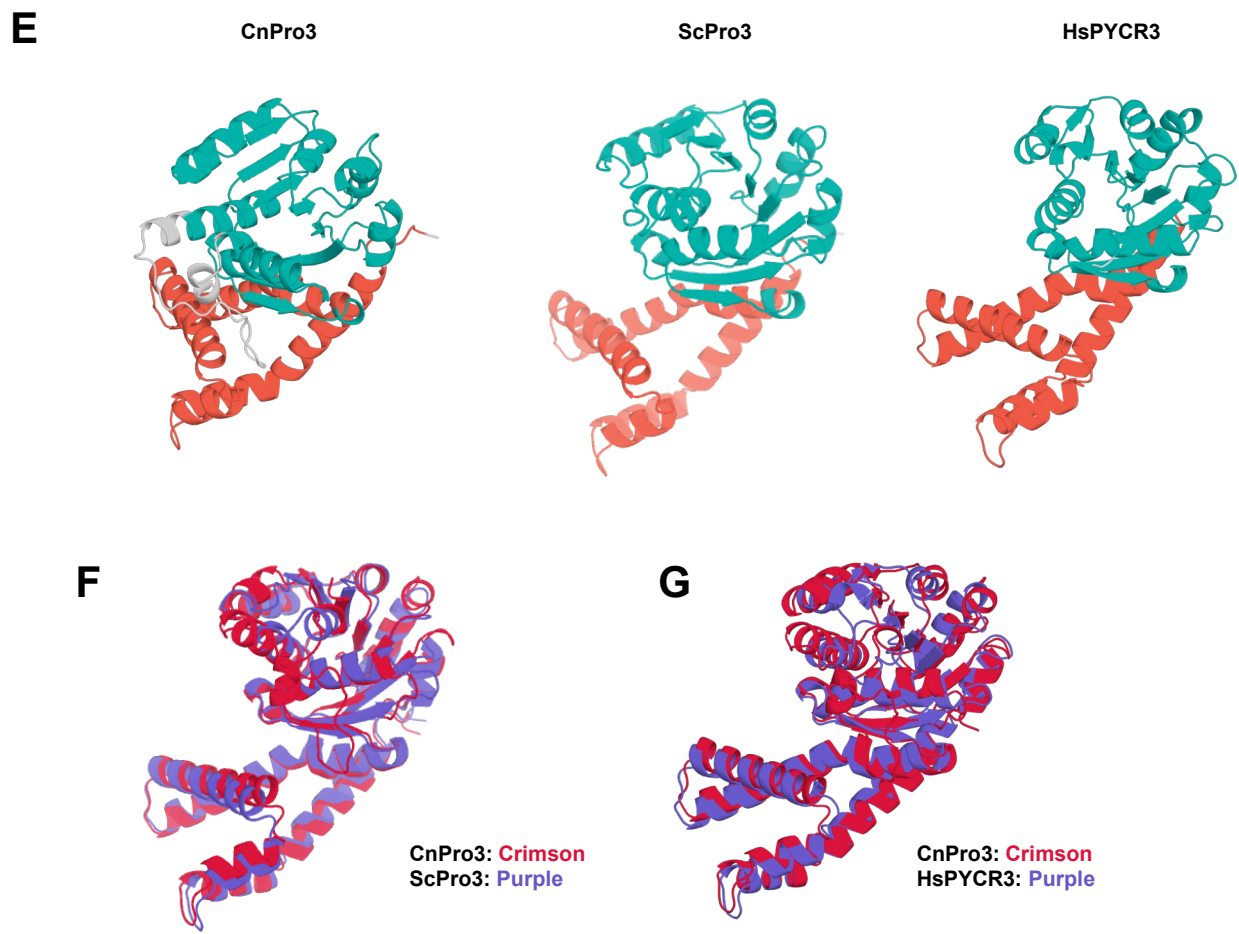

**Figure S1. Structural characterization of the proline biosynthetic enzymes in *Cryptococcus neoformans***

(A) Predicted structures of CnPro1, ScPro1, CnPro2, ScPro2, and human P5CS generated by AlphaFold3. The N-terminal acetylglutamate kinase-like domain is shown in yellow, the γ-glutamyl phosphate reductase (ALDH-like) domain in green, and the PUA\_G5K domain in light purple, and the overlap region of ALDH-like and PUA\_G5K domain in deep purple. (B) Structural alignment of CnPro1 (yellow) and ScPro1 (navy). Predicted by AlphaFold3 and aligned in PyMOL. (C) Structural alignment of CnPro2 (green) and ScPro2 (orange). Predicted by AlphaFold3 and aligned in PyMOL. (D) Structural alignment of *C. neoformans* Pro1 (AGK-like, yellow) and Pro2 (ALDH-like, green) with HsP5CS domains (AGK-like, navy; ALDH-like, orange). Predicted by AlphaFold3 and aligned in PyMOL. (E) Predicted structures of CnPro3, ScPro3, and human PYCR3 generated by AlphaFold3. The NAD(P)-binding Rossmann-like domain is shown in cyan, and the C-terminal ProC-like domain in red. (F) Structural alignment of *C. neoformans* Pro3 (crimson) with ScPro3 (purple). Predicted by AlphaFold3 and aligned in PyMOL. (G) Structural alignment of CnPro3 (crimson) with HsPYCR3 (purple). Predicted by AlphaFold3 and aligned in PyMOL.

**Figure S2 (Lee et al.)**

**A**

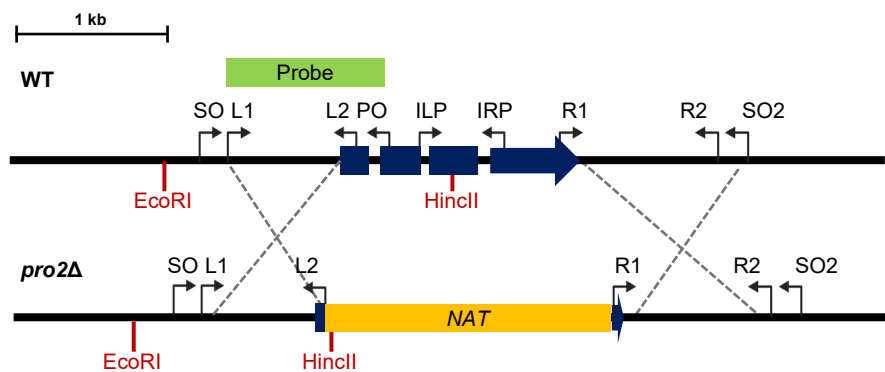

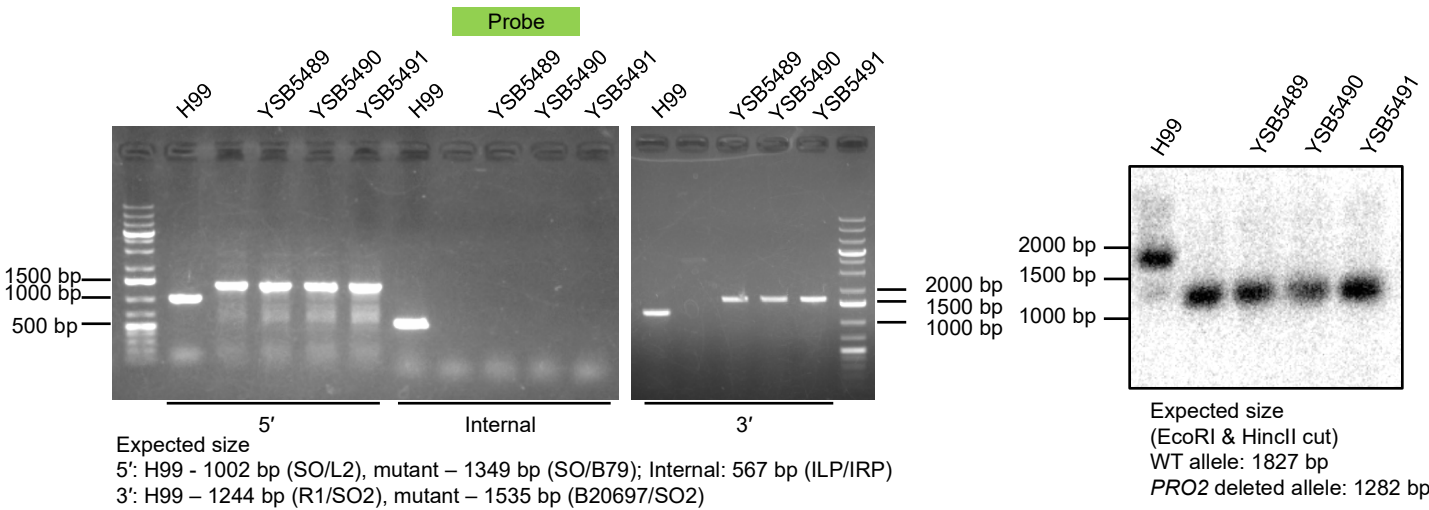

**B**

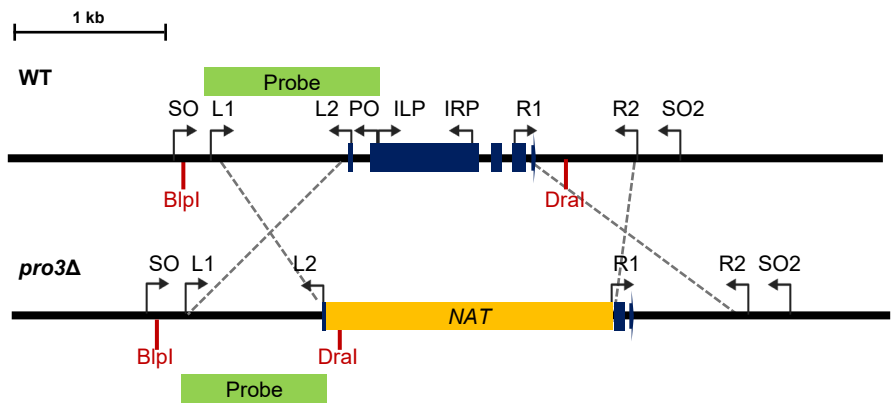

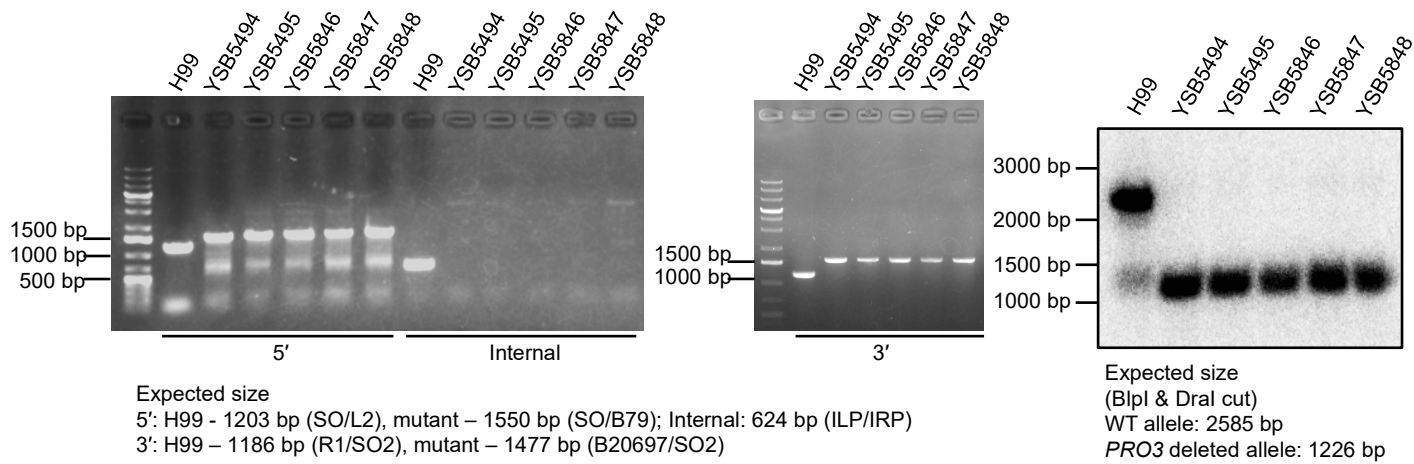

(Continued)

**Figure S2 (Lee et al.)**

**C**

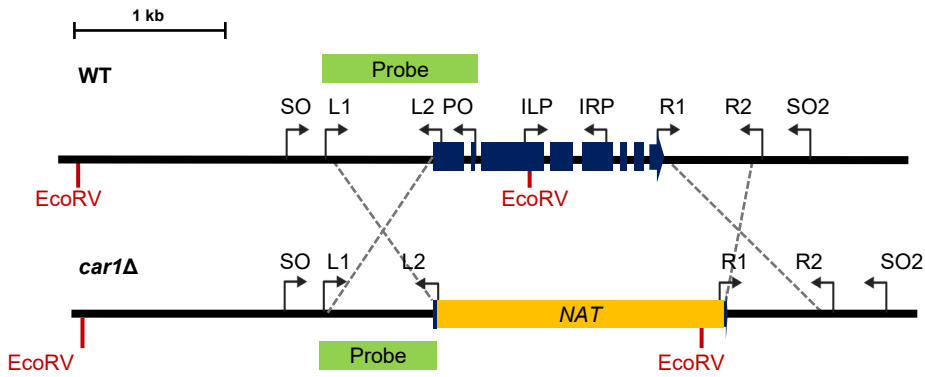

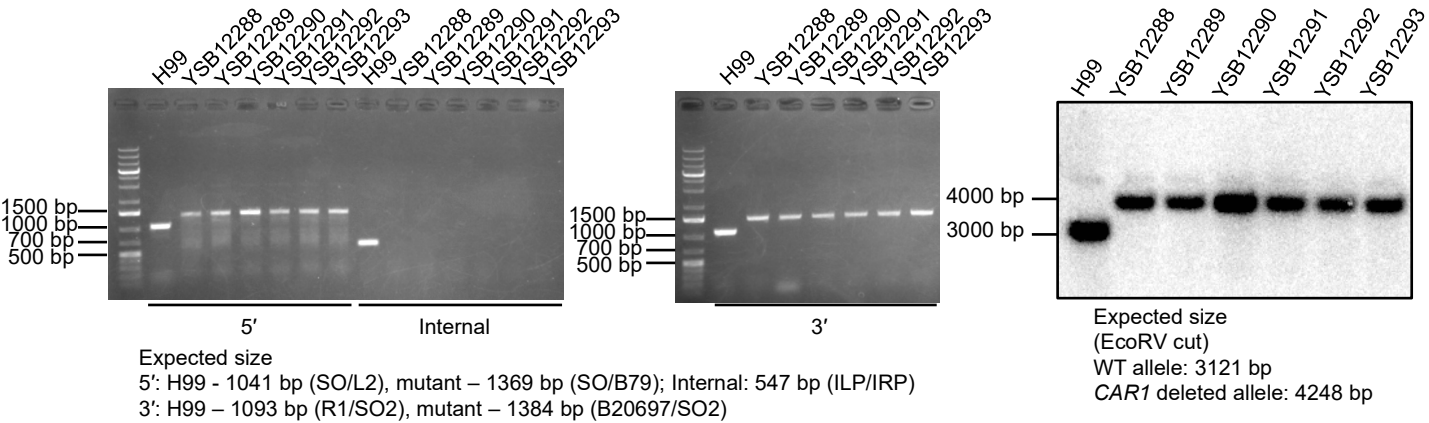

**D**

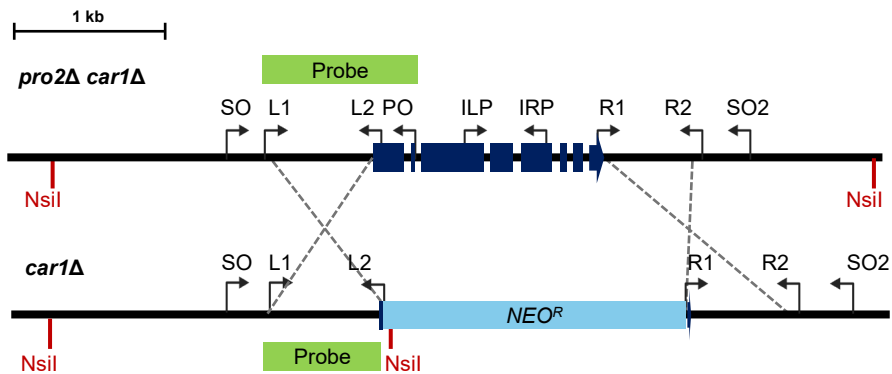

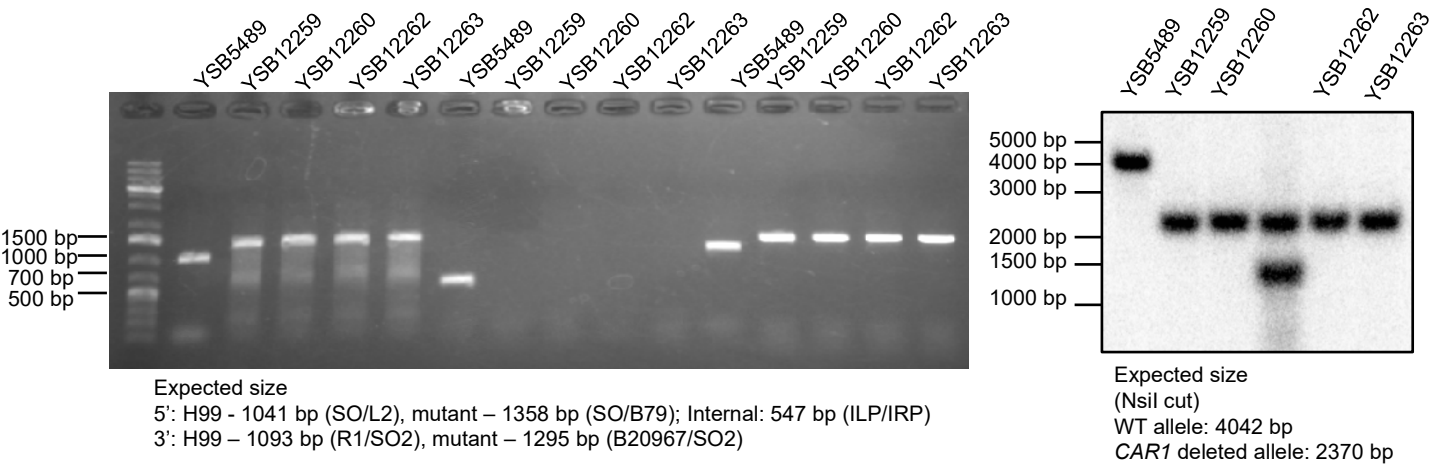

(Continued)

### Figure S2 (Lee et al.)

E

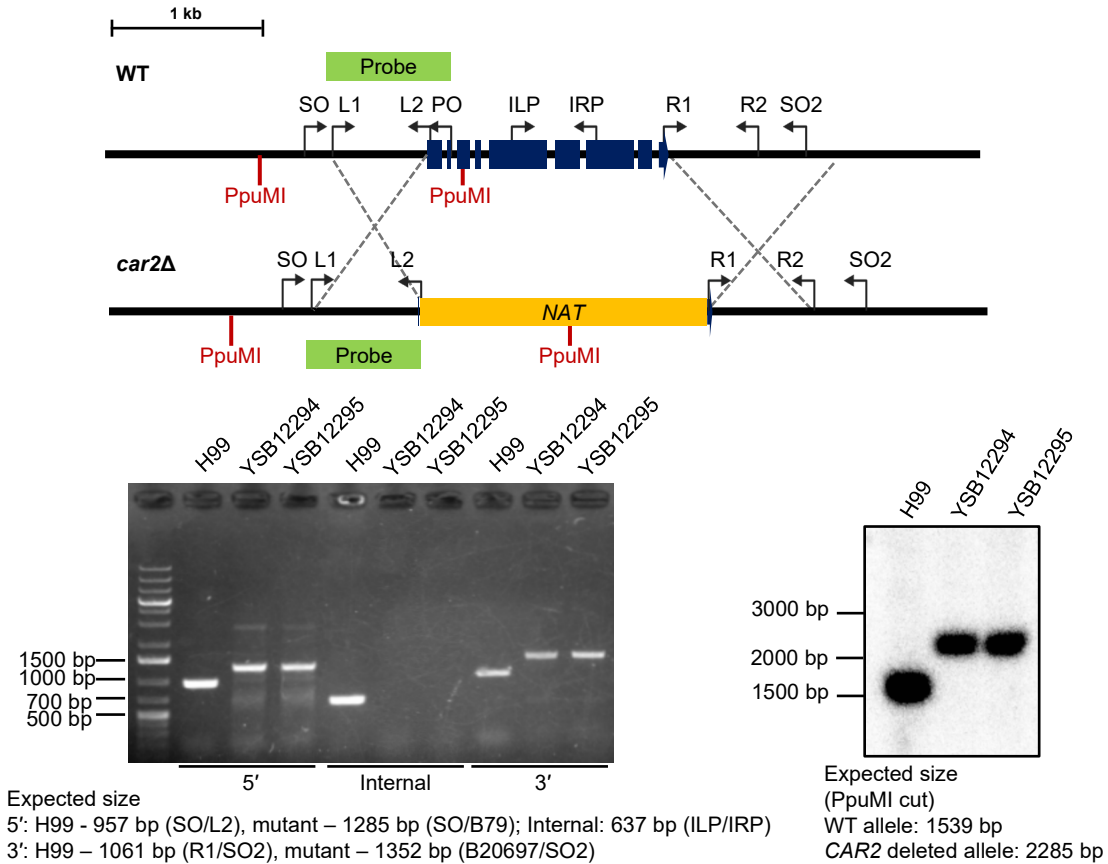

F

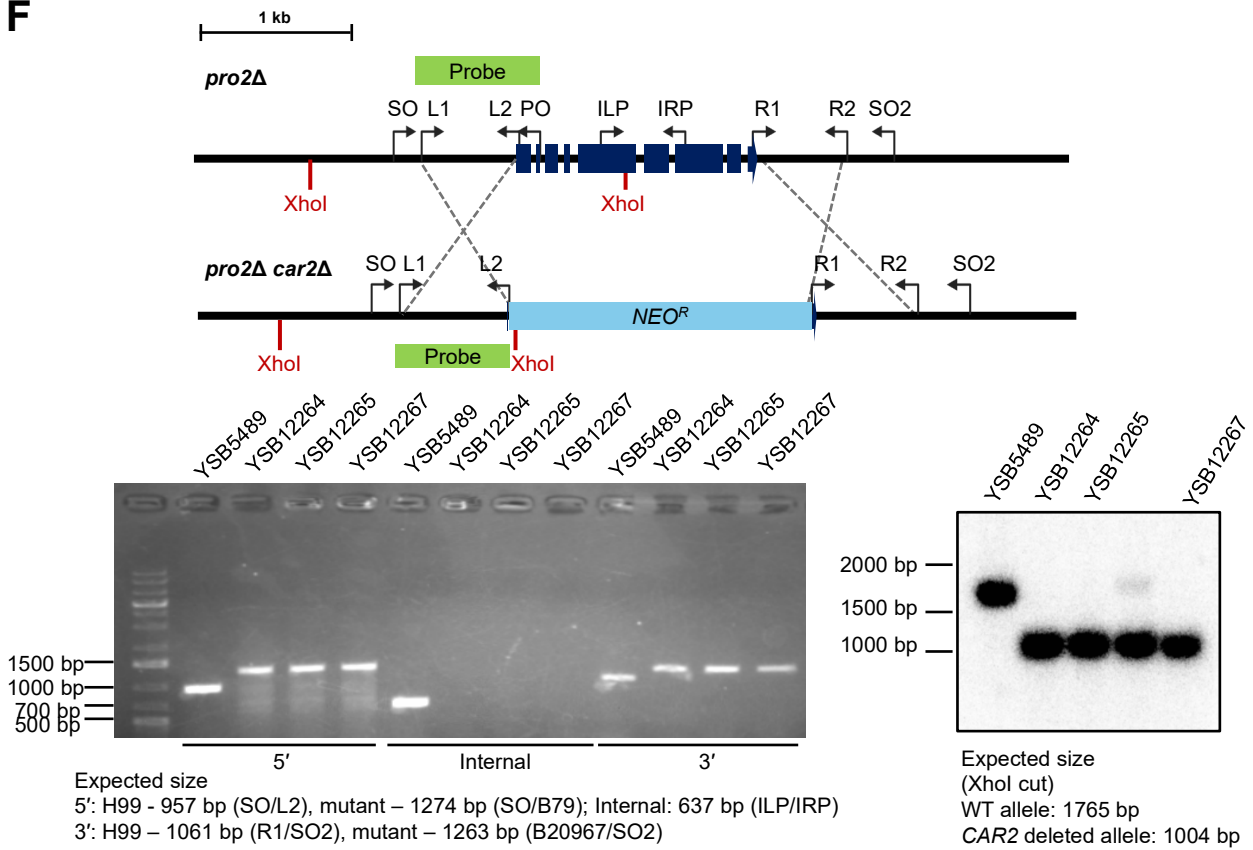

(Continued)

**Figure S2 (Lee et al.)**

**G**

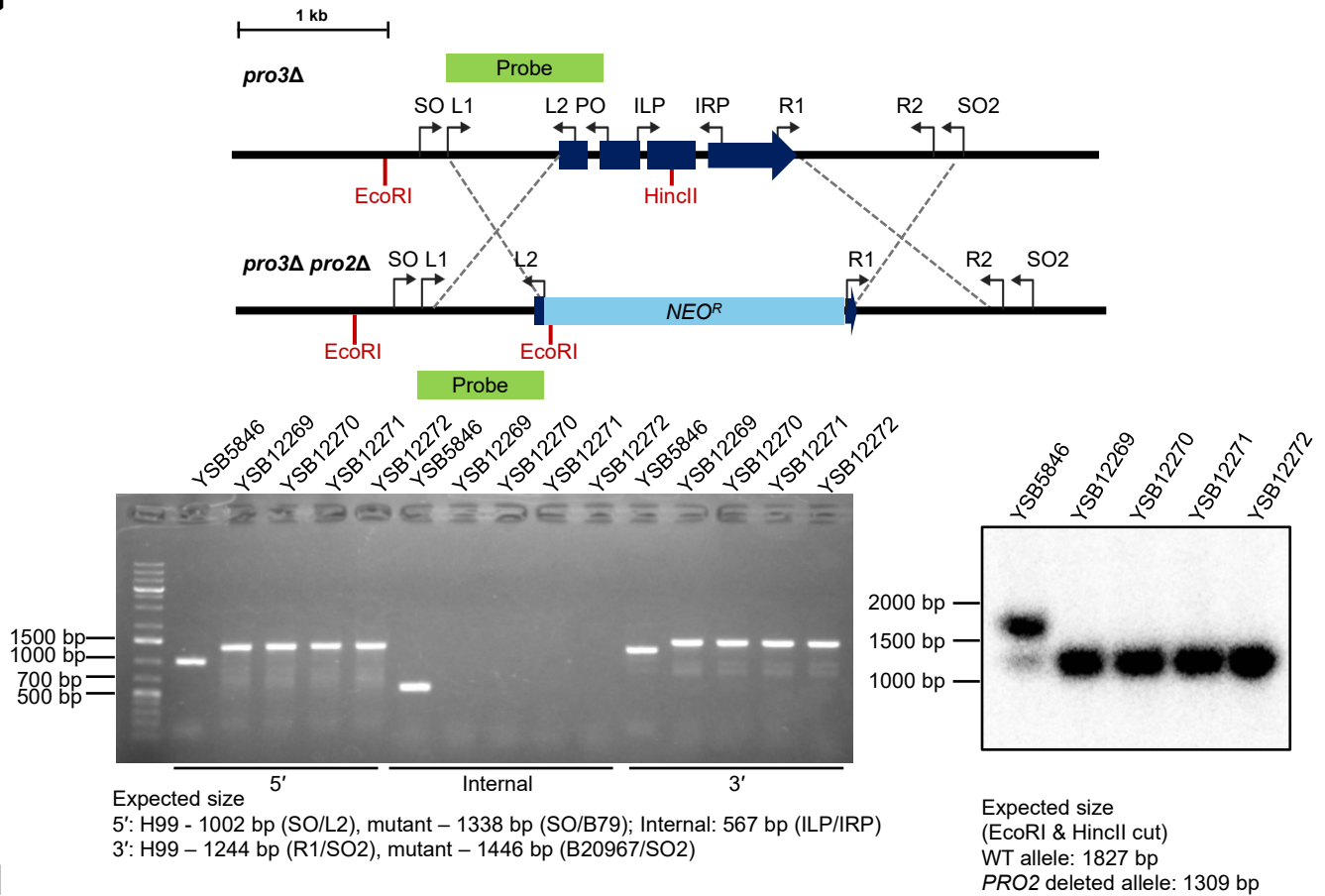

**H**

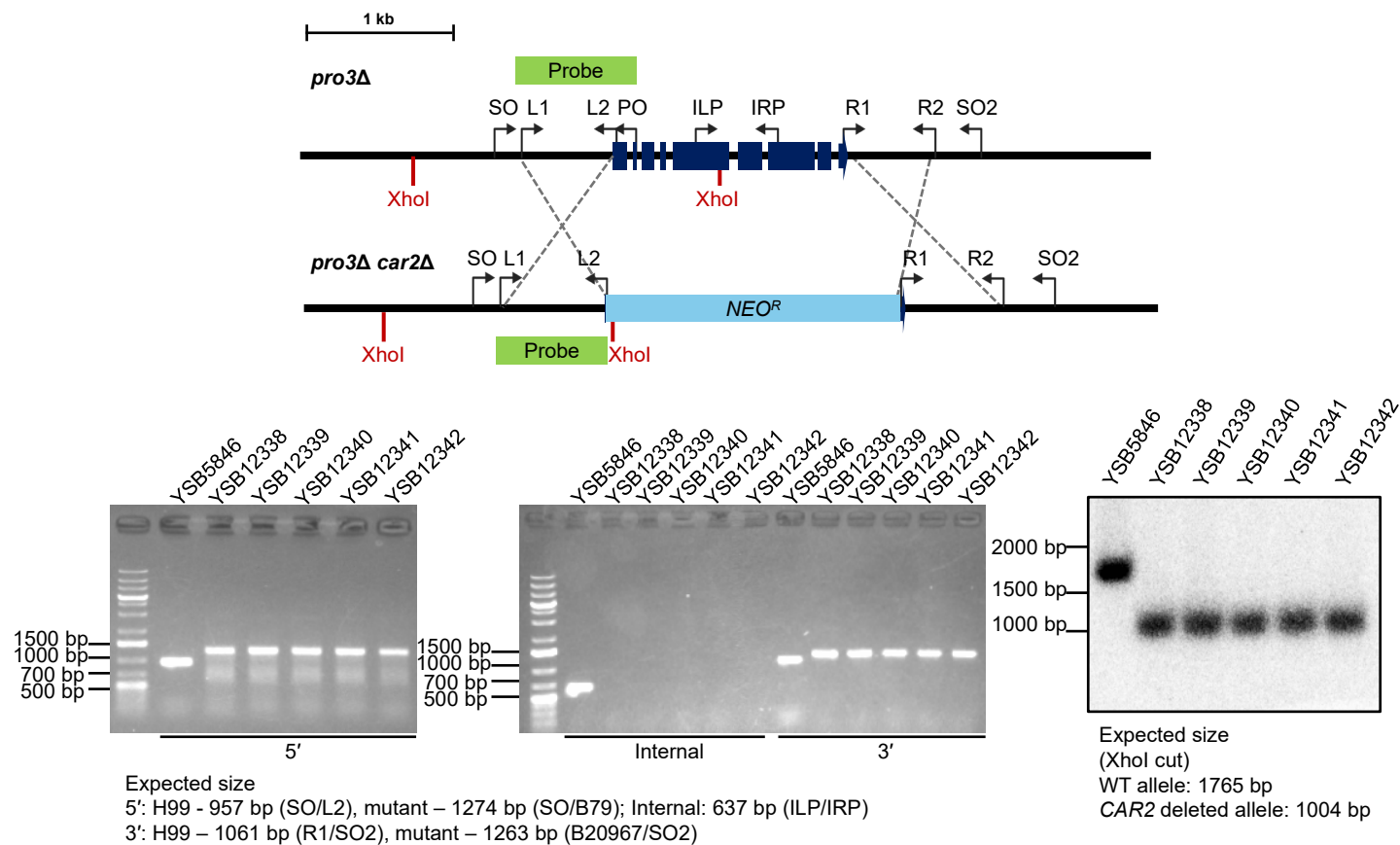

(Continued)

**Figure S2 (Lee et al.)**

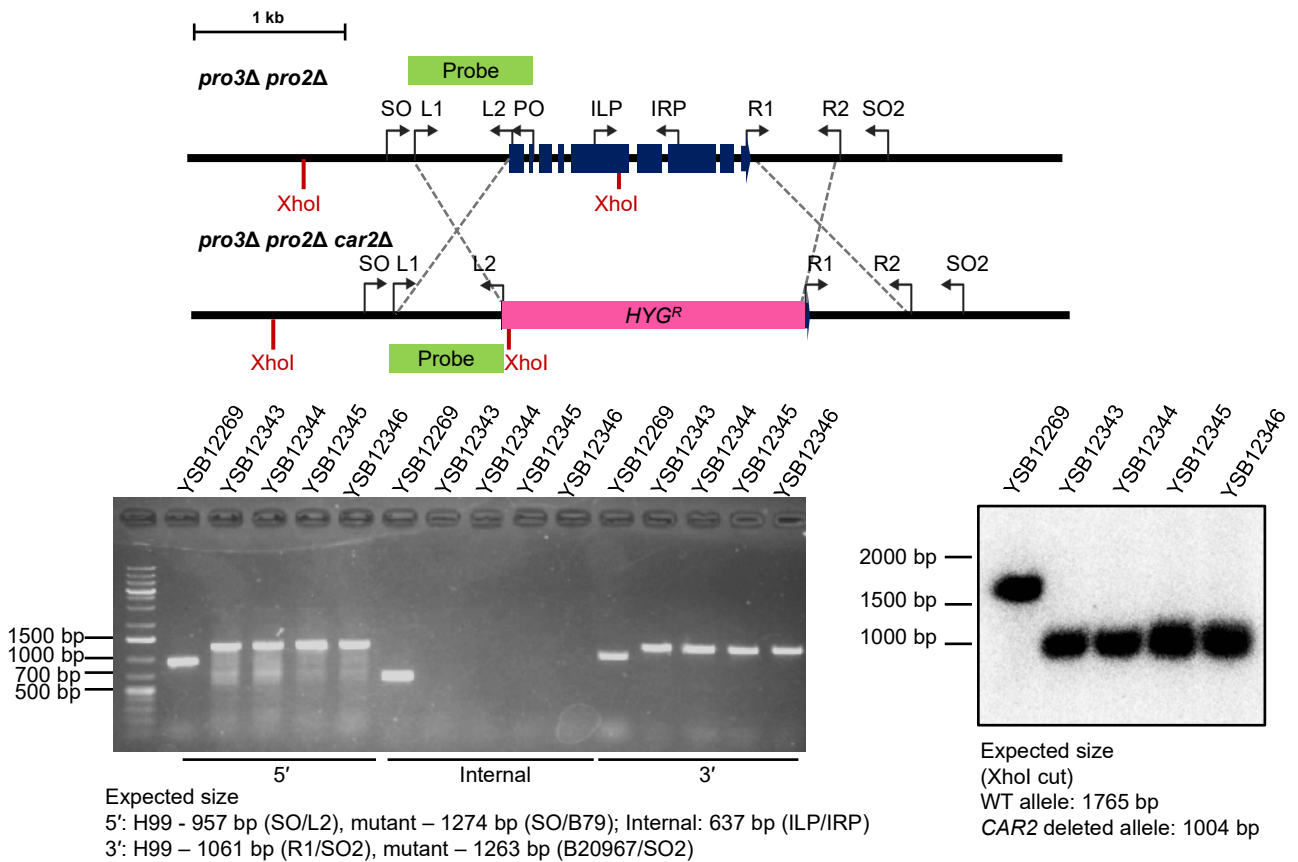

(Continued)

**Figure S2 (Lee et al.)**

**J**

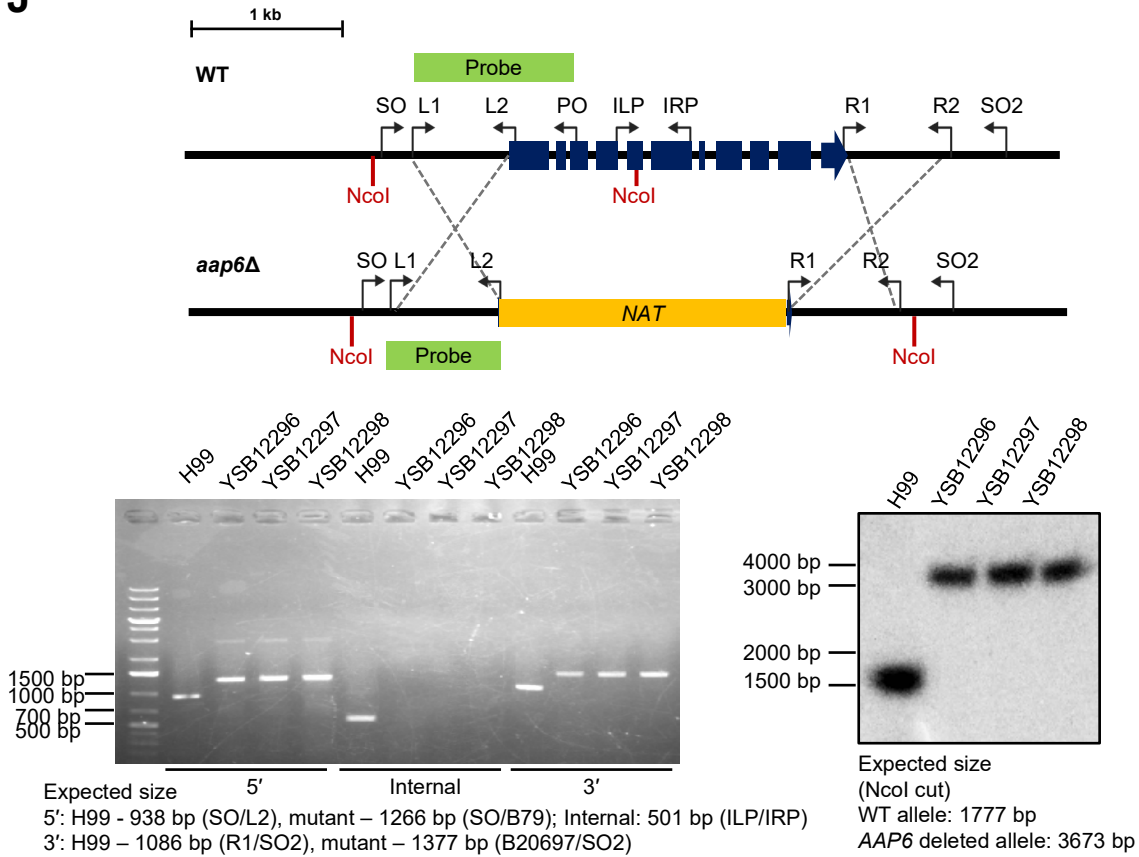

**K**

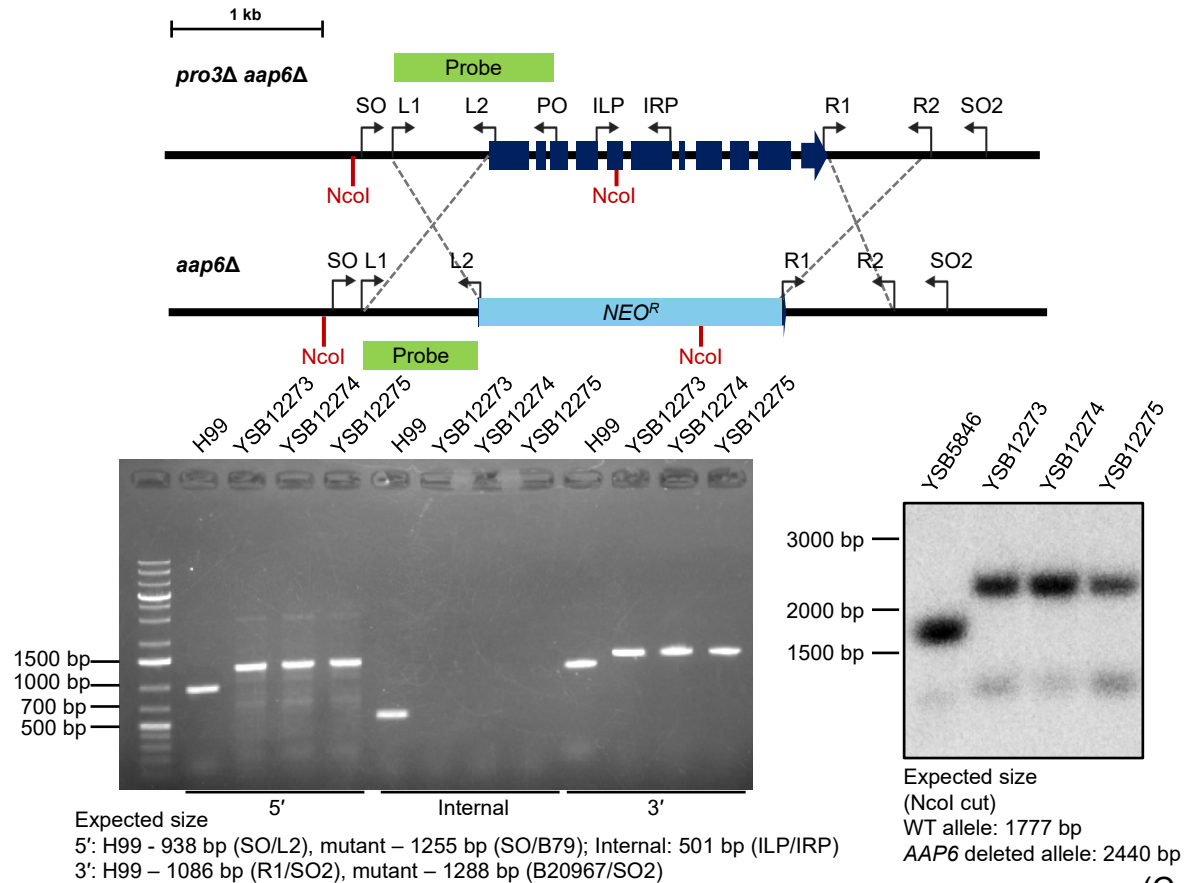

(Continued)

**Figure S2 (Lee et al.)**

**L**

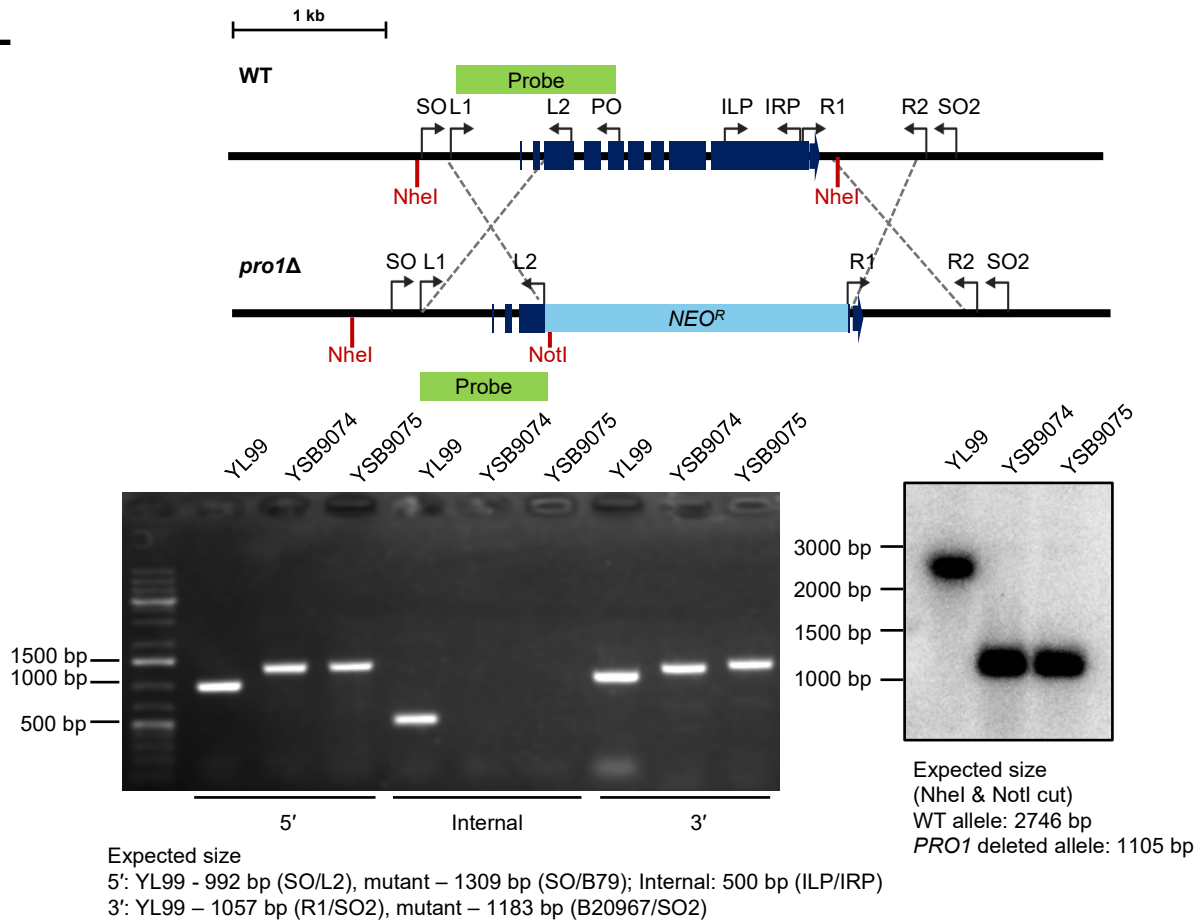

**M**

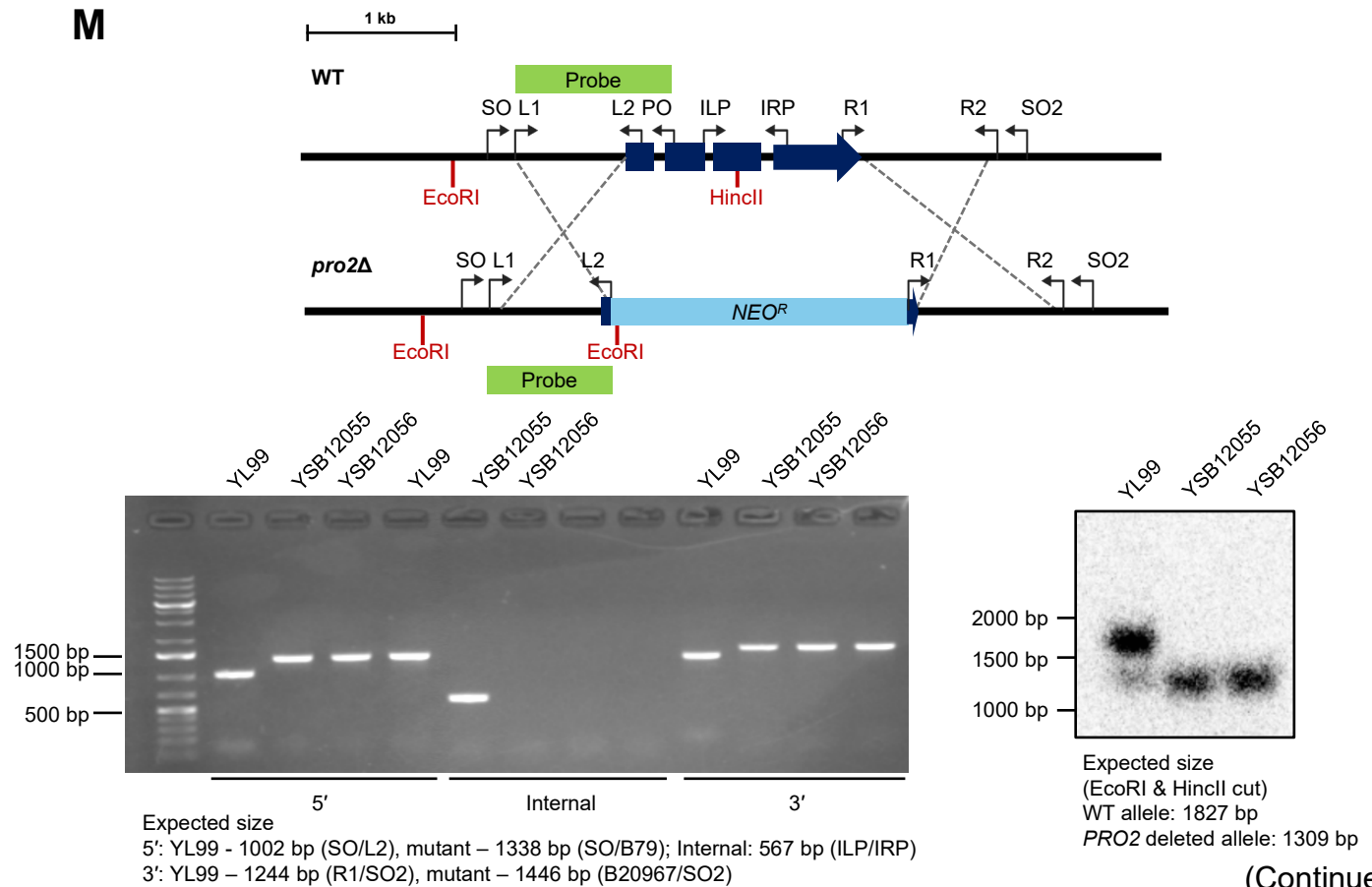

(Continued)

**Figure S2 (Lee et al.)**

**N**

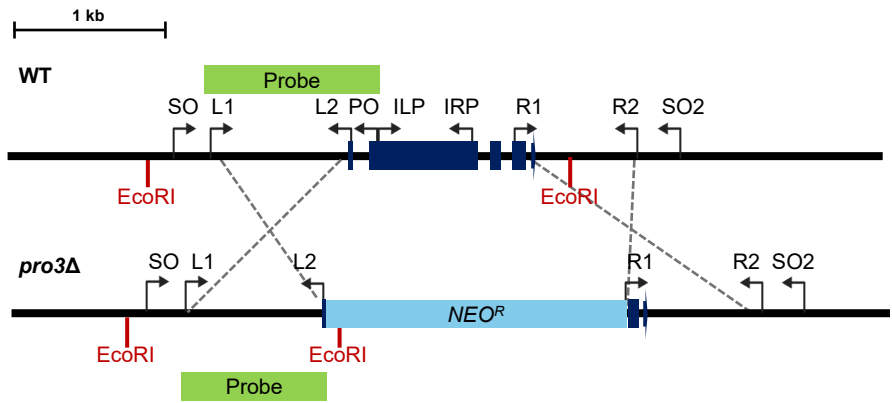

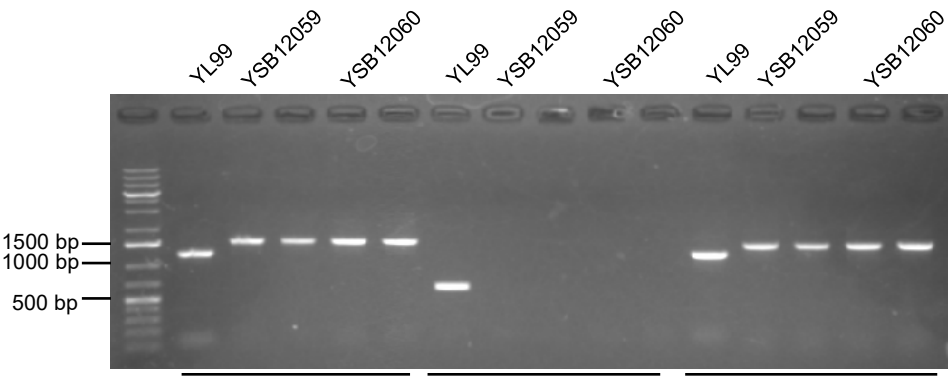

Expected size
5': YL99 - 1203 bp (SO/L2), mutant - 1539 bp (SO/B79); Internal: 624 bp (ILP/IRP)
3': YL99 - 1186 bp (R1/SO2), mutant - 1388 bp (B20967/SO2)

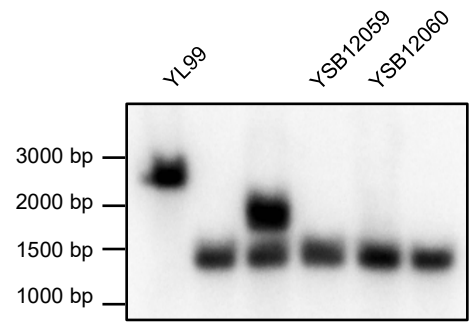

Expected size
(EcoRI cut)
WT allele: 2792 bp
PRO3 deleted allele: 1429 bp

(Continued)

Figure S2 (Lee et al.)

O

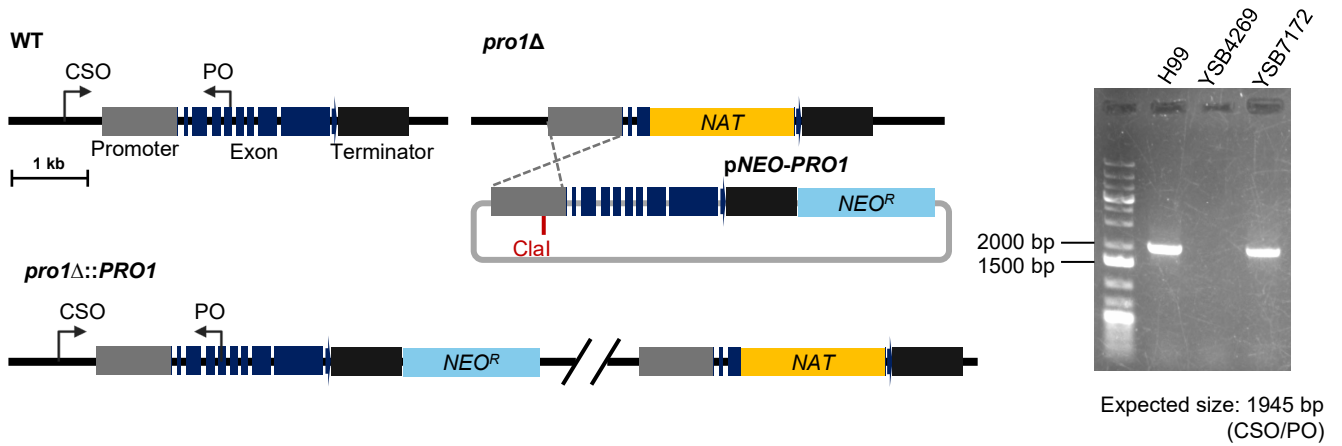

P

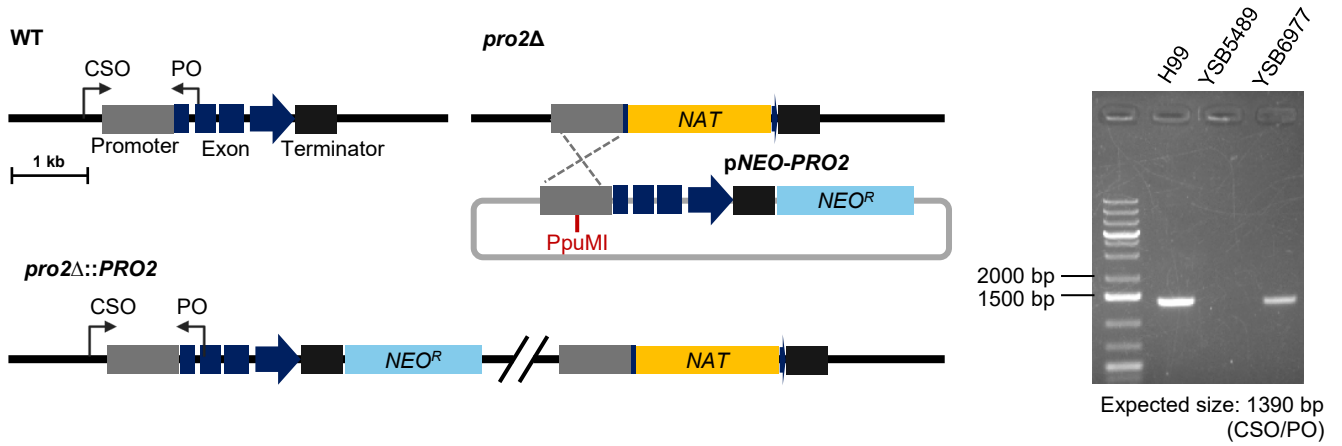

Q

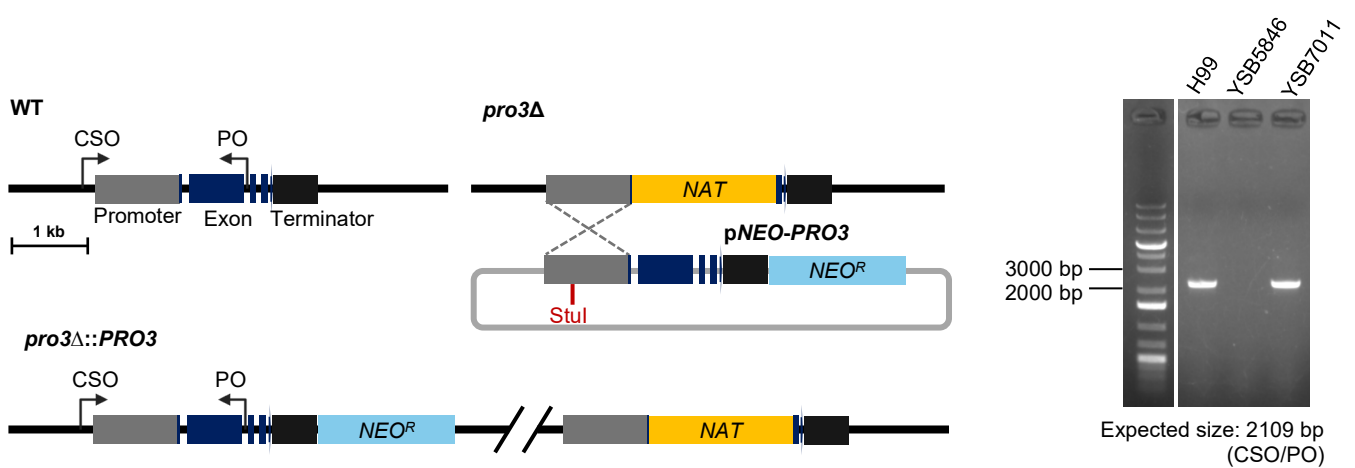

(Continued)

#### Figure S2 (Lee et al.)

**Figure S2. Construction and verification of gene deletion mutants and complemented strains.** (A-K) Generation and genotypic characterization of *MATα* proline metabolism-related mutants. The upper-left panels depict the homologous recombination strategies for constructing *pro2*Δ, *pro3*Δ, *car1*Δ, *pro2*Δ *car1*Δ, *car2*Δ, *pro2*Δ *car2*Δ, *pro3*Δ *pro2*Δ, *pro3*Δ *car2*Δ, *pro3*Δ *pro2*Δ *car2*Δ, *aap6*Δ, and *pro3*Δ *aap6*Δ. In these strategies, the target gene open reading frames (ORFs) were replaced by *NAT*, *NEO<sup>R</sup>*, or *HYG<sup>R</sup>* selection markers. Successful 5' and 3' flanking recombination, as well as internal deletion of the gene, were confirmed by diagnostic PCR (lower-left panels). The precise integration of the disruption cassette was further validated by Southern blot analysis (lower-right panels), showing the expected band sizes for the WT and mutant alleles. (L-N) Construction and verification of *MATα* proline biosynthetic mutants. The upper panels depict the homologous recombination strategies for constructing *pro1*Δ, *pro2*Δ, *pro3*Δ. The lower-left panels present diagnostic PCR results, and the lower-right panels show Southern blot data. (O-Q) The left panels illustrate the targeted insertion strategies used to reintroduce the *PRO1*, *PRO2*, and *PRO3* genes into their respective mutant backgrounds. The right panels present diagnostic PCR results verifying the site-specific integration of each complementation construct into the native locus.

**Figure S3 (Lee et al.)**

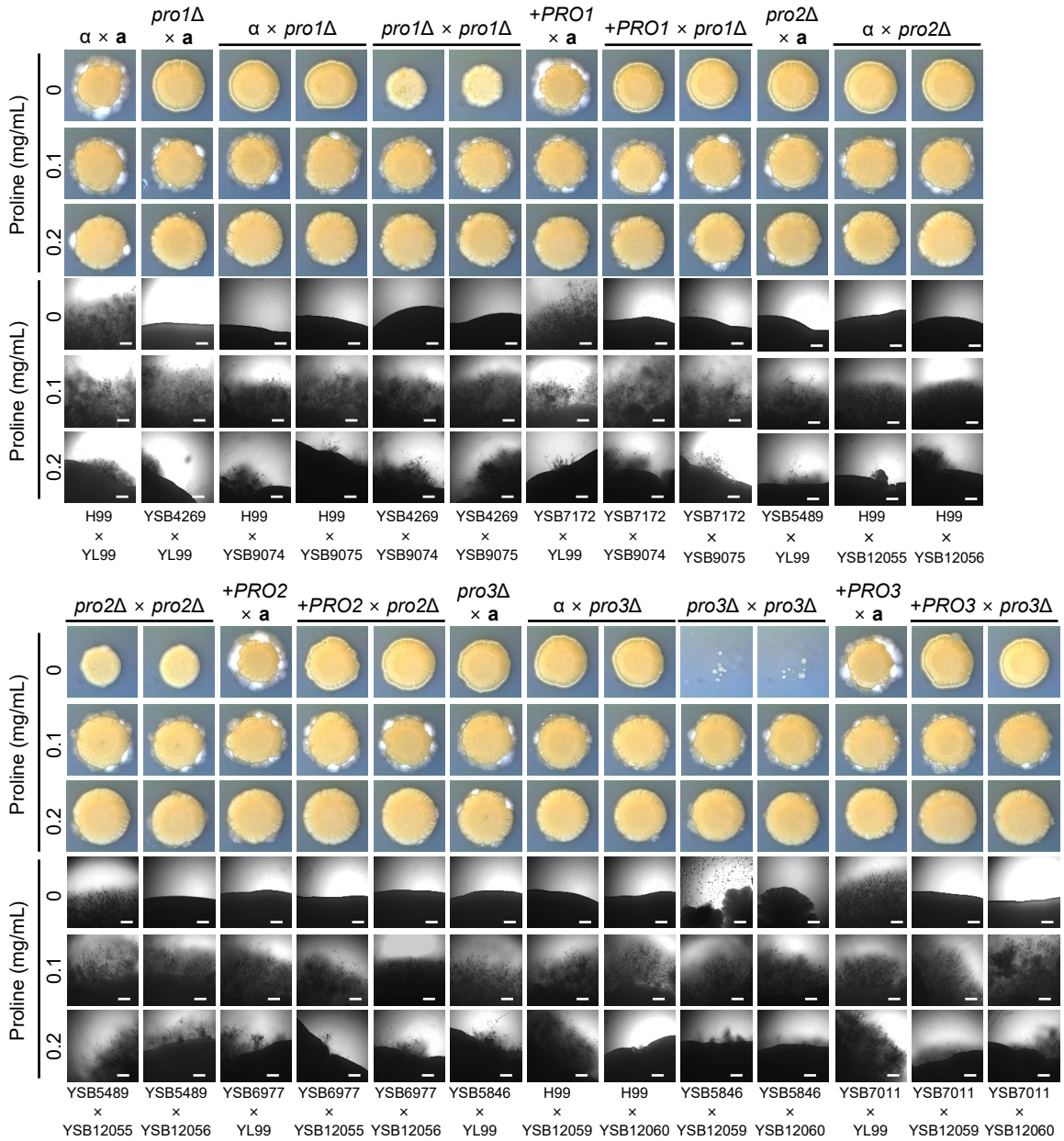

**Figure S3. Comprehensive analysis of sexual development in proline biosynthetic mutants of *Cryptococcus neoformans*.** Mating assays were performed by crossing the indicated strains on V8 agar medium supplemented with varying concentrations of proline (0, 0.1, or 0.2 mg/mL). Plates were incubated at 25°C in the dark for 14 days. The assays include WT crosses (H99 × YL99), unilateral crosses (e.g., $\alpha \times pro1\Delta$, $pro1\Delta \times a$), bilateral crosses (e.g., $pro1\Delta \times pro1\Delta$), and unilateral crosses involving complemented strains (e.g., $+PRO1 \times pro1\Delta$). Representative colony morphologies (top three rows) and filament formation at the colony edges (bottom three rows) are shown. Scale bar, 50 μm. Figure S3 expands the mating analysis shown in Figure 2C by including additional unilateral crosses and proline-supplementation conditions; the WT and bilateral mutant panels are included as reference controls to present the complete experimental matrix.

Figure S4 (Lee et al.)

A

(Continued)

**Figure S4 (Lee et al.)**

**B**

**C**

(Continued)

**Figure S4 (Lee et al.)**

**Figure S4. Additional stress sensitivity assays of proline biosynthetic mutants under nutrient-limited and nutrient-rich conditions.** (A) Stress resistance assay in nutrient-limited YNB media. WT (H99), *pro1Δ*, *pro2Δ*, *pro3Δ*, and their complemented strains were grown overnight in YPD, serially diluted (10-fold), and spotted onto YNB media containing ammonium sulfate (AS) with or without proline (0.2 mg/mL). Stress conditions tested include: 4 mM DIA, 0.03 mM MD, 0.8% CR, 0.8 mg/mL caffeine, 0.01% SDS, 4 mM dithiothreitol (DTT), 0.3 μg/mL TM, 13 μg/mL FLC, 3 μg/mL FDX, 0.02% MMS, 1.5 M NaCl, 1.5 M KCl, 2 M sorbitol, and 35 μM CdSO<sub>4</sub>. Plates were incubated at 30°C for 2 days and imaged. (B) Stress resistance assay in YPD media. Strains were grown overnight in YPD, serially diluted (10-fold), and spotted onto YPD agar plates with or without proline (0.2 mg/mL). Media contained 0.7 mM *tert*-butyl hydroperoxide (tBOOH), 18 mM dithiothreitol (DTT), 10 mg/mL Calcofluor White (CFW), or 0.03% sodium dodecyl sulfate (SDS). Plates were incubated at 30°C for 2 or 3 days and imaged. (C) Stress resistance assay in nutrient-rich YPD media. Resistance was evaluated under various stress conditions: 2.5 mM diamide (DIA), 0.03 mM menadione (MD), 3.5 mM hydrogen peroxide (H<sub>2</sub>O<sub>2</sub>), 1% Congo red (CR), 1.2 mg/mL caffeine, 0.3 μg/mL tunicamycin (TM), 10 μg/mL fluconazole (FLC), 500 μg/mL 5-fluorocytosine (5-FC), 1 μg/mL fludioxonil (FDX), 130 mM hydroxyurea (HU), 0.03% methyl methanesulfonate (MMS), 1.5 M NaCl, 1.5 M KCl, 2 M sorbitol, and 30 μM CdSO<sub>4</sub>. Plates were incubated at 30°C for 2 days and imaged. (D) Thermotolerance and CO<sub>2</sub> tolerance assay. Strains were serially diluted (10-fold) and spotted onto YPD agar plates. Plates were incubated at 30°C, 37°C, 38°C, or 37°C under 5% CO<sub>2</sub> conditions for 2 days before imaging.

**Figure S5 (Lee et al.)**

**Figure S5. *Drosophila melanogaster* survival assay.** Survival of *D. melanogaster* was monitored for 5 days following systemic infection with *C. neoformans* wild-type (H99), proline auxotrophic mutants (*pro1*Δ, *pro2*Δ, and *pro3*Δ) and their respective complemented strains. Fungal cells were grown in YPD at 30 °C, washed, and adjusted to $4 \times 10^8$ cells/mL prior to needle injection. PBS was injected as a negative control. Survival curves were analyzed using the Kaplan–Meier method, and statistical significance was determined by the log-rank (Mantel–Cox) test. Approximately 60 flies were used per experimental group.

**Figure S6 (Lee et al.)**

**Figure S6. Cell size distribution of proline biosynthetic mutants.** Cells from an overnight culture were sub-cultured into fresh YPD media at an initial $OD_{600}$ of 0.2. Once the culture reached the mid-exponential phase ($OD_{600} = 0.8$), this point was defined as 0 h and the cells were further incubated for 72 h with imaging performed at 24 h intervals. Violin plots show the distribution of cell diameters ($n = 100$ cells per strain per time point). Scale bar represents 10 μm. Statistical significance was determined by two-way ANOVA followed by Tukey's multiple-comparison test. The WT data shown here are from the same experimental dataset as those presented in Fig. 5A and are included for comparison.

Figure S7 (Lee et al.)

**Figure S7 (Lee et al.)**

**Figure S7. Mitochondrial staining of proline biosynthetic mutants under basal, elevated-temperature, and host-mimicking conditions.** WT (H99), *pro1Δ*, *pro2Δ*, and *pro3Δ* strains were cultured overnight in YPD at 30°C, inoculated into fresh YPD at an initial OD<sub>600</sub> of 0.2, and grown to an OD<sub>600</sub> of 0.8. Cells harvested at this point were used for the basal condition. For the 39°C and host-mimicking conditions (HMC), cells were washed three times with PBS and incubated with shaking for 3 h in YPD at 39°C or in RPMI 1640 supplemented with 10% FBS at 37°C under 5% CO<sub>2</sub>, respectively. Cells were stained separately with (A) MitoSOX Red, (B) MitoTracker Green FM, or (C) nonyl acridine orange (NAO). Representative DIC and fluorescence images are shown alongside violin plots displaying the distribution of mean fluorescence intensity measured within individual cell outlines using ImageJ (n = 50 per strain per condition). Fluorescence intensity is expressed in arbitrary units (a.u.). Scale bar, 10 μm. Data for each dye were analyzed separately using two-way ANOVA with genotype and culture condition as factors, followed by Tukey's multiple-comparison test to compare strains within each culture condition.

**Figure S8 (Lee et al.)**

**Figure S8. P5C does not significantly accumulate in the *pro3Δ* mutant.** Intracellular P5C quantification in WT, *pro3Δ*, and *pro3Δ::PRO3* strains grown in YPD at 30°C or 37°C for 72 h. P5C was extracted in trichloroacetic acid (TCA), derivatized with 2-aminobenzaldehyde (2-ABZ), and measured by absorbance at 444 nm. Absorbance was corrected against a TCA-buffer-only blank, and expressed as fold-change relative to WT. Data are mean ± SEM (n = 3 independent experiments), with individual data points shown. Statistical significance was determined by one-way ANOVA followed by Tukey's multiple-comparison test separately for each temperature.

**Figure S9 (Lee et al.)**

**Figure S9. Transcriptional responses of putative amino acid permease genes to amino acid supplementation.** Cells were grown overnight in YPD at 30°C and subcultured in fresh YPD from an initial OD<sub>600</sub> of 0.2 until reaching an OD<sub>600</sub> of 0.8, before transfer to the indicated YNB conditions. Transcript levels were quantified by qRT-PCR, normalized to *ACT1*, and calculated using the $2^{-\Delta\Delta C_t}$ method. Data are presented as mean ± SEM from three biological replicates, with individual data points shown. Technical replicates were averaged before statistical analysis. (A) *AAP6* transcript levels in WT (H99) cells incubated in YNB without ammonium sulfate (AS), with or without arginine (ARG, 0.5 mg/mL) for 15 or 30 min. The YPD control sampled immediately before the nutrient shift (time zero) was used as the calibrator and assigned a value of 1. Statistical comparisons were performed using unpaired two-tailed t-tests with Welch's correction between the conditions with and without arginine separately at each time point. (B) Transcript levels of *AAP1*–*AAP8*, *MUP1*, and *MUP3* in WT and *aap6*Δ cells incubated in YNB without AS, with or without proline (0.2 mg/mL) for 15 min. For each gene, the mean $\Delta C_t$ of the WT YPD control sampled immediately before the nutrient shift (time zero) was used as the common calibrator for all four groups. Statistical significance was assessed using one-way ANOVA followed by Tukey's multiple-comparison test.

**Table S1. Strains used in this study**

| Strain | Genotypes | Parent | References |
| --- | --- | --- | --- |
| H99 | *MATα* |  | (1, 2) |
| YL99 | *MATα* |  | (3) |
| YSB4269 | *MATα pro1Δ::NAT STM#5* | H99 | (4) |
| YSB7172 | *MATα pro1Δ::PRO1-NEO* | YSB4269 | In this study |
| YSB5489 | *MATα pro2Δ::NAT STM#58* | H99 | In this study |
| YSB5490 | *MATα pro2Δ::NAT STM#58* | H99 | In this study |
| YSB5491 | *MATα pro2Δ::NAT STM#58* | H99 | In this study |
| YSB6977 | *MATα pro2Δ::PRO2-NEO* | YSB5489 | In this study |
| YSB5494 | *MATα pro3Δ::NAT STM#43* | H99 | In this study |
| YSB5495 | *MATα pro3Δ::NAT STM#43* | H99 | In this study |
| YSB5846 | *MATα pro3Δ::NAT STM#43* | H99 | In this study |
| YSB5847 | *MATα pro3Δ::NAT STM#43* | H99 | In this study |
| YSB5848 | *MATα pro3Δ::NAT STM#43* | H99 | In this study |
| YSB7011 | *MATα pro3Δ::PRO3-NEO* | YSB5846 | In this study |
| YSB9074 | *MATα pro1Δ::NEO* | YL99 | In this study |
| YSB9075 | *MATα pro1Δ::NEO* | YL99 | In this study |
| YSB12055 | *MATα pro2Δ::NEO* | YL99 | In this study |
| YSB12056 | *MATα pro2Δ::NEO* | YL99 | In this study |
| YSB12059 | *MATα pro3Δ::NEO* | YL99 | In this study |
| YSB12060 | *MATα pro3Δ::NEO* | YL99 | In this study |
| YSB12288 | *MATα car1Δ::NAT STM#102* | H99 | In this study |
| YSB12289 | *MATα car1Δ::NAT STM#102* | H99 | In this study |
| YSB12290 | *MATα car1Δ::NAT STM#102* | H99 | In this study |
| YSB12291 | *MATα car1Δ::NAT STM#102* | H99 | In this study |
| YSB12292 | *MATα car1Δ::NAT STM#102* | H99 | In this study |
| YSB12293 | *MATα car1Δ::NAT STM#102* | H99 | In this study |
| YSB12259 | *MATα pro2Δ::NAT car1Δ::NEO* | YSB5489 | In this study |
| YSB12260 | *MATα pro2Δ::NAT car1Δ::NEO* | YSB5489 | In this study |
| YSB12262 | *MATα pro2Δ::NAT car1Δ::NEO* | YSB5489 | In this study |
| YSB12263 | *MATα pro2Δ::NAT car1Δ::NEO* | YSB5489 | In this study |
| YSB12294 | *MATα car2Δ::NAT STM#116* | H99 | In this study |
| YSB12295 | *MATα car2Δ::NAT STM#116* | H99 | In this study |
| YSB12264 | *MATα pro2Δ::NAT car2Δ::NEO* | YSB5489 | In this study |
| YSB12265 | *MATα pro2Δ::NAT car2Δ::NEO* | YSB5489 | In this study |
| YSB12267 | *MATα pro2Δ::NAT car2Δ::NEO* | YSB5489 | In this study |
| YSB12296 | *MATα aap6Δ::NAT STM#119* | H99 | In this study |
| YSB12297 | *MATa aap6Δ::NAT STM#119* | H99 | In this study |
| YSB12298 | *MATa aap6Δ::NAT STM#119* | H99 | In this study |
| YSB12273 | *MATa pro3Δ::NAT aap6Δ::NEO* | YSB5846 | In this study |
| YSB12274 | *MATa pro3Δ::NAT aap6Δ::NEO* | YSB5846 | In this study |
| YSB12275 | *MATa pro3Δ::NAT aap6Δ::NEO* | YSB5846 | In this study |
| YSB12269 | *MATa pro3Δ::NAT pro2Δ::NEO* | YSB5846 | In this study |
| YSB12270 | *MATa pro3Δ::NAT pro2Δ::NEO* | YSB5846 | In this study |
| YSB12271 | *MATa pro3Δ::NAT pro2Δ::NEO* | YSB5846 | In this study |
| YSB12272 | *MATa pro3Δ::NAT pro2Δ::NEO* | YSB5846 | In this study |
| YSB12338 | *MATa pro3Δ::NAT car2Δ::NEO* | YSB5846 | In this study |
| YSB12339 | *MATa pro3Δ::NAT car2Δ::NEO* | YSB5846 | In this study |
| YSB12340 | *MATa pro3Δ::NAT car2Δ::NEO* | YSB5846 | In this study |
| YSB12341 | *MATa pro3Δ::NAT car2Δ::NEO* | YSB5846 | In this study |
| YSB12342 | *MATa pro3Δ::NAT car2Δ::NEO* | YSB5846 | In this study |
| YSB12343 | *MATa pro3Δ::NAT pro2Δ::NEO car2Δ::HYG* | YSB12269 | In this study |
| YSB12344 | *MATa pro3Δ::NAT pro2Δ::NEO car2Δ::HYG* | YSB12269 | In this study |
| YSB12345 | *MATa pro3Δ::NAT pro2Δ::NEO car2Δ::HYG* | YSB12269 | In this study |
| YSB12346 | *MATa pro3Δ::NAT pro2Δ::NEO car2Δ::HYG* | YSB12269 | In this study |

**Table S2. Primers used in this study**

| Primer | Description | Sequence (5' to 3') |
| --- | --- | --- |
| B1846 | M13 FWD for NAT marker | GTAAAACGACGGCCAGTGAGC |
| B1847 | M13 REV for NAT marker | CAGGAAACAGCTATGACCATG |
| B1026 | M13 FWD for NEO marker | GTAAAACGACGGCCAGTGAAT |
| B1027 | M13 REV for NEO marker | CAGGAAACAGCTATGACCATG |
| B1454 | *NAT* split marker FWD | AAGGTGTTCCCCGACGACGAATCG |
| B1455 | *NAT* split marker REV | AACTCCGTCGCGAGCCCCATCAAC |
| B1886 | NEO split marker FWD | TGGAAGAGATGGATGTGC |
| B1887 | NEO split marker REV | ATTGTCTGTTGTGCCCAG |
| B79 | *NAT*/NEO marker 5' screening | TGTGGATGCTGGCGGAGGATA |
| B20697 | *NAT* marker 3' screening | CGAATCCGAGACAGACAT |
| B20967 | NEO marker 3' screening | CGAGACAGACATCGTGTC |
| B22496 | *PRO1* deletion L1 | CAATGGTTCTACTCTCTCCAGTC |
| B22554 | *PRO1* deletion L2 | TCACTGGCCGTCGTTTTACCCTTCCCTCTTTCCTTGAG |
| B23220 | *PRO1* deletion R1 | CATGGTCATAGCTGTTTCCTGGAAAGAGGTGAGCAACTATCG |
| B23409 | *PRO1* deletion R2 | GGGCTCTGAATCATCTTGG |
| B22553 | *PRO1* screening SO | GATACCATTTGCCGTTGC |
| B14291 | *PRO1* internal screening ILP | CTCCTTCCTGCTGGGGTAGT |
| B14292 | *PRO1* internal screening IRP | TTTCGTCCCTTCATCCTGTC |
| B14290 | *PRO1* 3' screening SO2 | GAGCAGAAGGGACAAGATGC |
| B22493 | *PRO1* probe PO | ACGATAGGAATAACGCCC |
| B22499 | *PRO1* CSO | GCTCAGGCGTTACGATTCTG |
| B10666 | *PRO1* CLP | GGAGGAAGAAAGTCCAGTATTG |
| B10667 | *PRO1* CRP | GAAGAGCAGAAGGGACAAG |
| B22481 | *PRO2* deletion L1 | GCCGAAAACATTTGAGTTCACAGG |
| B22482 | *PRO2* deletion L2 | TCACTGGCCGTCGTTTTACCAGCGATGGCGATAGATTC |
| B22483 | *PRO2* deletion R1 | CATGGTCATAGCTGTTTCCTGTGGAAGTGTGATTGCCGACC |
| B22484 | *PRO2* deletion R2 | TGCTGTAAGTCTGTTCGC |
| B22485 | *PRO2* screening SO | TTAGCGACCACCCATCCAAGTAGG |
| B22555 | *PRO2* internal screening ILP | CTTTGGCCATCAAAAGTGGT |
| B22556 | *PRO2* internal screening IRP | GATCGCATCGAAGACAGACA |
| B23066 | *PRO2* 3' screening SO2 | AGGCGATGGAGATGAACTTG |
| B22573 | *PRO2* probe PO | GCCTCAGCAGCCTATTTCTG |
| B10582 | *PRO2* CSO | AGTGGTTACAAAGGGTTCG |
| B10332 | *PRO2* CLP | ATAACCGTGTTTAGCGACC |
| B10333 | *PRO2* CRP | CTTTCGCCCTCCAAATAC |
| B22487 | *PRO3* deletion L1 | TTCGCCAGTATGTTCACAC |
| B22488 | *PRO3* deletion L2 | TCACTGGCCGTCGTTTTACAGCGAGAGTGTAGCCCATTG |
| B22489 | *PRO3* deletion R1 | CATGGTCATAGCTGTTTCCTGTGTTGACACTTGAAGACGG |
| B22490 | *PRO3* deletion R2 | CGAGATGAGGGTGATGTGAG |
| B22491 | *PRO3* screening SO | GTCGCTCCTATCAAATGTCTC |
| B22557 | *PRO3* internal screening ILP | TCTCTTCCTCGCAACCCTAA |
| B22558 | *PRO3* internal screening IRP | CAAACCCATCATCACACCAC |
| B23067 | *PRO3* 3' screening SO2 | CGGTGGAGATGAGGATCTGT |
| B22572 | *PRO3* probe PO_2 | TTAGGGTTGCGAGGAAGAGA |
| B22492 | *PRO3* complementation PO | CAGTTCACACAAGACTACTACC |
| B10583 | *PRO3* CSO | TTCGGTATTGGTCGTATCC |
| B10334 | *PRO3* CLP | TCCACGAAGTTTGTGAGTG |
| B10335 | *PRO3* CRP | CGTCCGAATCTTATCCAGA |
| B23362 | *CAR1* deletion L1 | TCCCTCGCATACATCATTCA |
| B23363 | *CAR1* deletion L2 | TCACTGGCCGTCGTTTTACAGAGATGGAGAAACGCGAGA |
| B23364 | *CAR1* deletion R1 | CATGGTCATAGCTGTTTCCTGGCGTCTTTGGGTGAGACTTT |
| B23365 | *CAR1* deletion R2 | CCTCAACCAAATCCTCCTGA |
| B23366 | *CAR1* screening SO | CACTCGTGAATGGATGATGG |
| B23368 | *CAR1* internal screening ILP | GGATGGTTGCCTGTGACTTT |
| B23369 | *CAR1* internal screening IRP | CAACGTCGAAACTGAGGTGA |
| B23370 | *CAR1* 3' screening SO2 | CCTGACCATCTTCGTCCAGT |
| B23367 | *CAR1* probe PO | GGCATCCGACAAGCTATCAT |
| B23373 | *CAR2* deletion L1 | CATAAGGCATTGGGCAAAGT |
| B23374 | *CAR2* deletion L2 | TCACTGGCCGTCGTTTTACTTGGGGACATTGCTTGGTAT |
| B23375 | *CAR2* deletion R1 | CATGGTCATAGCTGTTTCCTGGAGGAGCACAAGGCTGCTAT |
| B23376 | *CAR2* deletion R2 | CAGCCTGAAAATGGGTTACG |
| B23377 | *CAR2* screening SO | TCTCGAGGTGTTTGGCTTCT |
| B23379 | *CAR2* internal screening ILP | TCGCTCGTAAATGGGGTTAC |
| B23380 | *CAR2* internal screening IRP | CTCCAGGCTTAATGCAAAGC |
| B23381 | *CAR2* 3' screening SO2 | CCTCTCAACCGTCTTCAAGC |
| B23378 | *CAR2* probe PO | TGTGACTGCAAAACGTGTCA |
| B23643 | *AAP6* deletion L1 | TGTGGACGAATCGAATATGG |
| B23644 | *AAP6* deletion L2 | TCACTGGCCGTCGTTTTACCGGTCACCTTCGGATCATAG |
| B23645 | *AAP6* deletion R1 | CATGGTCATAGCTGTTTCCTGGATCATGGCCAAGCTGTTTT |
| B23646 | *AAP6* deletion R2 | AAGGTATGTCGCGCTGTTGT |
| B23647 | *AAP6* screening SO | TTGGAGGCCAAGGAACATAC |
| B23715 | *AAP6* internal screening ILP | TCGGTGAAGCTGAATGTGAG |
| B23716 | *AAP6* internal screening IRP | TTGGGTCATTGTAAGGCACA |
| B23651 | *AAP6* 3' screening SO2 | GGTTACAAATTCCCGGTCCT |
| B23648 | *AAP6* probe PO | CCGTCGAGTGGAAATAAGGA |
| B679 | *ACT1* qLP | CGCCCTTGCTCCTTCTTCTATG |
| B680 | *ACT1* qRP | GACTCGTCGTATTCGCTCTTCG |
| B23371 | *CAR1* qLP | GACATTGACGACGCTGAGAA |
| B23372 | *CAR1* qRP | TACTGAAGCGCGAGTTCCAT |
| B23382 | *CAR2* qLP | CAACGGTGTGGTTATTGACG |
| B23383 | *CAR2* qRP | GAGTAGGCTTGGCCAACAAG |
| B15938 | *LAC1* qLP | CACCCTTTGGAAGTTGTGG |
| B15939 | *LAC1* qRP | TGATAATTGCAGAGTACCG |
| B23560 | *AAP1* (CNAG_02539) qLP | GCCTTATGGTATACTCTATGATG |
| B23561 | *AAP1* (CNAG_02539) qRP | CCGTATGCCCGAGCACCGAGG |
| B23562 | *AAP2* (CNAG_07902) qLP | CATGGTATACGCGATGATGG |
| B23563 | *AAP2* (CNAG_07902) qRP | TCTGGCTCCCCAGAAGTTAATG |
| B23564 | *AAP3* (CNAG_01118) qLP | GTGGGGTTAATCATCCTTGG |
| B23565 | *AAP3* (CNAG_01118) qRP | ATGTACCACCGAGATAAAAG |
| B23566 | *AAP4* (CNAG_00597) qLP | CGAGGCAAAGAACCCACG |
| B23567 | *AAP4* (CNAG_00597) qRP | AATCAAGATGCAAGCGTTTATG |
| B23568 | *AAP5* (CNAG_07367) qLP | ACTTACTTGGACCTCTATCCTC |
| B23569 | *AAP5* (CNAG_07367) qRP | TTTTTCGGATCAGCTTGAAACC |
| B23570 | *AAP6* (CNAG_07449) qLP | CCTTGAAAGACCGTTTCGGC |
| B23571 | *AAP6* (CNAG_07449) qRP | TGTCACAAGTGTTGGGTCATTG |
| B23572 | *AAP7* (CNAG_05345) qLP | CACTGGAAATGCTGCTGTGT |
| B23573 | *AAP7* (CNAG_05345) qRP | CATGTATGTGAAAGCGATGG |
| B23574 | *AAP8* (CNAG_00574) qLP | TCTCTTTCTAGGGATTCTTATC |
| B23575 | *AAP8* (CNAG_00574) qRP | CTCCGCCATATGGGCAGAAGC |
| B23576 | *MUP1* (CNAG_07693) qLP | GTTCAACTTCATTGTTGCCC |
| B23577 | *MUP1* (CNAG_07693) qRP | AGGAACACGATCCCTAGTC |
| B23578 | *MUP3* (CNAG_03955) qLP | TATGTGGCAGGCGAAATGC |
| B23579 | *MUP3* (CNAG_03955) qRP | CCTTGCAGTTGTGTACAATC |

#### REFERENCES

1. Perfect JR, Ketabchi N, Cox GM, Ingram CW, Beiser CL. 1993. Karyotyping of *Cryptococcus neoformans* as an epidemiological tool. J Clin Microbiol 31:3305–3309. doi:10.1128/jcm.31.12.3305-3309.1993.
2. Arras SDM, Ormerod KL, Erpf PE, Espinosa MI, Carpenter AC, Blundell RD, Stowasser SR, Schulz BL, Tanurdzic M, Fraser JA. 2017. Convergent microevolution of *Cryptococcus neoformans* hypervirulence in the laboratory and the clinic. Sci Rep 7:17918. doi:10.1038/s41598-017-18106-2.
3. Semighini CP, Averette AF, Perfect JR, Heitman J. 2011. Deletion of *Cryptococcus neoformans* AIF ortholog promotes chromosome aneuploidy and fluconazole-resistance in a metacaspase-independent manner. PLoS Pathog 7:e1002364. doi:10.1371/journal.ppat.1002364.
4. Lee KT, So YS, Yang DH, Jung KW, Choi J, Lee DG, Kwon H, Jang J, Wang LL, Cha S, Meyers GL, Jeong E, Jin JH, Lee Y, Hong J, Bang S, Ji JH, Park G, Byun HJ, Park SW, Park YM, Adedoyin G, Kim T, Averette AF, Choi JS, Heitman J, Cheong E, Lee YH, Bahn YS. 2016. Systematic functional analysis of kinases in the fungal pathogen *Cryptococcus neoformans*. Nat Commun 7:12766. doi:10.1038/ncomms12766.
